# Floating search methodology for combining classification models for site recognition in DNA sequences

**DOI:** 10.1101/320309

**Authors:** Javier Pérez-Rodríguez, Aida de Haro-García, Nicolás García-Pedrajas

**Affiliations:** Department of Computing and Numerical Analysis, University of Córdoba, Spain

**Author notes:** All of the authors contributed equally to this work.

## Abstract

Recognition of the functional sites of genes, such as translation initiation sites, donor and acceptor splice sites and stop codons, is a relevant part of many current problems in bioinformatics. Recognition of the functional sites of genes is also a fundamental step in gene structure predictions in the most powerful programs. The best approaches to this type of recognition use sophisticated classifiers, such as support vector machines. However, with the rapid accumulation of sequence data, methods for combining many sources of evidence are necessary as it is unlikely that a single classifier can solve this type of problem with the best possible performance.

A major issue is that the number of possible models to combine is large and the use of all of these models is impractical. In this paper, we present a framework that is based on floating search for combining as many classifiers as needed for the recognition of any functional sites of a gene. The methodology can be used for the recognition of translation initiation sites, donor and acceptor splice sites and stop codons. Furthermore, we can combine any number of classifiers that are trained on any species. The method is also scalable to large datasets, as is shown in experiments in which the whole human genome is used. The method is also applicable to other recognition tasks.

We present experiments on the recognition of these four functional sites in the human genome, which is used as the target genome, and use another 20 species as sources of evidence. The proposed methodology shows significant improvement over state-of-the-art methods for use in a thorough evaluation process. The proposed method is also able to improve heuristic selection of species to be used as sources of evidence as the search finds the most useful datasets.

**Author summary:** In this paper we present a methodology for combining many sources of information to recognize some of the most important functional sites in a genomic sequence. The functional sites of the sequences, such as, translation start sites, translation initiation sites, acceptor and donor splice sites and stop codons, play a very relevant role in many Bioinformatics tasks. Their accurate recognition is an important task by itself and also as part of gene structure prediction programs.

Our approach uses a methodology usually termed in Computer Science as “floating search”. This is a powerful heuristics applicable when the cost of evaluating each possible solution is high. The methodology is applied to the recognition of four different functional sites in the human genome using as additional sources of evidence the annotated genomes of other twenty different species.

The results show an advantage of the proposed method and also challenge the standard assumption of using only genomes not very close and not very far from the human to improve the recognition of functional sites in the human genome.

## Introduction

The recognition of functional sites within the genome is one of the most important problems in bioinformatics research. Determining where different functional sites, such as promoters, translation start sites, translation initiation sites (TISs), donors, acceptors and stop codons are located provides useful information for many tasks [1]. For instance, the recognition of translation initiation sites, donors, acceptors and stop codons [2] is one of the most critical tasks for gene structure prediction.

Many of the most successful gene recognizers that are currently in use implement an initial step of site recognition [3], which is followed by a process of combining the sites into meaningful gene structures. Accurate recognition is of the utmost importance for the whole gene structure prediction process. Actual sites that are not found by the classification models likely result in exons not being considered by the remaining steps of the recognition program. Furthermore, many false positives might inundate the second step, thereby making it difficult to predict gene structures accurately. State-of-the-art approaches use powerful classifiers, such as support vector machines (SVMs), and consider moderately large sequences around the functional site of interest [2, 4–6].

In recent years, information about the genomes of many species has been accumulated. This information can be used to improve the recognition of functional sites. However, the arbitrary selection of species using the widely assumed hypothesis that we must consider moderately distant evolutionary relatives is clearly a suboptimal procedure [7]. In addition, the classifier models are chosen a priori, without considering the possible benefits of combining various models.

It would be more efficient to learn all of the available classification models and obtain the best combination using an automatic method. The problem of finding the best combination can be tackled as a search problem over all possible combinations. An exhaustive search is unfeasible even for a small number of models. Other common search heuristics, such as evolutionary computation and swarm intelligence, are also prohibitively costly in terms of running time.

In cases when those heuristics cannot be used, floating search is an inexpensive yet sufficiently powerful methodology that is able to achieve very good solutions. Floating search has been used when the cost of each search step is high [8]. Thus, in this work, we propose using floating search to obtain a near-optimal combination of classification models, in which we can consider as many sources of evidence as are available and use as many classifiers as needed using various floating search methods, namely, Sequential Forward Selection, Sequential Backward Selection, Plus-*l* Minus-*r* Selection, Sequential Forward Floating Selection, Sequential Backward Floating Selection, Random Sequential Forward Floating Selection and Random Sequential Backward Floating Selection. Although the first two methods are not actually floating search methods but sequential greedy approaches, we included them for completeness.

To evaluate the proposed method, we show results for the recognition of the four functional sites that are cited above in five chromosomes of the human genome. To demonstrate the ability of our method to combine many classifiers we used for TIS and stop codon recognition 6 models for each of the 21 complete genomes, for a total of 126 classifiers. For donor and acceptor recognition, we used 5 models for the same 21 genomes, for a total of 105 classifiers.

## Materials and methods

As stated in the introduction, our major aim is to develop a combination method for obtaining optimal, or near-optimal, subsets of classification models that are trained for site recognition in DNA sequences. An exhaustive search would require the evaluation of 2*^N^ −* 1 combinations of models given a set of *N* trained classifiers. This type of search is infeasible even for a small value of *N*. Therefore, we must use a search algorithm to find the best possible model combination efficiently. Many powerful metaheuristics are available in the machine learning literature, such as evolutionary computation [9], particle swarm optimization [10], ant colonies [11] and differential evolution [12]. However, all of these methodologies require the repetitive evaluation of many solutions to achieve their optimization goal. In the problem of site recognition, the evaluation of a possible solution is a costly process due to the large datasets that are involved. Thus, these metaheuristics are not feasible.

Instead, we propose a simpler approach, namely, floating search, which has obtained successful results in other research fields, such as feature selection [13–16]. Floating search, which will be described in depth in the following section, is a set of stepwise search methods that are fast and efficient at solving problems in which the evaluation of many possible candidate solutions is too computationally expensive.

The process for obtaining the best combination of classifiers for various species is composed of two steps: a training step and validation step. Before starting the learning process, we need to obtain the training datasets, testing dataset and validation dataset. Without a loss of generality and to provide the necessary focus for our description, we use the same setup as in the experiments that are reported below. We address the problem of site recognition in the human genome. To solve this problem, we use a test set of sites of a specific chromosome, which we denote as *T*. The training set includes all of the remaining human chromosomes and genomes of all of the species we choose to evaluate. For validation, we use one of the human chromosomes in the training set, which we denote as *V* and remove it from the training set.

### Floating search

As stated above, the use of complex heuristics for combining tens or hundreds of models would incur an infeasible computational cost. Thus, we propose the use of simpler, yet still powerful, heuristics. We state our problem as a search problem to enable the application of those heuristics. We have *N* trained classifiers *C* = {*c*_1_, *c*_2_,…, *c_N_*}, which are trained using any types of sequences that could be useful, and use any genome that we consider interesting. Our aim is to obtain a subset of classifiers *C′* ⊂ *C* that is the best possible combination. Evaluation of the combination of models is carried out using cross-validation. Thus, our objective function for maximization is the accuracy of the combination of classifiers over a validation set *V*, which is denoted as *J*(*V*).

Among the simplest methods, Sequential Forward Selection (SFS) [17] (see Algorithm 1) and Sequential Backward Selection (SBS) [18] (see Algorithm 2) are widely used because of their easy implementation and speed. The SFS method starts with an empty set and adds one classifier at a time to the selected subset by choosing the classifier that maximizes *J*(*V*). The method terminates when the value of *J*(*V*) is no longer improving or a desired number of classifiers has been reached. SBS starts from the opposite side by considering all of the classifiers and removing one classifier at a time. For classifier removal, *J*(*V*) is evaluated and the model that maximizes *J*(*V*) is removed. The stop criterion could be a number of classifiers that are removed or a decrease of *J*(*V*) is observed. In our experiments, we removed classifiers while *J*(*V*) does not decrease. These two methods can be generalized to add or remove *r* ≥ 1 classifiers in every iteration. These methods are fast and can obtain good results, but have two major problems: They easily become trapped in local minima and suffer from the “nesting effect” [19]. The nesting effect means that to obtain an optimal solution of size *M*, it must contain the optimal solution of size *M −* 1, which is not often the case in practice.

#### Algorithm 1

Sequential Forward Selection (SFS).

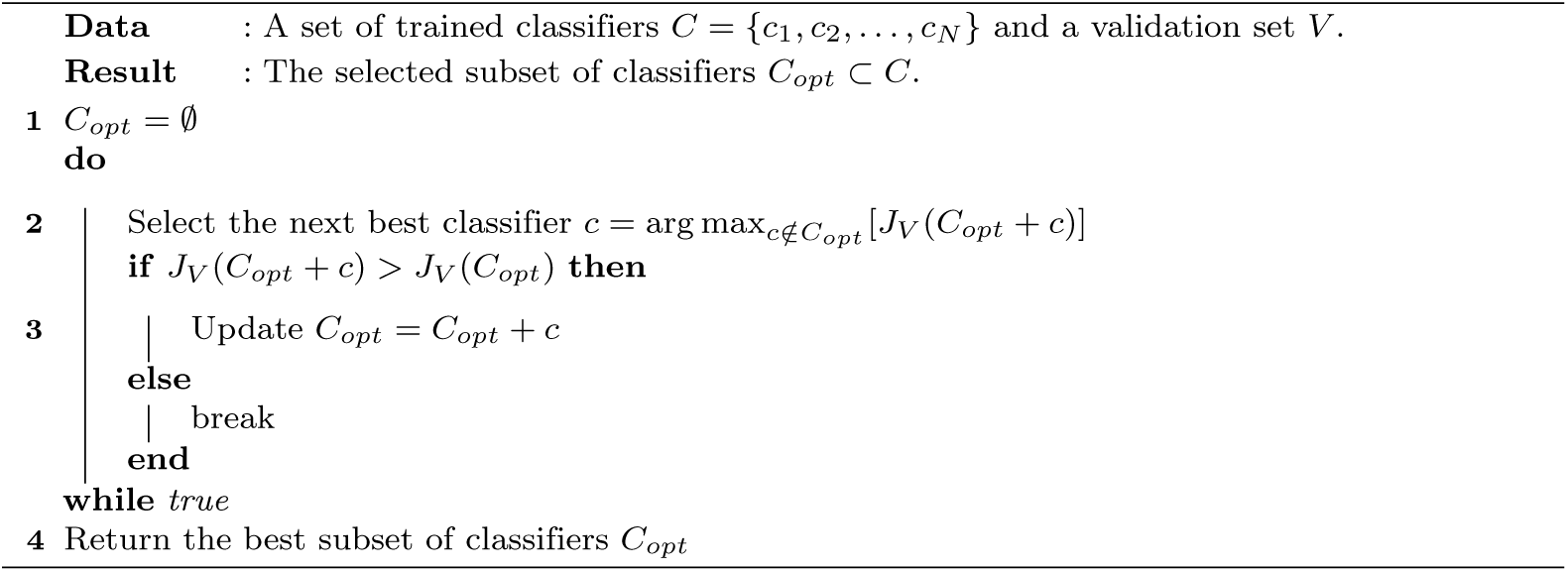

#### Algorithm 2

Sequential Backward Selection (SBS).

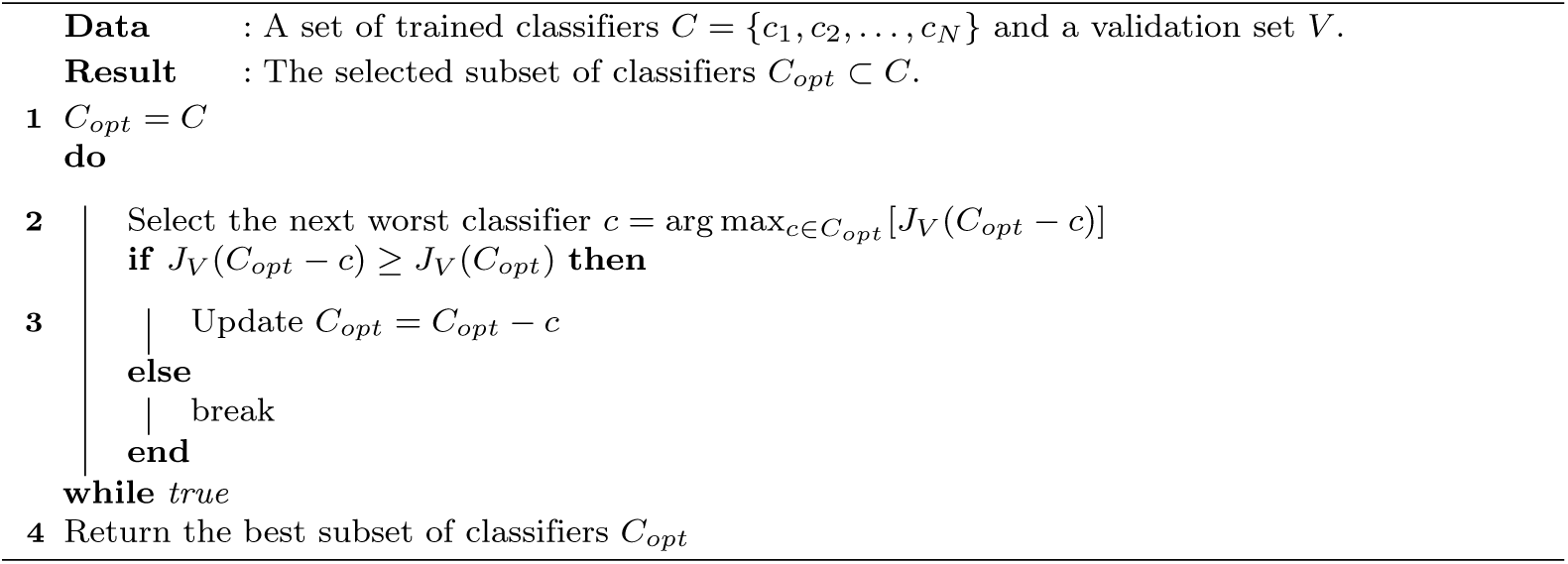

The nesting problem can be avoided using the Plus-*l* Minus-*r* Selection (LRS) search method [20]. LRS adds backtracking capabilities by using SFS to add *l* models and SBS to remove *r* models. However, one major problem is that there is no rule for choosing the best values of *l* and *r*. The LRS method is shown as Algorithm 3.

A more advanced approach is floating search. In floating search, we let the size of the solution “float” and adapt to the problem using a backtracking mechanism. In that way, Sequential Forward Floating Selection (SFFS) and Sequential Backward Floating Selection (SBFS) [8] overcome the nesting problem and the local minimum problem by backtracking after adding (or removing) a new model. SFFS starts with an empty set and proceeds as SFS. However, after adding a new model, SFFS allows any of the previously added models to be removed until the value of *J* worsens. SBFS does the opposite: it follows the SBS method and allows removed models to be added. Algorithms 4 and 5 show the SFFS and SBFS methods, respectively. The comparisons [21] usually demonstrate better performances of SFFS and SBFS compared to SFS and SBS.

Somole et al. [13] proposed an adaptive version for feature selection in which the number of models to add or remove was incremented when the desired number of features was small. However, the method achieved only marginal improvement and required a longer execution time.

#### Algorithm 3

Plus-*l* Minus-*r* Selection (LRS).

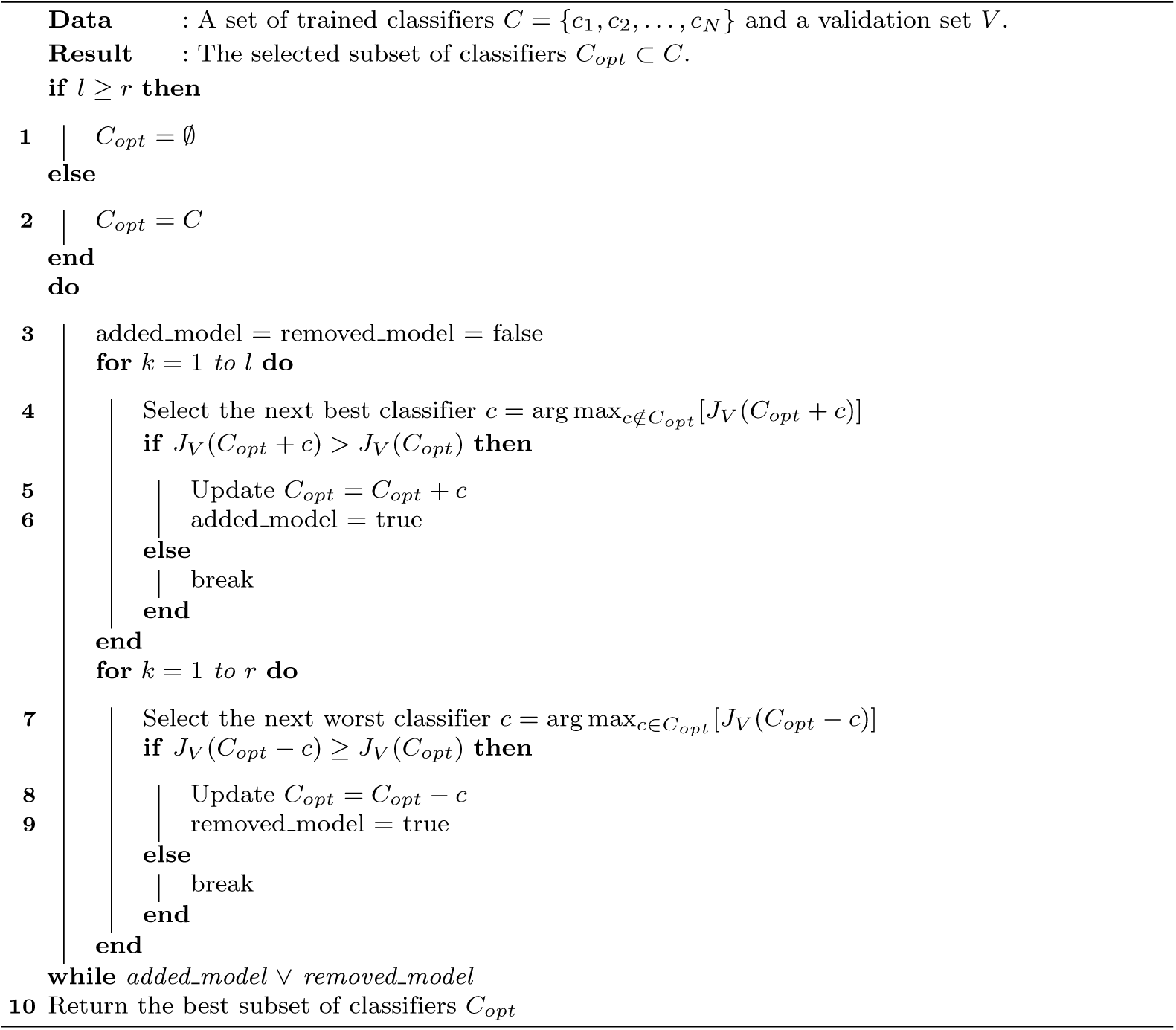

#### Algorithm 4

Sequential Forward Floating Selection (SFFS).

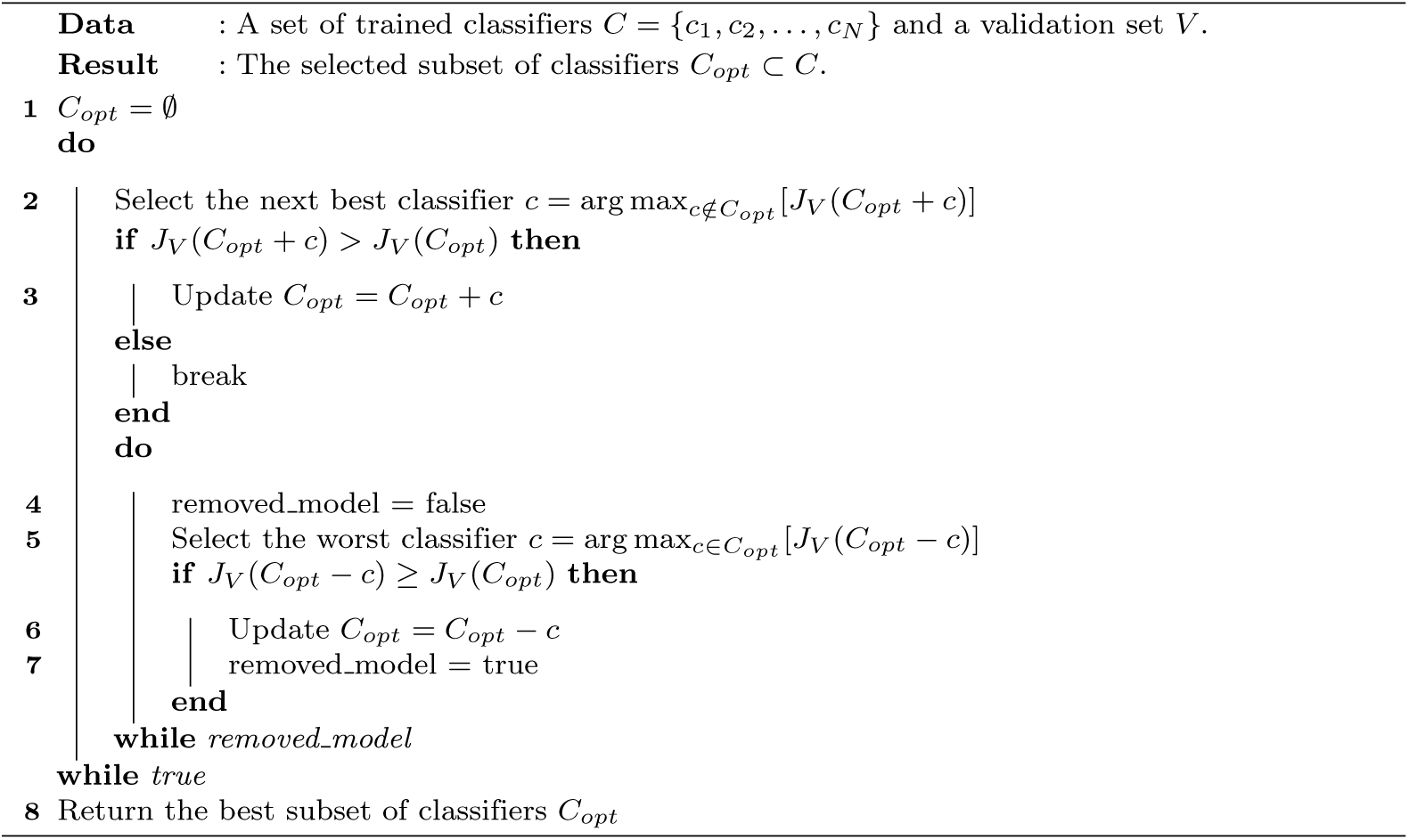

#### Algorithm 5

Sequential Backward Floating Selection (SBFS).

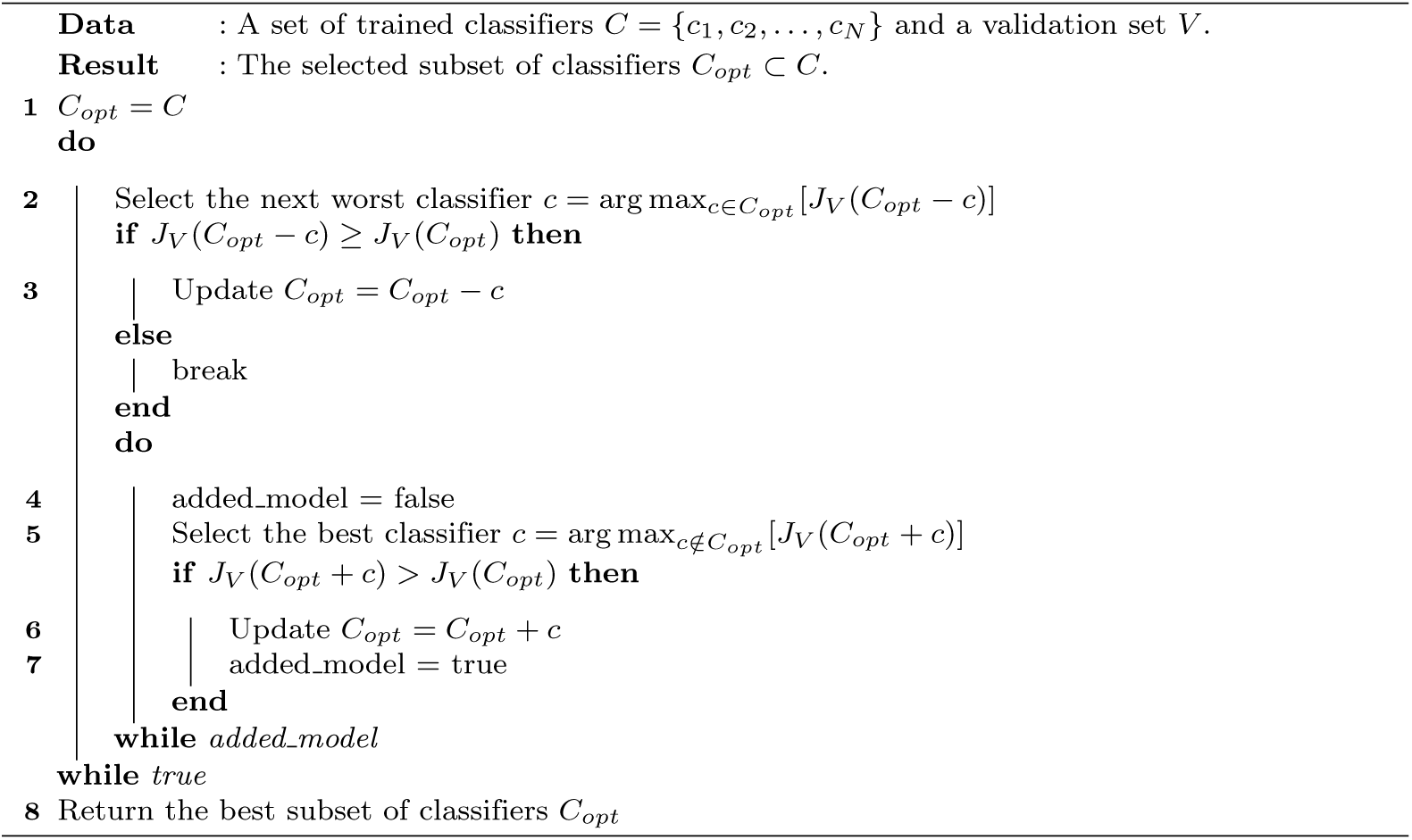

We can also consider randomized versions of SFFS and SFBS as possible improvements. These algorithms are a combination of the random generation of a subset of models and a floating search selection algorithm. We propose the use of Random Sequential Forward Floating Selection (RSFFS) and Random Sequential Backward Floating Selection (RSBFS). Both algorithms start with a random subset of models^1^, and with this random subset, an SFFS or SBFS algorithm is implemented. Thus, in the experiment, we will consider the SFS, SBS, LRS (with *l* = 3, *r* = 1 and *l* = 1, *r* = 3), SFFS, SBFS, RSFFS and RSBFS algorithms.

As we are combining various models, there are many ways of combining the outputs of those models. For the combination, we use three simple methods as our major aim is efficient execution. Although there are more complex approaches [22], their advantage is not large due to over-fitting problems. These methods are: i) the sum of the outputs of the classifiers; ii) the majority voting; and iii) the maximum output, where the sequence is classified using only the model with the highest output. In the machine learning literature, combining different sources of evidence for a classification problem is a common task [23]. Although various sophisticated methods have been developed for combining many classifiers [24–27]; in practice, none of them are able to significantly outperform the simpler methods on a regular basis.

Two of the problems of combining many different classification models that are trained on different datasets are that their outputs may not be in the same range and the optimal classification threshold might be different for each model. The problem of the different ranges is solved by scaling all of the outputs to the interval [−1, 1]. Regarding the threshold, we obtain the optimal threshold for each model, which is denoted as Θ*_opt_*, using the validation set, and for the inclusion of the model in any combination, we use *y*(**x**) − **Θ***_opt_*, where *y*(**x**) is the actual output of the model for sequence **x**.

For the training stage, we can select as many species as we deem useful for our problem. We need not select the most appropriate species because the floating search will discard the useless classifiers. Once we have selected the set of species whose genomes we are going to use, we train as many classifiers as we want from those species. For every organism, we can train various classifiers, such as support vector machines (SVMs), neural networks (NNs), decision trees (DTs), and the *k-*Nearest Neighbor (*k*-NN) rule, and the same classifiers with different parameters. Because the validation stage can consider hundreds of classifiers, any method of potential interest can be used. Again, the floating search process will remove unneeded classifiers.

### Experimental setup

To test our model, we chose the human genome together with those of other 20 species. Our aim was to test whether any species, regardless of the similarity of its genome with the human genome, could be useful. The following species were considered: *Anolis carolinensis, Bos primigenius taurus, Caenorhabditis elegans, Callithrix jacchus, Canis lupus familiaris, Danio rerio, Drosophila melanogaster, Equus caballus, Ficedula albicollis, Gallus gallus, Homo sapiens, Macaca mulatta, Monodelphis domestica, Mus musculus, Ornithorhynchus anatinus, Oryctolagus cuniculus, Pan troglodytes, Rattus norvegicus, Schistosoma mansoni, Sus scrofa* and *Takifugu rubripes*. These genomes were selected to consider a wide variety of organisms whose genomes are fully annotated.

Five classifiers were trained from every dataset for the four functional sites: a decision tree, a *k-*nearest neighbor rule, a positional weight matrix, a support vector machine with a string kernel and a support vector machine with a spectrum kernel. Additionally, for TIS and stop codon recognition, we used the stop codon method [28]. The parameters for every classifier were obtained using 10-fold cross-validation.

To evaluate our approach, we used five human chromosomes for testing purposes, namely, chromosomes 1, 3, 13, 19 and 21, and we used chromosome 16 for validation purposes. For each chromosome, we trained the classifiers with all of the remaining chromosomes except 16 and obtained the best combination method using our approach, and we used chromosome 16 for validation. We tested the selected models with all of the true TIS, donor and acceptor sites and stop codons and all of the negative samples of the given chromosome. That is, for chromosome 1, we trained the models with chromosomes 2 to 22 and X and Y, leaving out chromosome 16. Then, we chose the best combination method using chromosome 16 and tested this combination of models using chromosome 1. A summary of these datasets is shown in Table 1. The chromosomes were selected with the aim of choosing chromosomes of different lengths and coding densities. Chromosome 16 was chosen as a validation set because it is a chromosome of average length and coding density. We used the CCDS Update Released for Human of September 7, 2011. This update uses Human NCBI build 37.3 and includes a total of 26,473 CCDS IDs, which correspond to 18,471 GeneIDs. The validation set consisted of 836 positives samples and 2,721,460 negative samples for TIS, 28,567 positive samples and 8,011,785 negative samples for donor sites, 28,567 positive samples and 11,448,673 negative samples for acceptor sites and 838 positive samples and 7,480,457 negative samples for stop codons.

**Table 1.** Summary of datasets for chromosomes 1, 3, 13, 19 and 21. Random undersampling was used for training; thus, the number of negative instances was equal to the number of positive instances for the training dataset.

| Dataset | Site | Training data | Testing data |  |
| --- | --- | --- | --- | --- |
|  |  | Positives/Negatives | Positives | Negatives |
| Chr. 1 | TIS | 17,638 | 2,156 | 8,074,590 |
|  | STOP | 17,404 | 2,154 | 23,573,031 |
|  | DONOR | 630,985 | 81,378 | 22,634,283 |
|  | ACCEPTOR | 630,985 | 81,378 | 32,121,966 |
| Chr. 3 | TIS | 18,631 | 1,163 | 7,291,951 |
|  | STOP | 18,444 | 1,114 | 21,522,500 |
|  | DONOR | 663,884 | 48,479 | 19,578,976 |
|  | ACCEPTOR | 663,884 | 48,479 | 26,998,110 |
| Chr. 13 | TIS | 19,454 | 340 | 3,664,164 |
|  | STOP | 19,225 | 333 | 10,878,302 |
|  | DONOR | 696,352 | 16,011 | 9,613,960 |
|  | ACCEPTOR | 696,352 | 16,011 | 12,871,316 |
| Chr. 19 | TIS | 18,383 | 1,411 | 1,698,891 |
|  | STOP | 18,136 | 1,422 | 4,665,804 |
|  | DONOR | 678,673 | 33,690 | 5,673,086 |
|  | ACCEPTOR | 678,673 | 33,690 | 8,298,325 |
| Chr. 21 | TIS | 19,561 | 233 | 1,303,634 |
|  | STOP | 19,558 | 237 | 3,726,959 |
|  | DONOR | 704,725 | 7,638 | 3,555,622 |
|  | ACCEPTOR | 704,725 | 7,638 | 4,819,053 |

One of the key aspects of the evaluation of any newly proposed method is the set of previous methods that are considered in the comparison. Many methods have been proposed for recognizing functional sites [2, 28–30]. However, these previous works and our own research [7, 31] have shown that an SVM with a string kernel is the best state-of-the-art method for TISs, stop codons and splice sites [6]. To evaluate the general advantage of SVMs with string kernels, we performed a preliminary study of the available methods, which included position weight matrices, decision trees, *k-*nearest neighbors, the stop codon method [28], Wang et al.’s method [30], Salzberg’s method [32] and SVMs with linear and Gaussian kernels and four string kernels: the locality improved (LI) kernel, the weighted degree kernel (WD), the weighted degree kernel with shifts [33] (WDS) and the spectrum kernel [34]. SVMs with WD kernels consistently provided the best results. Thus, we chose this method as the method to be compared with our proposed method. WDS provided marginally better results than WD, but with a far higher computational complexity. To ensure a fair comparison, we considered not only these methods but also all of the others that were used as classifiers. Then, for every experiment, we compared our approach to the best performing method in terms of the validation performance. SVM with WD kernel was always the best individual classifier.

Another key parameter of the learning process is the window around the functional site that is used to train the classifiers. An additional advantage of our approach is that it allows the use of a suitable window for each dataset and even the combination of models that are trained using different windows. The value of the window for each classifier was obtained by cross-validation. We considered the site to be offset by 0 and tested the performance of the following windows: [−100, 0], [−75, 25], [−50, 0], [−50, 50], [−25, 0], [−25, 25], [−25, 75], [−10, 15], [−10, 40], [−10, 90], [0, 25], [0, 50] and [0, 100]. For each trained classifier, the best window was chosen. For the stop codon method, we used the additional window values of [0, 200], [0, 300], [0, 400] and [0, 500] for TIS recognition and the window values of [−200, 0], [−300, 0], [−400, 0] and [−500, 0] for stop codon recognition. For donor and acceptor sites, due to the many training instances, validation of the window around the site was not feasible. Thus, we chose a fixed window for both sites of [−25, 25].

Furthermore, SVMs are very sensitive to the learning parameters; thus, we also performed cross-validation to obtain their values. The WD kernel has two parameters: the standard *C* parameter of any SVM and the window width of the string kernel. We tested values of 1, 10, 100 and 1000 for C and 12 and 24 for the window width. All 8 combinations were evaluated using 10-fold cross-validation, and the best combinations was chosen. Although it can be argued that this method might result in suboptimal parameters, it represents a good compromise between the high performance of SVM and the high computational cost of evaluating each set of parameters. The spectrum kernel is too time consuming for cross-validation of the parameters in the same way as the WD kernel. Therefore, we fixed the values of the kernel to the values that are recommended by the authors [34] and only validated the value of *C* using the same values as for the WD kernel.

For PWM and C4.5, there are no parameters that have a significant effect on their performance. For *k-*NN, the number of neighbors *k* was chosen by cross-validation in the interval [1, 100].

To train the models, we used random undersampling [35] because previous studies have demonstrated its usefulness for TIS recognition [31]. For random undersampling, we used a ratio of 1, which means that the majority class was randomly undersampled until both classes had the same number of instances. To avoid any contamination of the experiments, for every training set, regardless of the species, we removed the genes that were shared with the test chromosome for all the training datasets.

To evaluate the obtained classifiers, we used the standard measures for imbalanced data. Given the number of true positives (TP), false positives (FP), true negatives (TN) and false negatives (FN), we used the sensitivity (Sn):

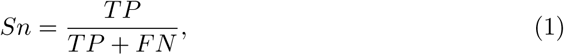

and the specificity (Sp):

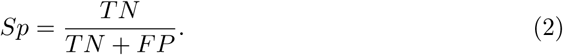

The geometric mean of these two measures, namely, 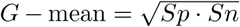, was our first classification metric. As a second measure, we used the area under the receiver operating characteristic (ROC) curve (auROC). However, auROC is independent of the class ratios and can be less meaningful when we have very unbalanced datasets [6]. In such cases, the area under the precision-recall curve (auPRC) can be used. The recall measure is equivalent to the sensitivity measure that was defined above. The precision (P) is given by:

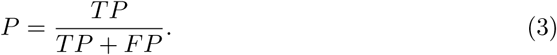

The auPRc measure is especially relevant if we are mainly interested in the positive class. However, the auPRc measure can be very sensitive to subsampling. In our results, we use all the positive and negative instances for each of the five tested chromosomes; thus, no subsampling is used. This also yields small auPRC values.

We use these three metrics because they provide two views of the performance of the classifiers. The auROC and auPRC values describe the general behavior of the classifier. However, when used in practice, we must establish a threshold for the classification of a query pattern. *G-*mean provides the required snapshot of the performance of the classifier when we set the required threshold.

The recognition of sites is usually a first step within a larger task, such as a gene structure prediction program. Therefore, depending on the subsequent steps, our focus was centered on obtaining models that perform well in terms of various accuracy measures. Thus, we performed experiments that were aimed at optimizing the three measures that were described above. We carried out experiments with eight search algorithms, namely, SFS, SBS, LRS (with *l* = 3, *r* =1 and *l* = 1, *r* = 3), SFFS, SBFS, RSFFS and RSBFS; three combination methods, namely, the sum of outputs, majority voting and maximum; and three measures as optimization objectives, namely, *G-*mean, auROC and auPRC. The best model was always selected using the validation set.

## Results and Discussion

We performed experiments on the recognition of TISs, donor and acceptor sites and stop codons to address the four most important sites in any gene recognition task. However, our approach is applicable to other recognition tasks, such as promoter and transcription start site (TSS) prediction. Our method has two main advantages: First, it has the ability to improve the performance of previous methods. Second, the chosen combination of classifiers that are trained on different genomes can provide information on which species are more interesting for human site recognition. In the following four sections, we discuss the results of the recognition of the four sites.

One of the main advantages of our approach is that we can optimize the performance measure in which we are interested, which can be the *G-*mean, auROC, auPRC or any other measure that is useful for our application. Thus, we conducted our experiments using three performance measures: *G-*mean, auROC and auPRC. The first relevant result is that the combination of the best models that was obtained for each measure was different. This result means that, depending on the aim of the work, different combinations of classifiers are needed. For each of the five studied chromosomes, we obtained three combinations of models, each optimized for one of the three measures that are discussed above.

### Results for TIS recognition

The results for the recognition of TISs for human chromosomes 1, 3, 13, 19 and 21 are shown in Table 2. Regarding the search method, the results for TIS support our approach of using different methods and selecting the best method for each case, as there is no clear winner. Although SFFS achieved the best results most often, SFS, LRS and RSFBS also perform well. For the combination method, the sum of outputs was always the best method for auROC and auPRC, with the exception of auROC for chromosome 13. For *G-*mean, majority voting was always the best-performing approach.

**Table 2.** Results for TIS recognition for human chromosomes 1, 3, 13, 19 and 21. The table shows the results for the three considered measures, auROCm auPRC and *G-*mean. The best search method, the best combination method and the classification performance values are shown.

| Chrom. | Objective | Method | Combination | G | auROC | auPRC | TP | FN | TN | FP |
| --- | --- | --- | --- | --- | --- | --- | --- | --- | --- | --- |
|  | State-of-the-art |  |  | 0.8528 | 0.9390 | 0.0701 | 1,697 | 459 | 7,460,252 | 614,338 |
| 1 | auROC | SFFS | Sum | 0.8782 | **0.9781** | 0.1296 | 1,688 | 468 | 7,954,516 | 120,074 |
|  | auPRC | SFFS | Sum | 0.8344 | 0.9690 | **0.1701** | 1,518 | 638 | 7,984,683 | 89,907 |
|  | G | LRS | Majority | **0.9284** | 0.9701 | 0.0157 | 1,956 | 200 | 7,671,456 | 403,134 |
|  | State-of-the-art |  |  | 0.8316 | 0.9265 | 0.0578 | 862 | 301 | 6,804,392 | 487,559 |
| 3 | auROC | SFFS | Sum | 0.8428 | **0.9732** | 0.1295 | 834 | 329 | 7,222,397 | 69,554 |
|  | auPRC | RSFBS | Sum | 0.8030 | 0.9623 | **0.1720** | 756 | 407 | 7,233,272 | 58,679 |
|  | G | SFFS | Majority | **0.9142** | 0.9284 | 0.0037 | 1,033 | 130 | 6,861,085 | 430,866 |
|  | State-of-the-art |  |  | 0.8520 | 0.9396 | 0.0575 | 264 | 76 | 3,425,104 | 239,060 |
| 13 | auROC | RSFBS | Majority | 0.8678 | **0.9748** | 0.0610 | 259 | 81 | 3,622,541 | 41,623 |
|  | auPRC | LRS | Sum | 0.8033 | 0.9670 | **0.1611** | 221 | 119 | 3,637,190 | 26,974 |
|  | G | SFFS | Majority | **0.9083** | 0.9239 | 0.0045 | 299 | 41 | 3,437,702 | 226,462 |
|  | State-of-the-art |  |  | 0.8437 | 0.9368 | 0.0997 | 1,084 | 327 | 1,574,213 | 124,678 |
| 19 | auROC | SFFS | Sum | 0.8748 | **0.9680** | 0.1555 | 1,114 | 297 | 1,646,904 | 51,987 |
|  | auPRC | LRS | Sum | 0.8496 | 0.9587 | **0.1841** | 1,048 | 363 | 1,650,949 | 47,942 |
|  | G | SFFS | Majority | **0.9229** | 0.9482 | 0.0107 | 1,335 | 76 | 1,529,295 | 169,596 |
|  | State-of-the-art |  |  | 0.8132 | 0.9183 | 0.0434 | 163 | 70 | 1,232,367 | 71,267 |
| 21 | auROC | SFS | Sum | 0.8462 | **0.9757** | 0.1098 | 169 | 64 | 1,287,106 | 16,528 |
|  | auPRC | LRS | Sum | 0.8339 | 0.9683 | **0.1658** | 164 | 69 | 1,287,965 | 15,669 |
|  | G | SFFS | Majority | **0.9274** | 0.9439 | 0.0036 | 215 | 18 | 1,215,086 | 88,548 |

In terms of auROC, our approach achieved a clear improvement over the SVM method alone. The improvement ranged from 3.32% for the worst case, namely, chromosome 19, to 5.74% for the best case, namely, chromosome 21. We must take into account that improvement refers to many sites being correctly classified compared to the standard approach. The standard approach obtained a total of 1,536,902 FPs; this number was reduced to 299,766, which means more than one million fewer FPs. For any gene recognition program, that would mean a far better point from which to start for constructing correct genes.

For auPRC, the improvement was more dramatic^2^. The improvement is greater than 10% for all five chromosomes. This is a remarkable result if we take into account the low values of auPRC for all methods. For *G-*mean, the results also showed a clear advantage of our method with an improvement of over 5% for the worst case.

The reported reduction is relevant because most current gene recognizers heavily rely on the classification of sites as a basic step; therefore, it is very likely that those genes whose TIS is not recognized would be completely missed by any gene recognizer. Our approach has the potential to significantly improve the accuracy of any annotation system.

Another interesting result is that the behaviors of the TPs, FNs, TNs and FPs values depended on the measure that we were optimizing. Thus, if we are interested in obtaining the best TP and FN results, we should select the optimization of *G-*mean. If our interest is in TNs and FPs, auPRC should be our objective. If we want a satisfactory overall behavior of the four measures, we should use auROC as our objective. The ability of our proposed method to offer such flexibility is an important asset in any practical application.

Once we established the usefulness of our proposed method in terms of performance, we examined the results in terms of the species that were involved in the best combinations. Table 3 shows the models that were selected for the best combination for each measure and each chromosome. Regardless of the optimized measure, there was no species that never appeared in the best combination. This result indicates that although the contributions of some species are more relevant than those of others, the information of all of the genomes was useful for the prediction of human TISs, even those species that are very distant relatives of humans. Another interesting result is that for the three measures, namely, auROC, auPRC and *G-*mean, the obtained combinations of models were substantially different. This result indicates that we must consider our aims before designing our classifier. In most previous works, that is not taken into account.

**Table 3.**
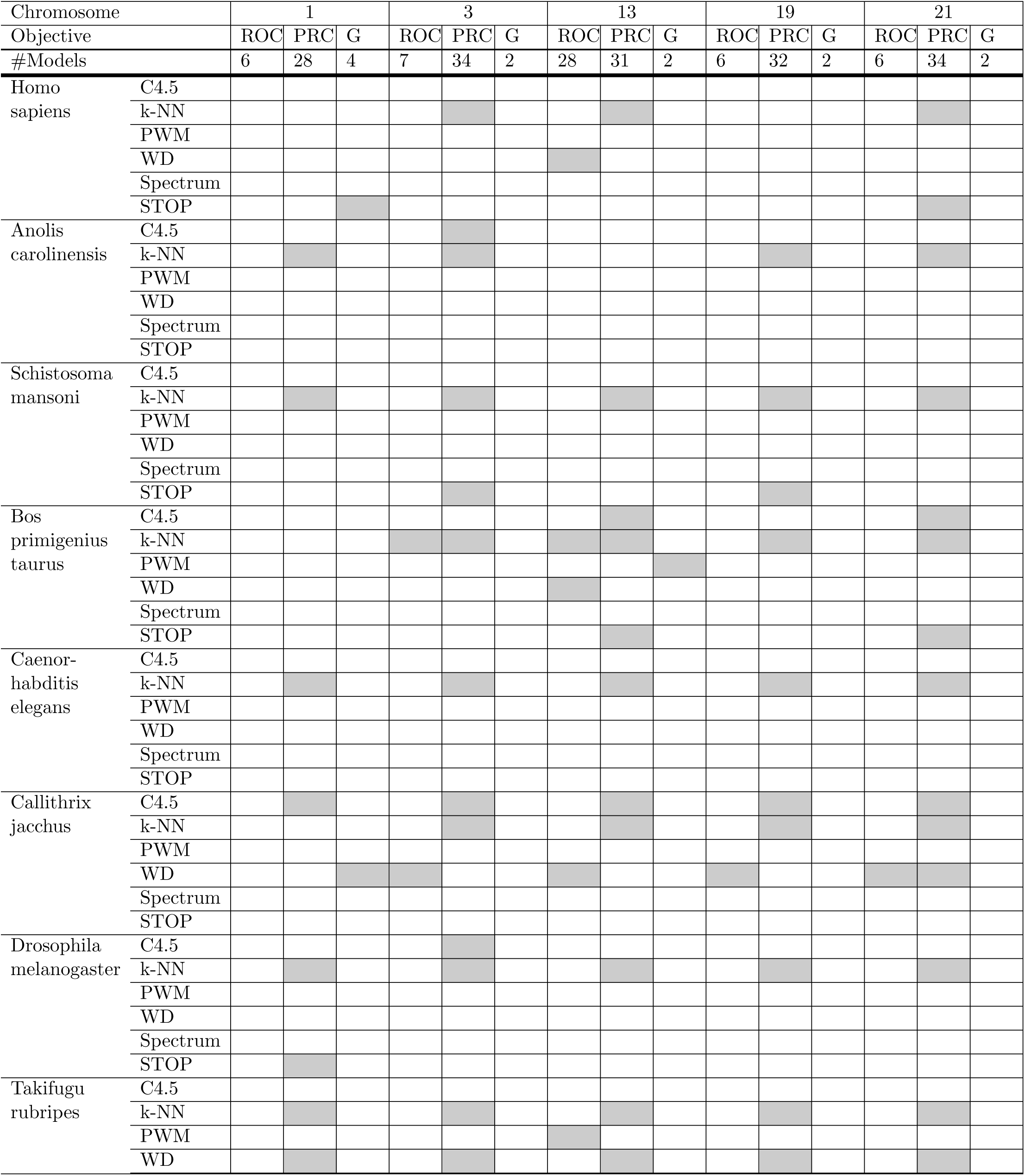
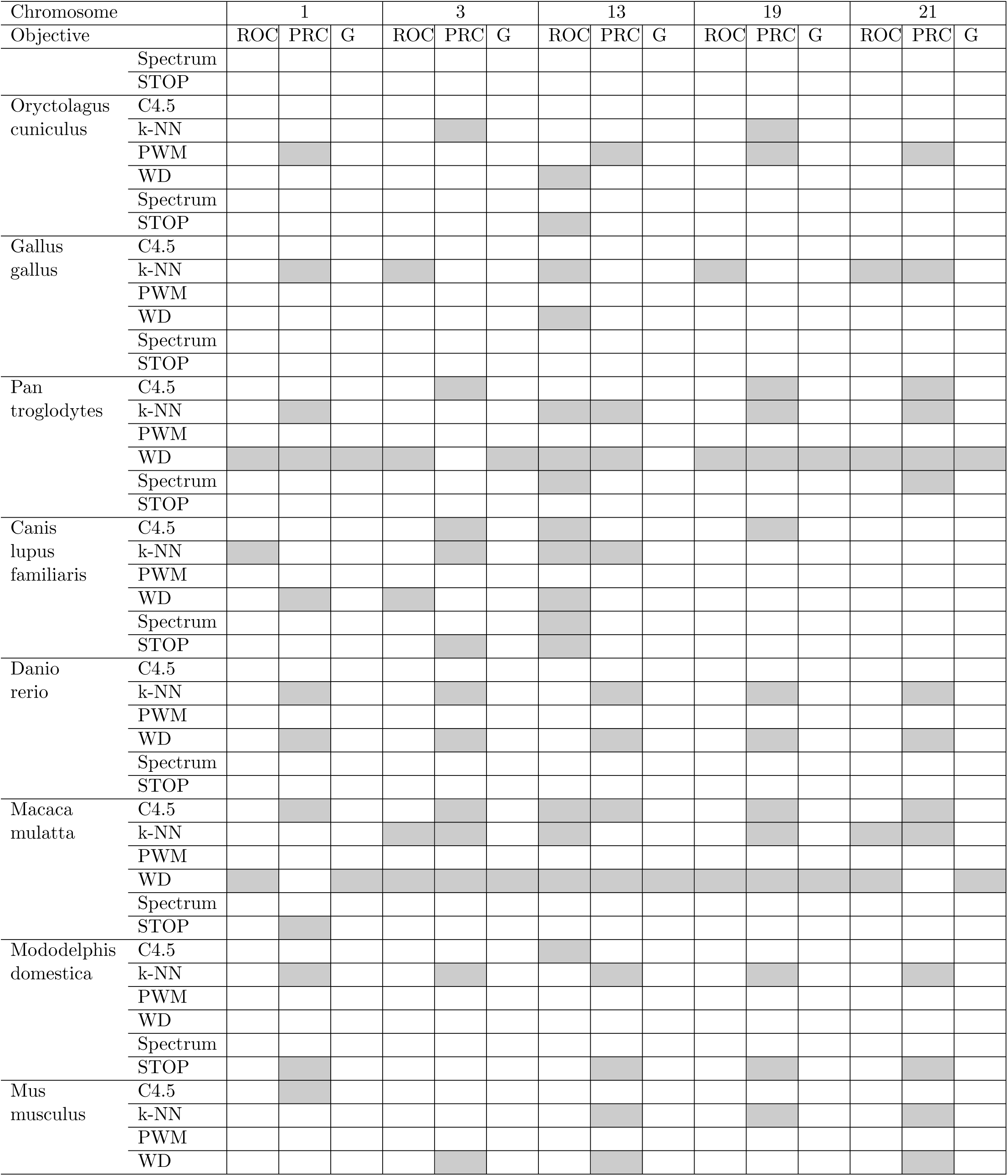
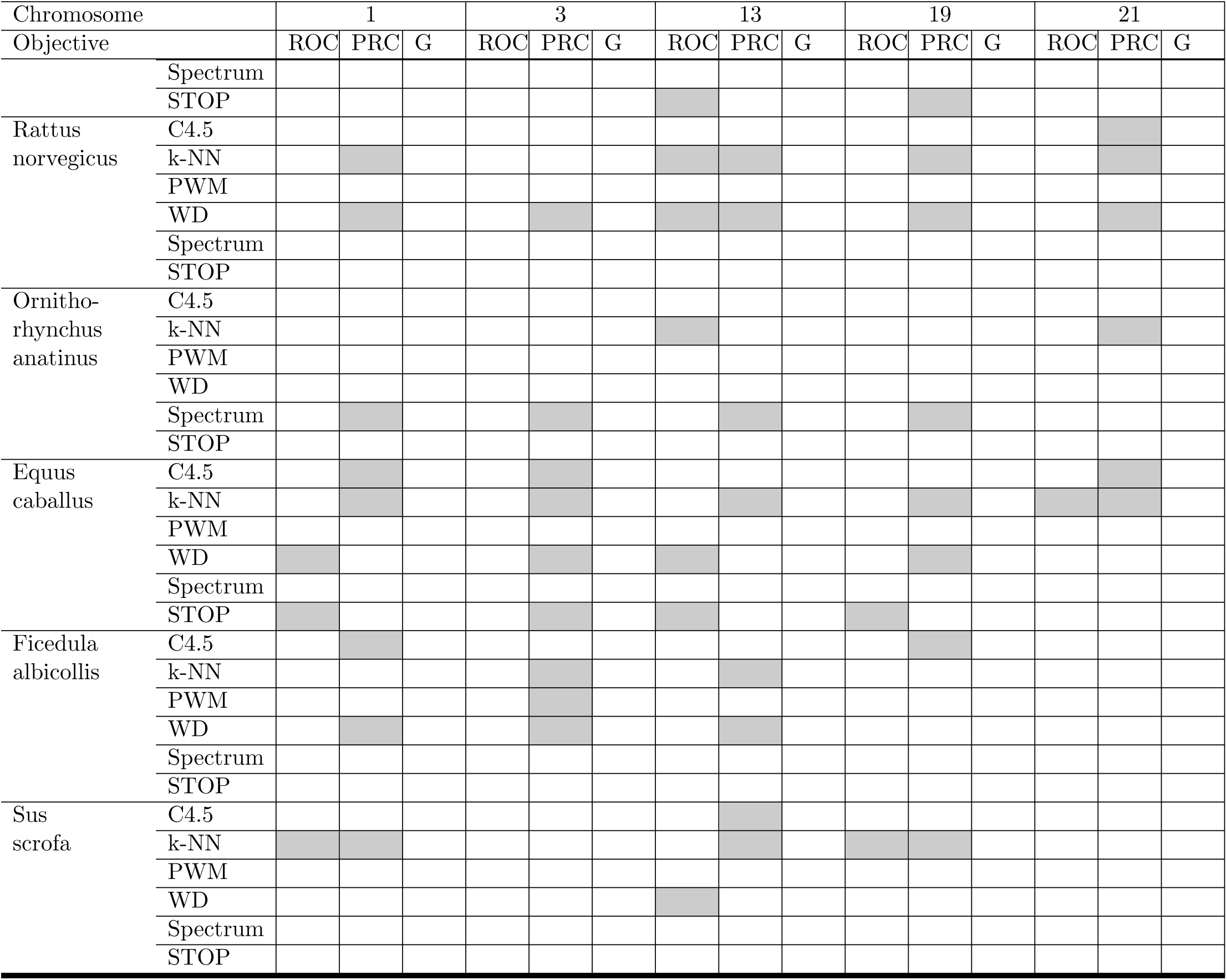
Models selected for TIS recognition.

Regarding the classification models, all methods were selected at least once. The *k-*NN rule and SVM with a string kernel were the most frequently selected methods. The case of *k-*NN is remarkable as this approach is not usually used for this task [2, 28–30]. It appears that the diversity that *k-*NN introduced into the models was useful for the overall performance of the combinations, despite that *k-*NN alone showed a worse performance than SVM alone. The explanation of this result may be found in the behavior of the ensembles of classifiers. It is well known [36] that a diverse ensemble of classifiers improves the performance of the set of classifiers.

The five most frequently used genomes were *Macaca mulatta, Pan troglodytes, Equus caballus, Callithrix jacchus* and *Rattus norvegicus. Homo sapiens* was not among the most often used genomes. Moreover, other genomes that are further removed from the human genome, such as *Takifugu rubripes*, were also frequently used.

With respect to the three objectives, optimizing the *G-*mean yielded the most stable results. For the five chromosomes, the selected models were always the SVM method for *Macaca mulatta* and *Pan troglodytes*, with the exception of chromosome 13,where *Pan troglodytes* was replaced with *Bos primigenius taurus*. For chromosome 1, another two classification models were used. For auROC, six or seven models were usually selected, with chromosome 13 requiring 28. The SVM method was always chosen for *Macaca mulatta* and *Pan troglodytes*, but the remaining methods and species depended on the chromosome. This is another interesting result because most TIS recognition programs mainly rely on common models for any task. Finally, for auPRC, significantly more models were selected, from 28 to 34, with a significant variation among the chromosomes. Here, the large number of negative samples made this task harder than optimizing the other two criteria.

The ROC and PRC curves are shown in Figs. 1–5. These figures show that our approach improved the auROC and auPRC for all five studied chromosomes. These results demonstrate that the proposed method outperformed the best model overall. The ROC and PRC curves show that the curves that correspond to our proposed method are always above the curves of the best model. This result indicates better performance for all the possible thresholds of classification.

**Figure 1.**
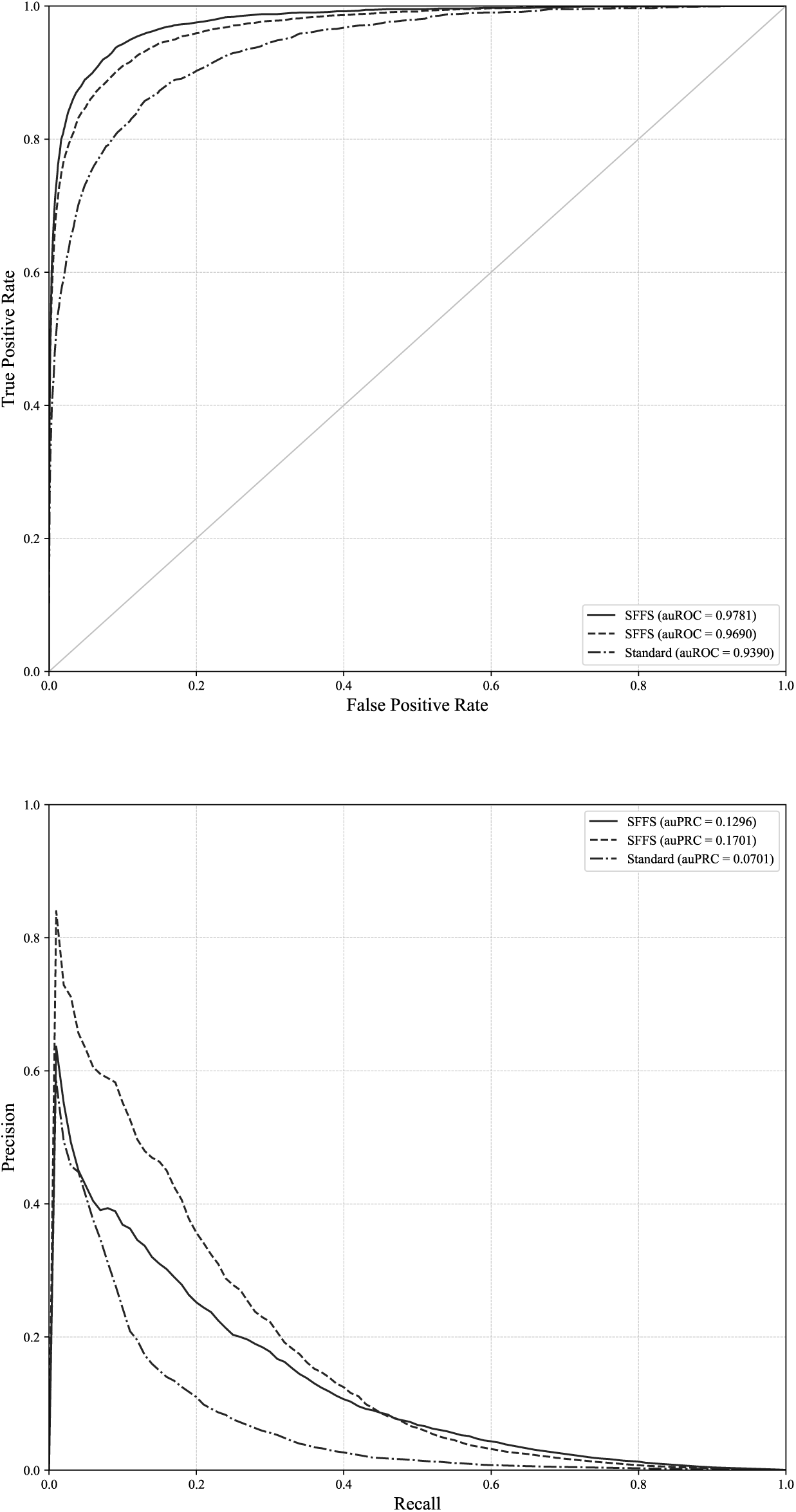
ROC and PRC curves for TIS and chromosome 1. ROC and PRC curves for chromosome 1 and the standard approach and our proposed method when auROC and auPRC are optimized.

**Figure 2.**
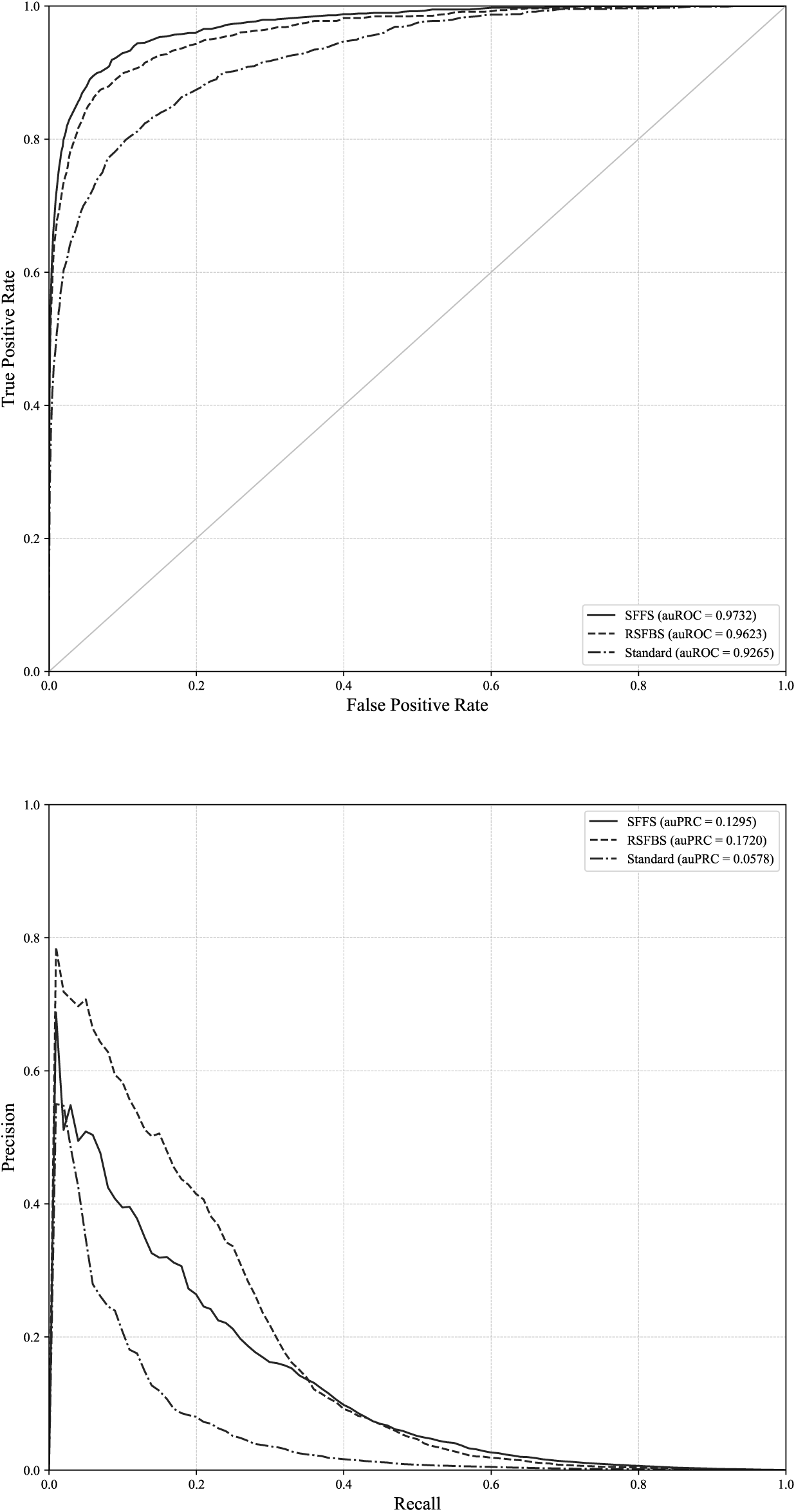
ROC and PRC curves for TIS and chromosome 1. ROC and PRC curves for chromosome 3 and the standard approach and our proposed method when auROC and auPRC are optimized.

**Figure 3.**
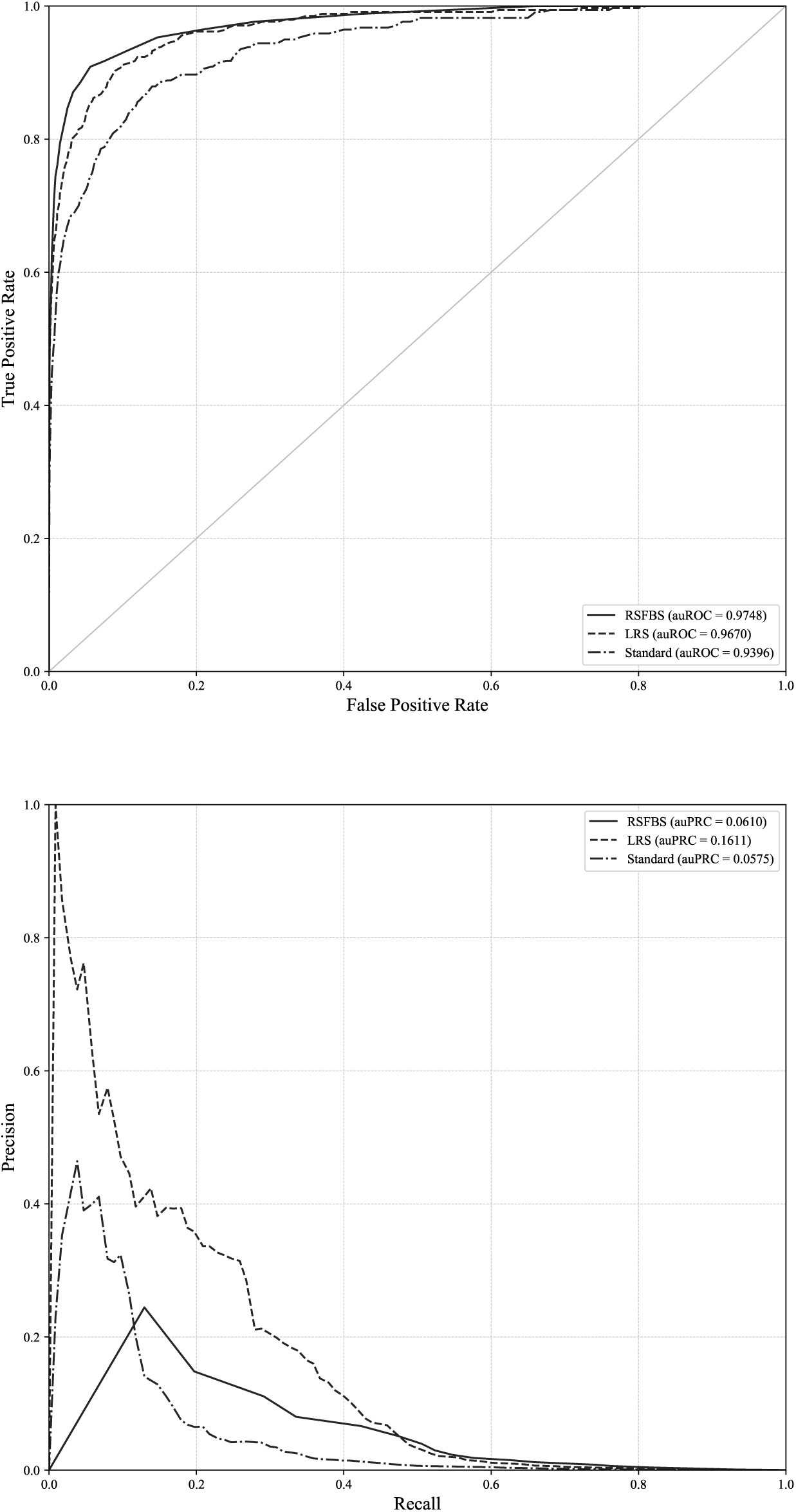
ROC and PRC curves for TIS and chromosome 1. ROC and PRC curves for chromosome 13 and the standard approach and our proposed method when auROC and auPRC are optimized.

**Figure 4.**
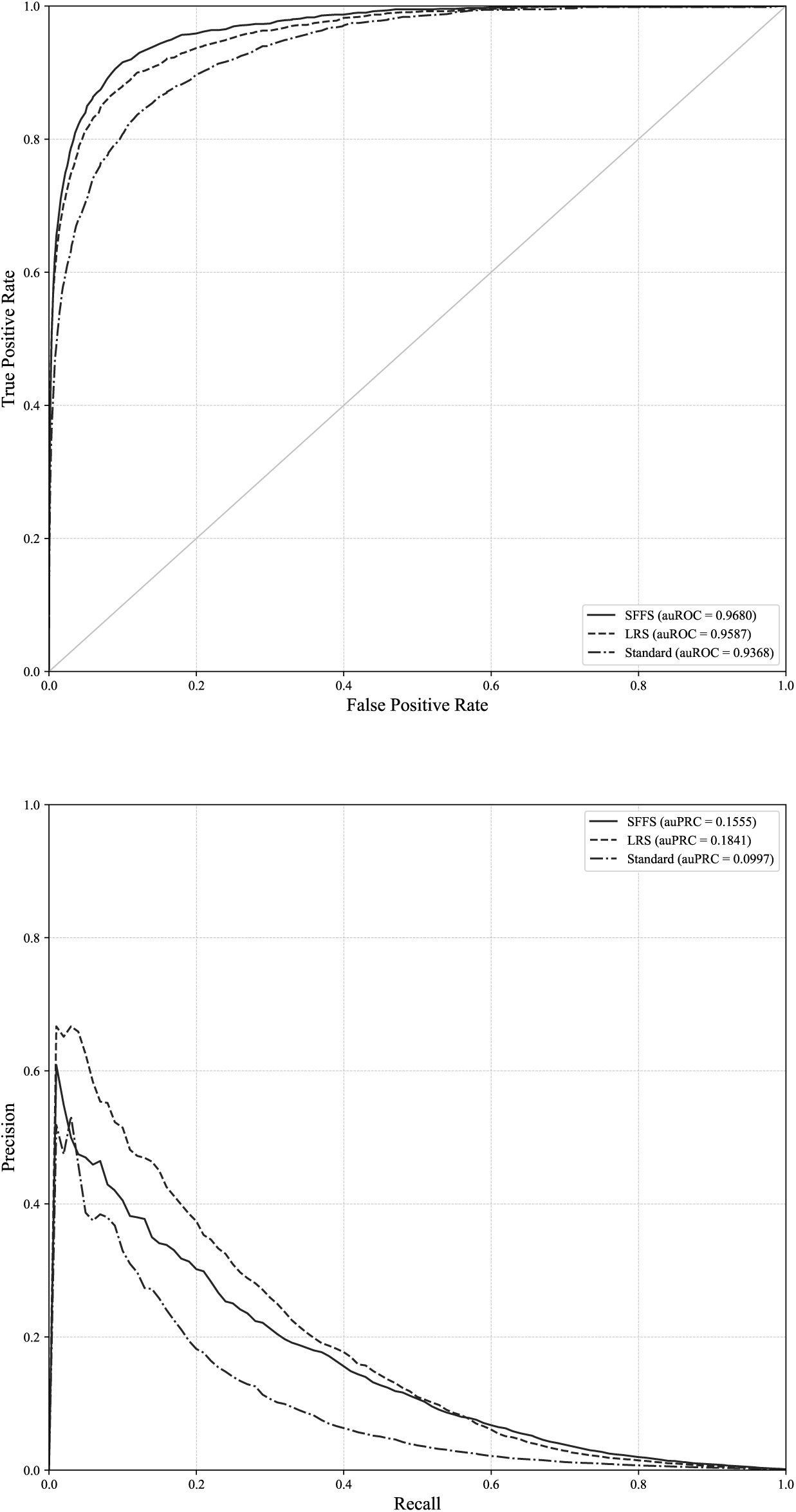
ROC and PRC curves for TIS and chromosome 19. ROC and PRC curves for chromosome 19 and the standard approach and our proposed method when auROC and auPRC are optimized.

**Figure 5.**
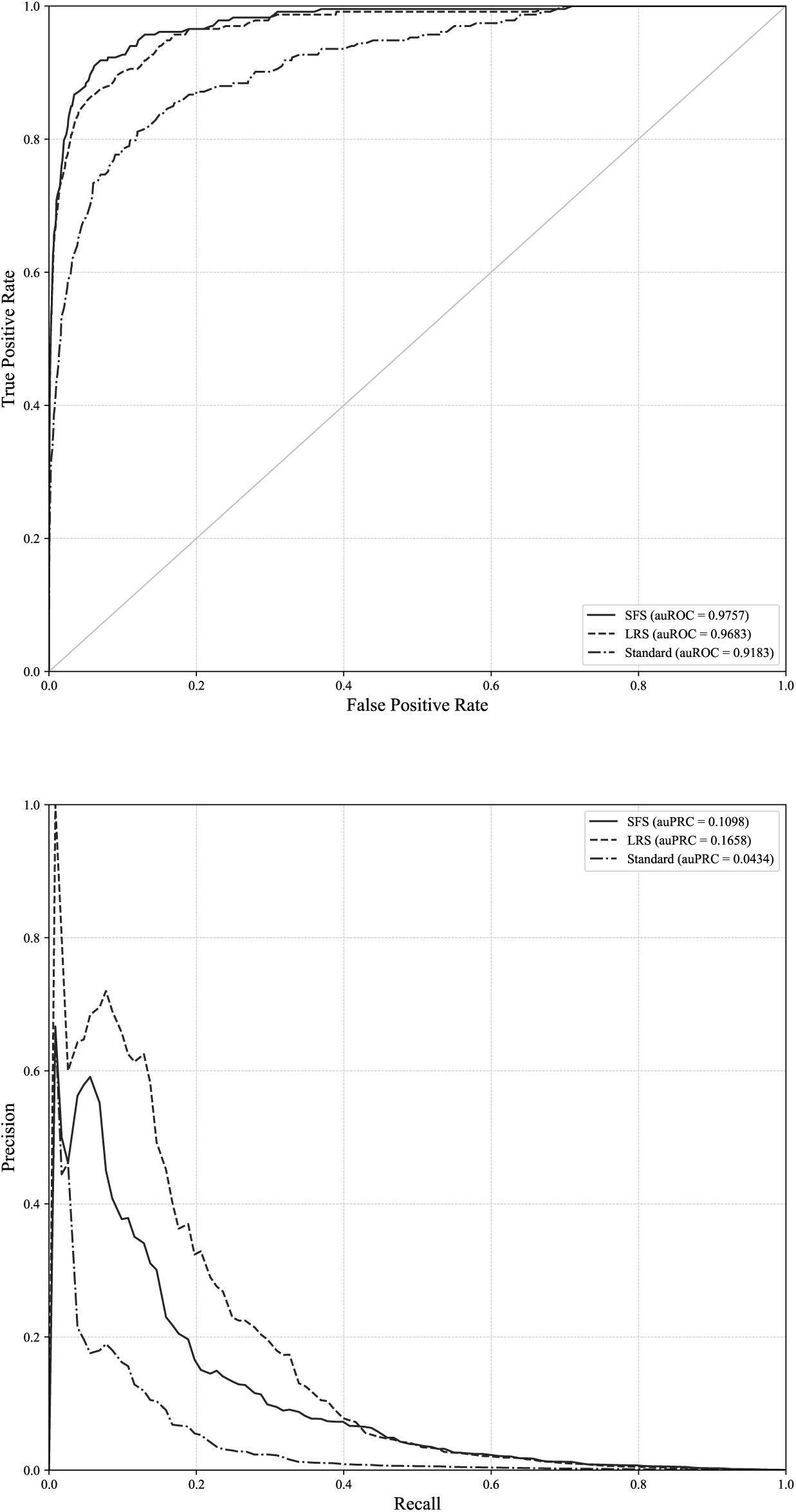
ROC and PRC curves for TIS and chromosome 1. ROC and PRC curves for chromosome 21 and the standard approach and our proposed method when auROC and auPRC are optimized.

### Results for donor site recognition

The results for the recognition of donor sites for human chromosomes 1, 3, 13, 19 and 21 are shown in Table 4. For auROC, the achieved results were close to or above 99%, so there was little room for improvement. A similar trend was followed by *G-*mean, with an improvement of approximately 1%. auPRC was significantly improved, from 5% in the worst case to 11% in the best case. However, since the number of negative samples was large, these small improvements corresponded to the correction of many erroneous predictions. For example, the standard approach obtained 3,616,750 FPs, while our approach for auROC optimization reduced this number by almost one million to 2,796,742 FPs.

**Table 4.**
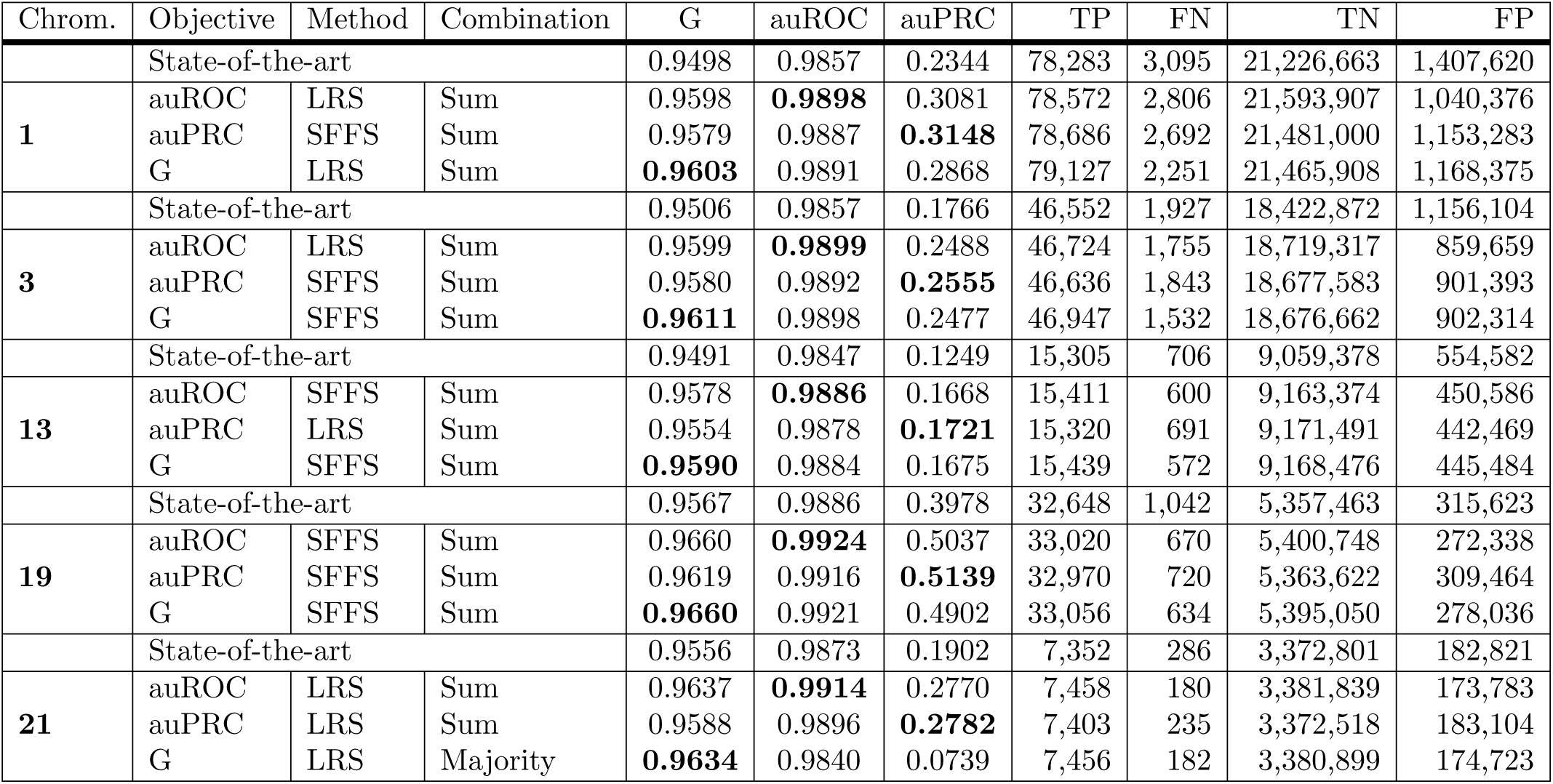
Results for donor site recognition for human chromosomes 1, 3, 13, 19 and 21.

| Chrom. | Objective | Method | Combination | G | auROC | auPRC | TP | FN | TN | FP |
| --- | --- | --- | --- | --- | --- | --- | --- | --- | --- | --- |
|  | State-of-the-art |  |  | 0.9498 | 0.9857 | 0.2344 | 78,283 | 3,095 | 21,226,663 | 1,407,620 |
| 1 | auROC | LRS | Sum | 0.9598 | **0.9898** | 0.3081 | 78,572 | 2,806 | 21,593,907 | 1,040,376 |
|  | auPRC | SFFS | Sum | 0.9579 | 0.9887 | **0.3148** | 78,686 | 2,692 | 21,481,000 | 1,153,283 |
|  | G | LRS | Sum | **0.9603** | 0.9891 | 0.2868 | 79,127 | 2,251 | 21,465,908 | 1,168,375 |
|  | State-of-the-art |  |  | 0.9506 | 0.9857 | 0.1766 | 46,552 | 1,927 | 18,422,872 | 1,156,104 |
| 3 | auROC | LRS | Sum | 0.9599 | **0.9899** | 0.2488 | 46,724 | 1,755 | 18,719,317 | 859,659 |
|  | auPRC | SFFS | Sum | 0.9580 | 0.9892 | **0.2555** | 46,636 | 1,843 | 18,677,583 | 901,393 |
|  | G | SFFS | Sum | **0.9611** | 0.9898 | 0.2477 | 46,947 | 1,532 | 18,676,662 | 902,314 |
|  | State-of-the-art |  |  | 0.9491 | 0.9847 | 0.1249 | 15,305 | 706 | 9,059,378 | 554,582 |
| 13 | auROC | SFFS | Sum | 0.9578 | **0.9886** | 0.1668 | 15,411 | 600 | 9,163,374 | 450,586 |
|  | auPRC | LRS | Sum | 0.9554 | 0.9878 | **0.1721** | 15,320 | 691 | 9,171,491 | 442,469 |
|  | G | SFFS | Sum | **0.9590** | 0.9884 | 0.1675 | 15,439 | 572 | 9,168,476 | 445,484 |
|  | State-of-the-art |  |  | 0.9567 | 0.9886 | 0.3978 | 32,648 | 1,042 | 5,357,463 | 315,623 |
| 19 | auROC | SFFS | Sum | 0.9660 | **0.9924** | 0.5037 | 33,020 | 670 | 5,400,748 | 272,338 |
|  | auPRC | SFFS | Sum | 0.9619 | 0.9916 | **0.5139** | 32,970 | 720 | 5,363,622 | 309,464 |
|  | G | SFFS | Sum | **0.9660** | 0.9921 | 0.4902 | 33,056 | 634 | 5,395,050 | 278,036 |
|  | State-of-the-art |  |  | 0.9556 | 0.9873 | 0.1902 | 7,352 | 286 | 3,372,801 | 182,821 |
| 21 | auROC | LRS | Sum | 0.9637 | **0.9914** | 0.2770 | 7,458 | 180 | 3,381,839 | 173,783 |
|  | auPRC | LRS | Sum | 0.9588 | 0.9896 | **0.2782** | 7,403 | 235 | 3,372,518 | 183,104 |
|  | G | LRS | Majority | **0.9634** | 0.9840 | 0.0739 | 7,456 | 182 | 3,380,899 | 174,723 |

Table 5 shows the classification models and genomes that were selected for every case. There are several differences with TIS recognition. First, there are a few genomes that were not used at all in any final best model, namely, *Anolis carolinensis, Drosopila melanogaster, Takifugu rubripes, Danio rerio, Monodelphis domestica* and *Ornithorhynchus anatinus*. Second, *G-*mean and auROC optimization required more models, whereas auPRC used significantly fewer models for TIS prediction. The models that were selected for every optimized measure showed a large variety, thereby supporting the claim of our work that as many genomes as available should be used instead of selecting some of them *a priori*.

**Table 5.**
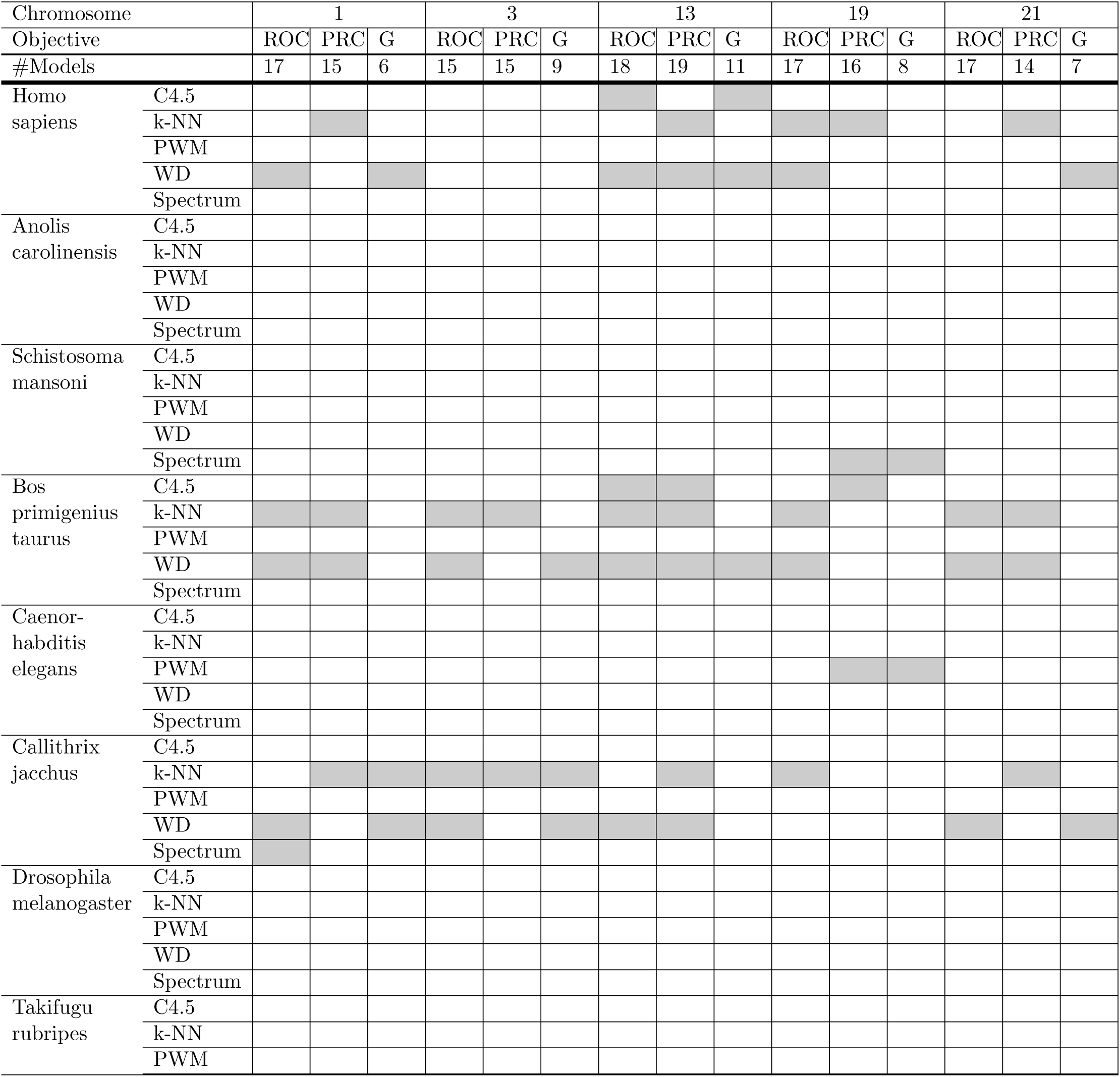
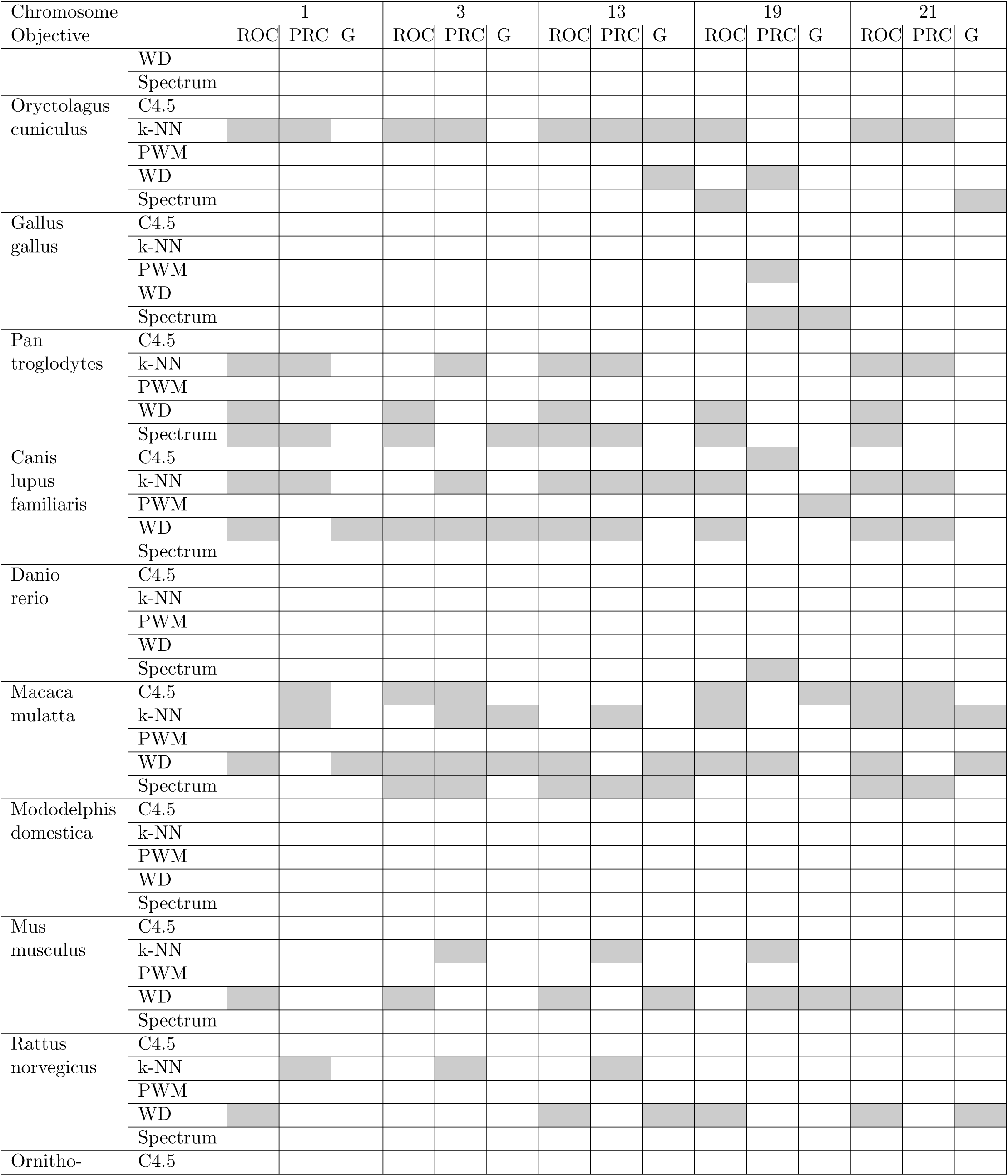
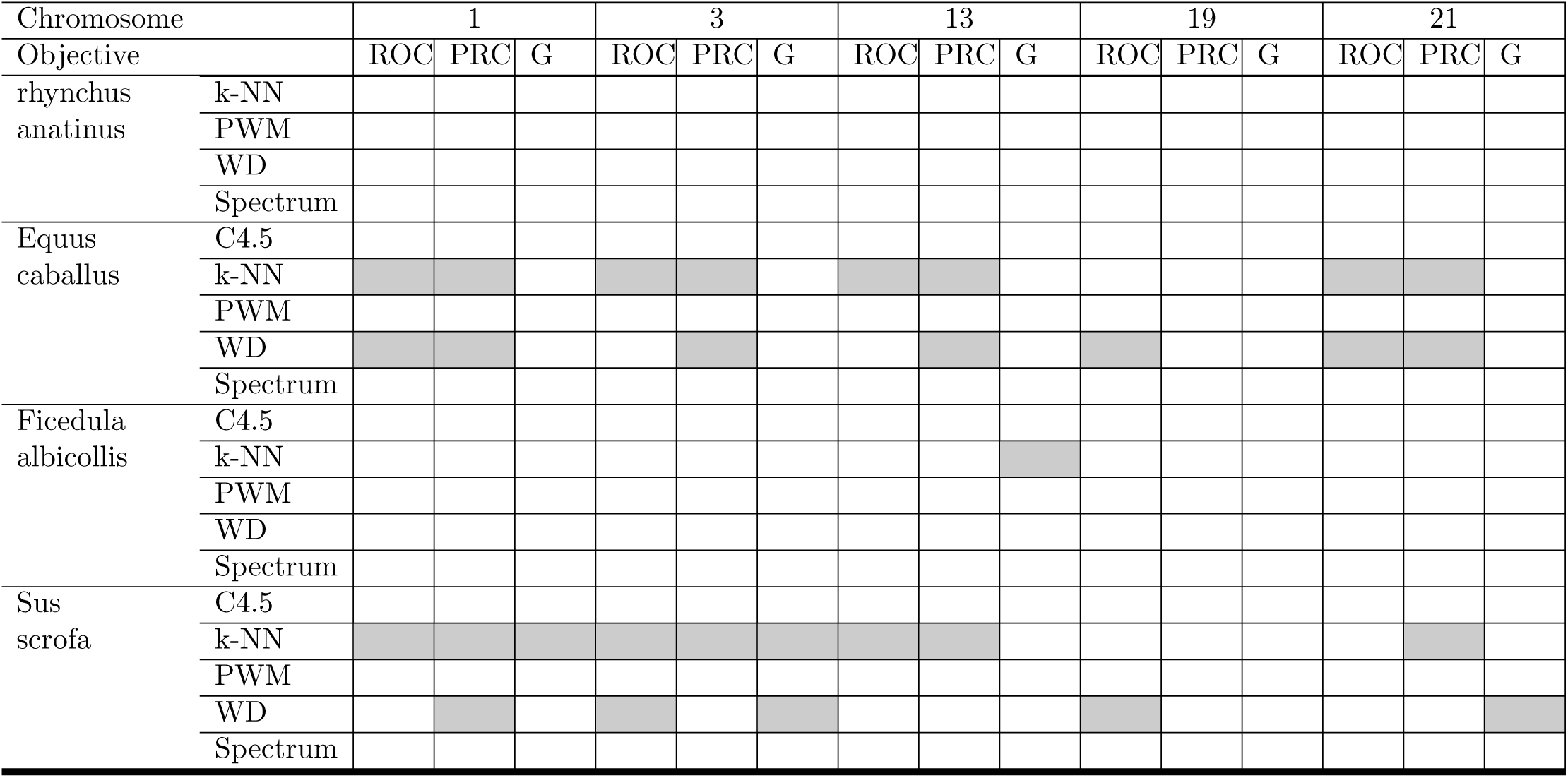
Models selected for donor site recognition.

Regarding the classification models, the behavior was more similar. *k-*NN and SVM with a string kernel were the models that were more frequently used, with a large difference with the remaining methods. PWM was never used, and C4.5 and the SVM with the spectrum kernel were used only on a few occasions. The ROC and PRC curves for our approach and the standard method are shown in Figs. 6 to 10.

**Figure 6.**
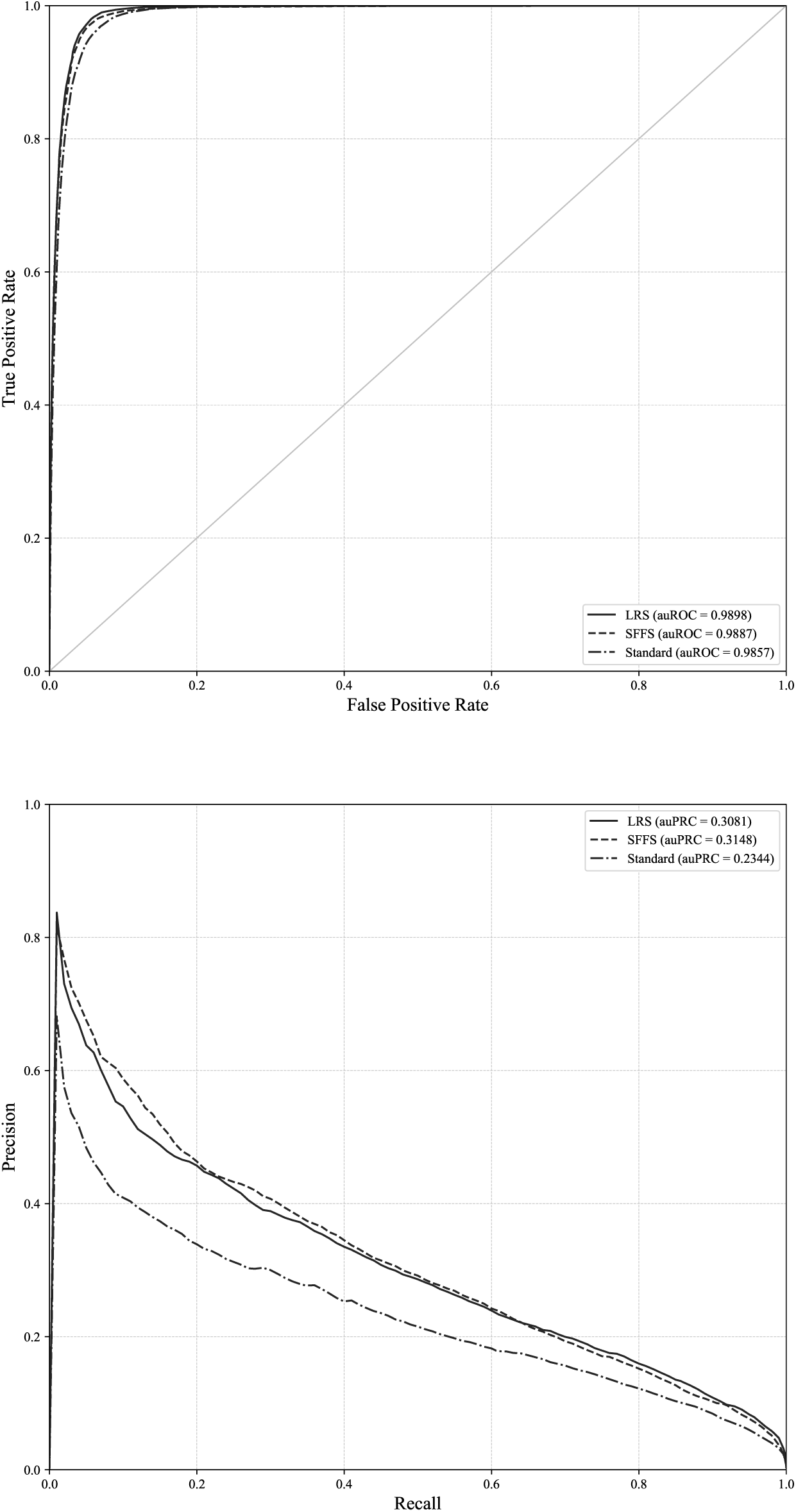
ROC and PRC curves for the donor site and chromosome 1. ROC and PRC curves for chromosome 1 and the standard approach and our proposed method when auROC and auPRC are optimized.

**Figure 7.**
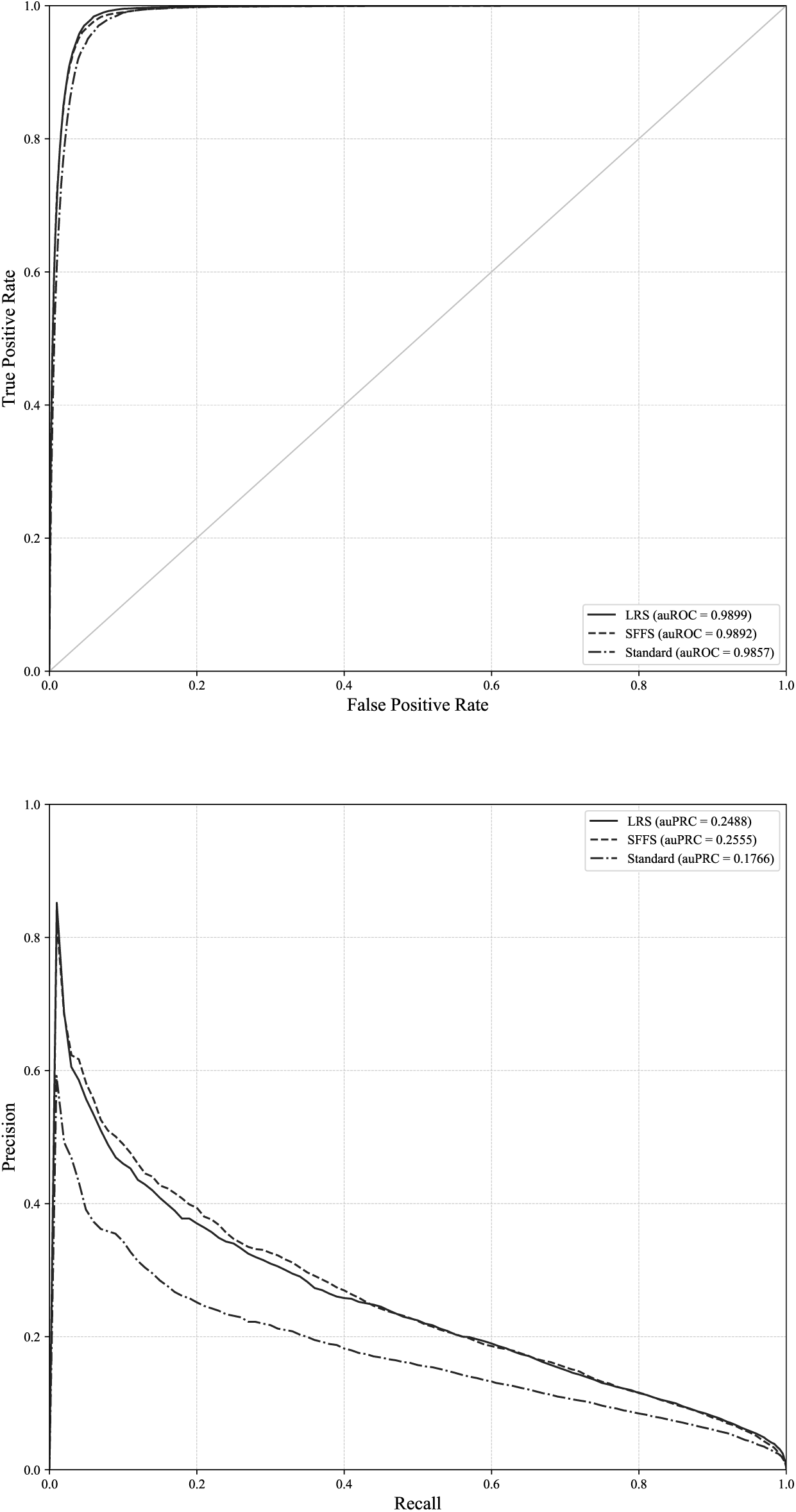
ROC and PRC curves for the donor site and chromosome 1. ROC and PRC curves for chromosome 3 and the standard approach and our proposed method when auROC and auPRC are optimized.

**Figure 8.**
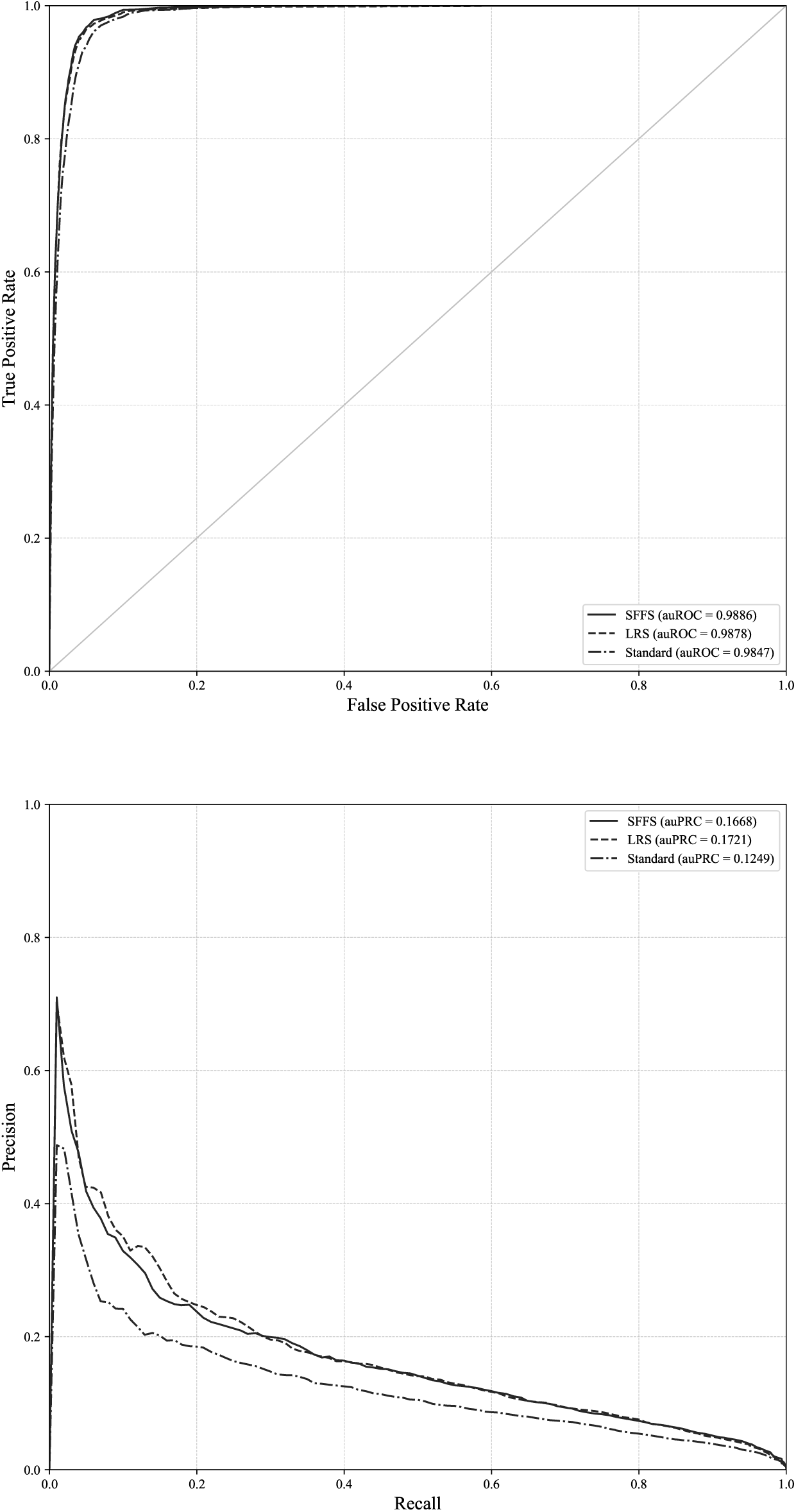
ROC and PRC curves for the donor site and chromosome 1. ROC and PRC curves for chromosome 13 and the standard approach and our proposed method when auROC and auPRC are optimized.

**Figure 9.**
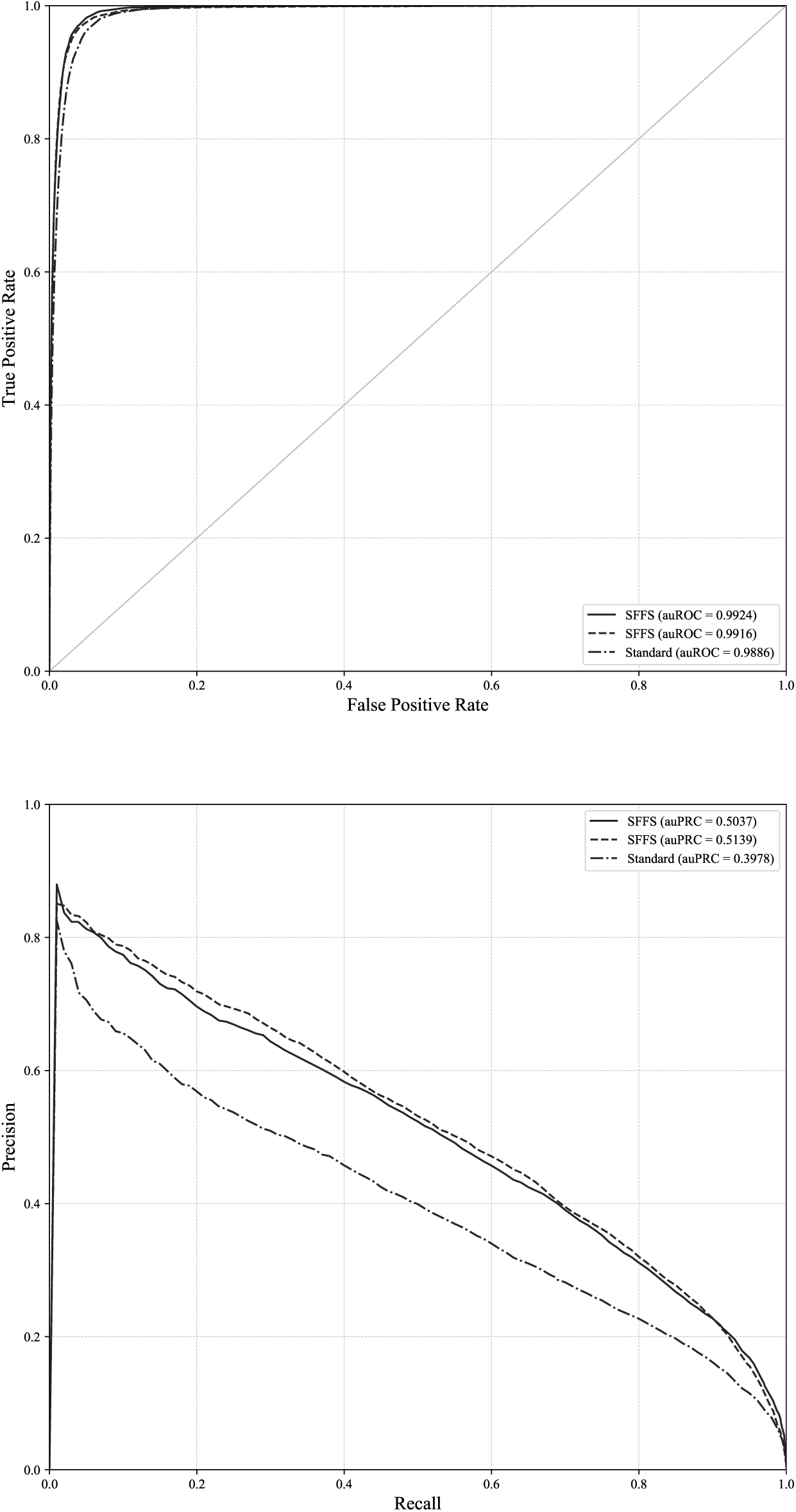
ROC and PRC curves for the donor site and chromosome 19. ROC and PRC curves for chromosome 19 and the standard approach and our proposed method when auROC and auPRC are optimized.

**Figure 10.**
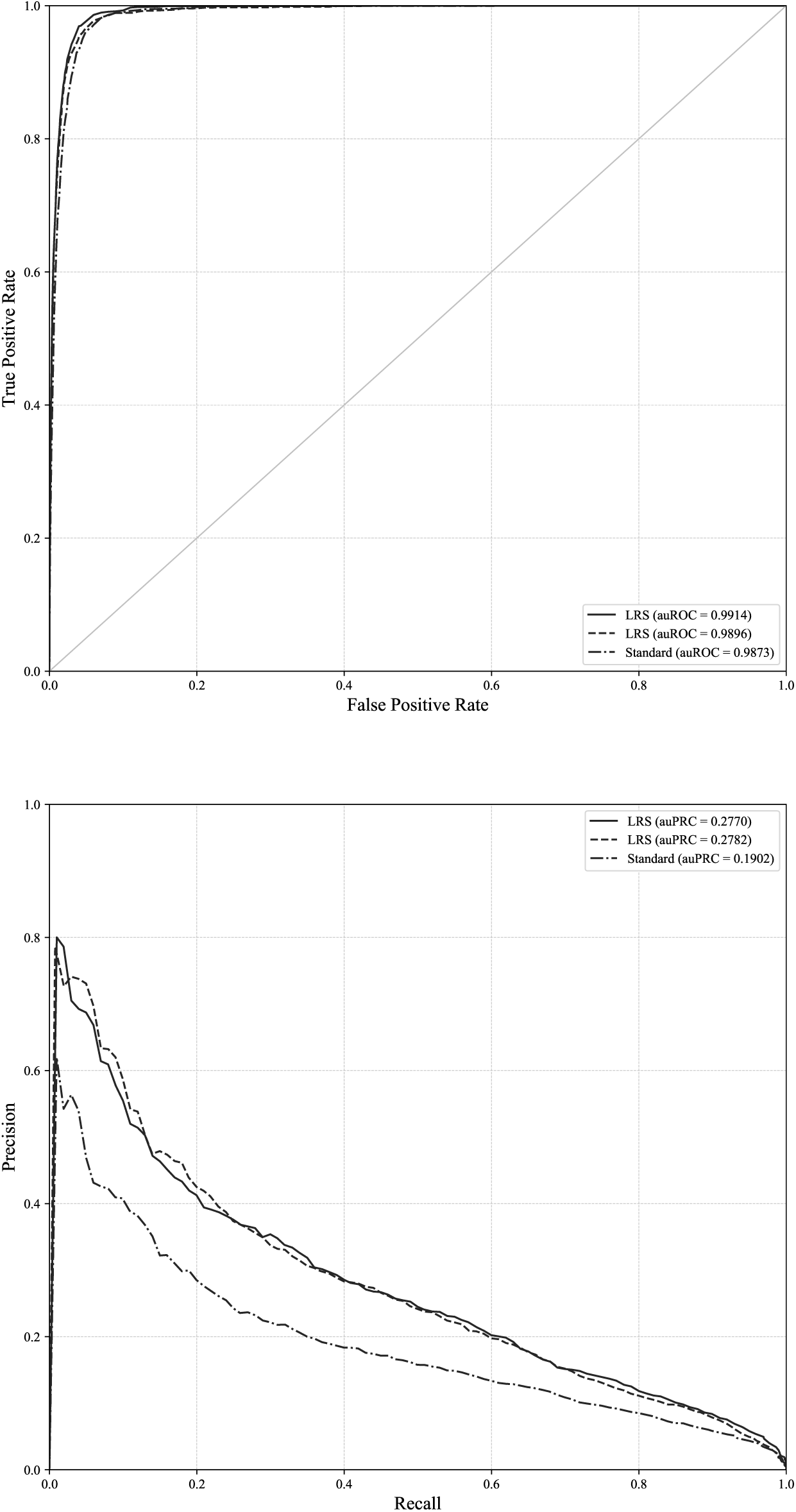
ROC and PRC curves for the donor site and chromosome 1. ROC and PRC curves for chromosome 21 and the standard approach and our proposed method when auROC and auPRC are optimized.

### Results for acceptor site recognition

The results for the recognition of acceptor sites for human chromosomes 1, 3, 13, 19 and 21 are shown in Table 6. The results for the acceptor site prediction were similar to those for the donor site prediction. The improvement in terms of auROC was small due to the little room for increasing the values of the standard method. However, the small improvement corresponded to many negative instances being correctly classified. The standard method obtained 6,053,645 FPs, whereas our approach when optimizing the auROC measure achieved 4,345,285 FPs, reducing by more than 1.5 million the total FPs.

**Table 6.** Results for acceptor site recognition for human chromosomes 1, 3, 13, 19 and 21.

| Chrom. | Objective | Method | Combination | G | auROC | auPRC | TP | FN | TN | FP |
| --- | --- | --- | --- | --- | --- | --- | --- | --- | --- | --- |
|  | State-of-the-art |  |  | 0.9442 | 0.9843 | 0.2242 | 78,145 | 3,233 | 29,823,601 | 2,298,365 |
| 1 | auROC | SFFS | Sum | 0.9562 | **0.9894** | 0.2912 | 78,284 | 3,094 | 30,530,255 | 1,591,711 |
|  | auPRC | LRS | Sum | 0.9550 | 0.9892 | **0.2940** | 78,179 | 3,199 | 30,496,996 | 1,624,970 |
|  | G | LRS | Majority | **0.9553** | 0.9827 | 0.0954 | 78,694 | 2,684 | 30,312,652 | 1,809,314 |
|  | State-of-the-art |  |  | 0.9397 | 0.9826 | 0.1620 | 46,158 | 2,321 | 25,037,728 | 1,960,382 |
| 3 | auROC | LRS | Sum | 0.9543 | **0.9885** | 0.2280 | 46,467 | 2,012 | 25,652,104 | 1,346,006 |
|  | auPRC | SFFS | Sum | 0.9537 | 0.9880 | **0.2274** | 46,485 | 1,994 | 25,607,518 | 1,390,592 |
|  | G | LRS | Sum | **0.9555** | 0.9884 | 0.2250 | 46,796 | 1,683 | 25,537,090 | 1,461,020 |
|  | State-of-the-art |  |  | 0.9428 | 0.9832 | 0.1123 | 15,357 | 654 | 11,928,112 | 943,204 |
| 13 | auROC | SFFS | Sum | 0.9561 | **0.9881** | 0.1517 | 15,496 | 515 | 12,157,237 | 714,079 |
|  | auPRC | SFFS | Sum | 0.9555 | 0.9881 | **0.1545** | 15,441 | 570 | 12,185,657 | 685,659 |
|  | G | SFFS | Sum | **0.9554** | 0.9880 | 0.1530 | 15,459 | 552 | 12,169,043 | 702,273 |
|  | State-of-the-art |  |  | 0.9455 | 0.9846 | 0.3016 | 32,213 | 1,477 | 7,758,186 | 540,139 |
| 19 | auROC | LRS | Sum | 0.9601 | **0.9906** | 0.4209 | 32,796 | 894 | 7,857,893 | 440,432 |
|  | auPRC | LRS | Sum | 0.9594 | 0.9905 | **0.4275** | 32,766 | 924 | 7,853,884 | 444,441 |
|  | G | LRS | Sum | **0.9596** | 0.9903 | 0.4171 | 32,752 | 938 | 7,860,411 | 437,914 |
|  | State-of-the-art |  |  | 0.9400 | 0.9840 | 0.1470 | 7,216 | 422 | 4,507,498 | 311,555 |
| 21 | auROC | SFFS | Sum | 0.9551 | **0.9887** | 0.2314 | 7,353 | 285 | 4,565,996 | 253,057 |
|  | auPRC | SBS | Sum | 0.9484 | 0.9870 | **0.2185** | 7,266 | 372 | 4,556,928 | 262,125 |
|  | G | LRS | Majority | **0.9531** | 0.9809 | 0.0621 | 7,360 | 278 | 4,543,109 | 275,944 |

The models that were selected for every chromosome are shown in Table 7. The selected classification methods and genomes were similar to those in the previous donor site recognition. The number of models for every measure was also similar, with the exception of auPRC for chromosome 21, which required more models, namely, 46. *Anolis carolinensis* was the only genome that was never used, although *Danio rerio, Drosophila melanogaster* and *Schistosoma mansoni* appeared only rarely. As in previous results, *k-*NN and SVM with a string kernel were the most commonly selected classification methods.

**Table 7.**
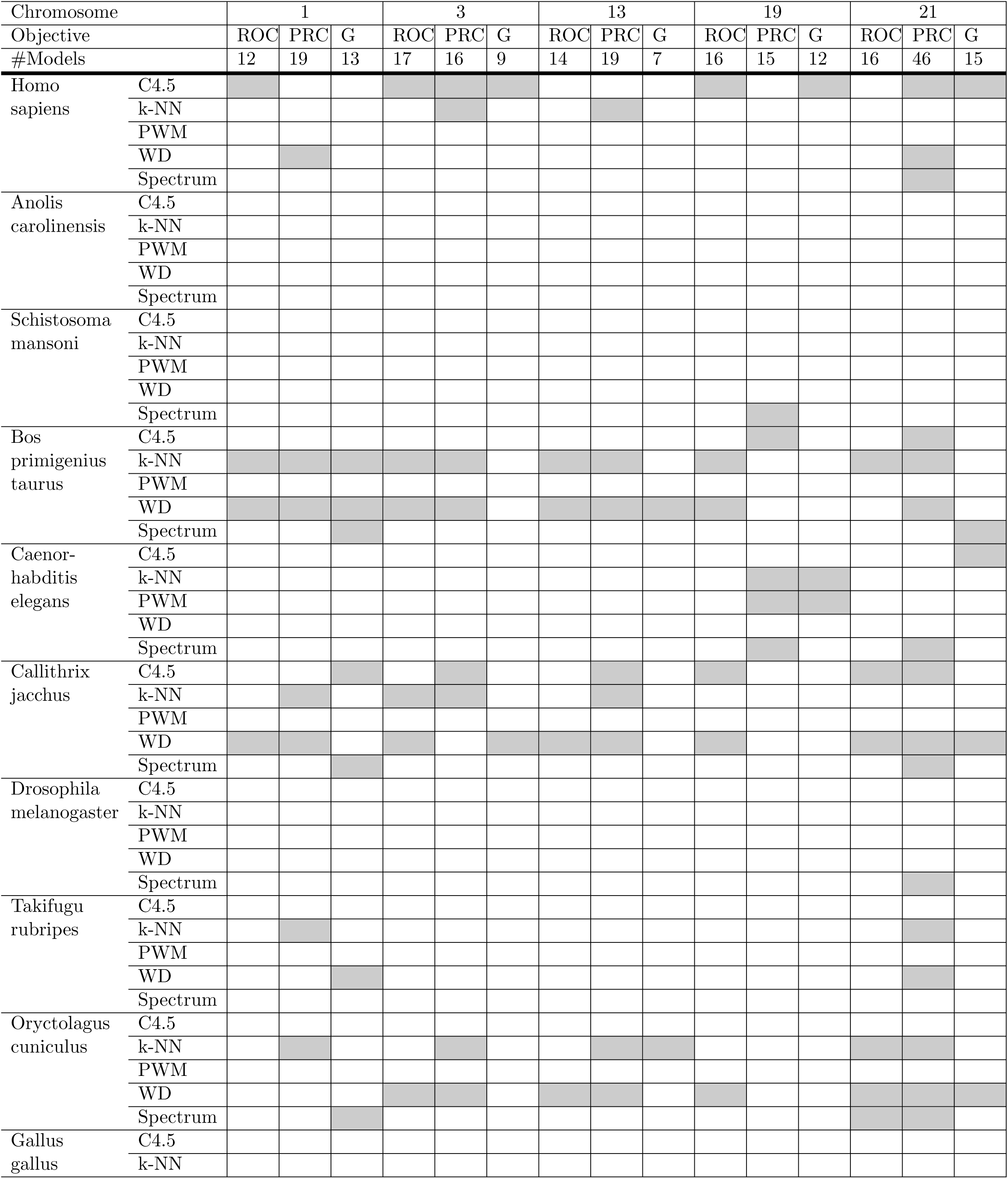
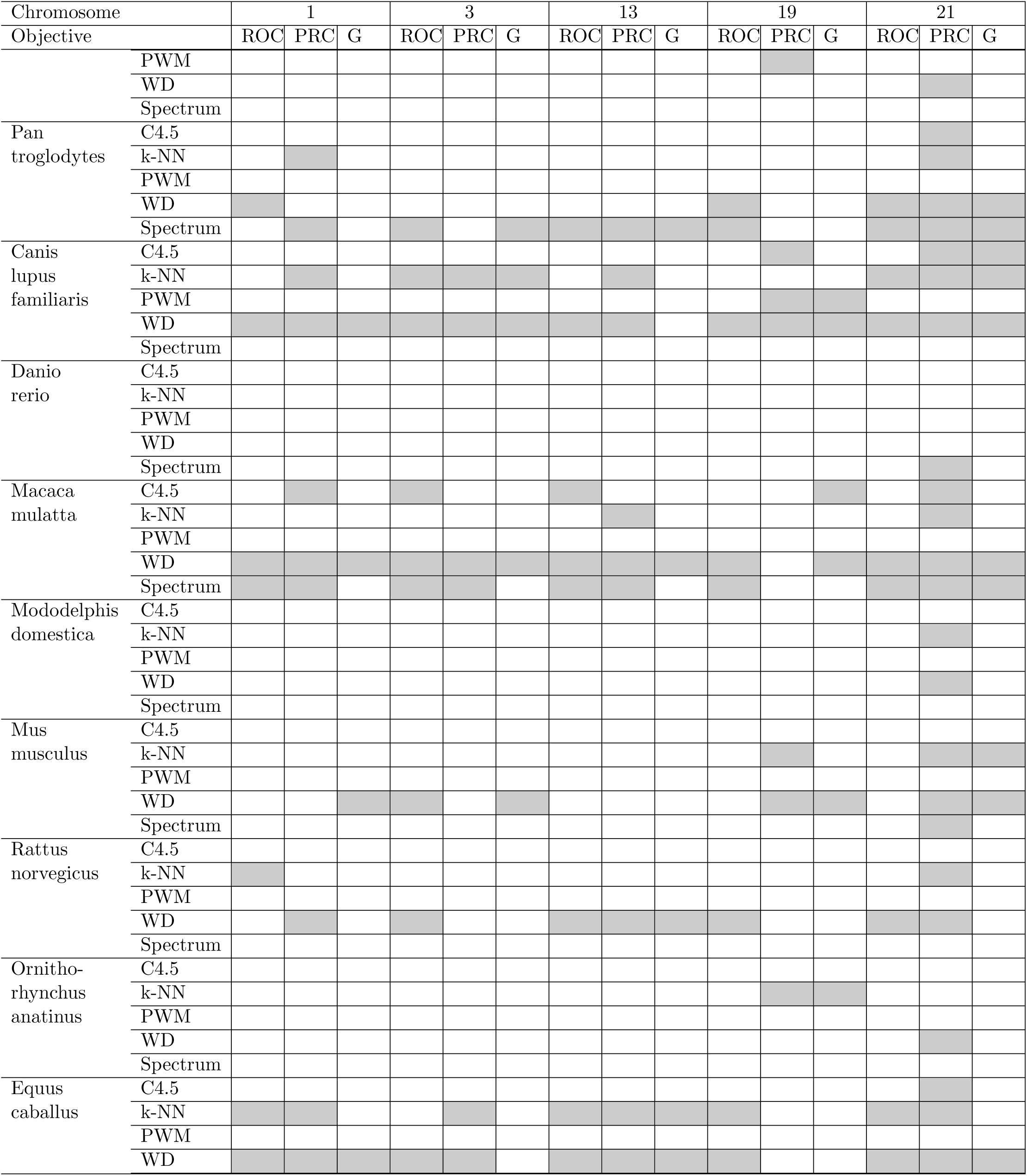
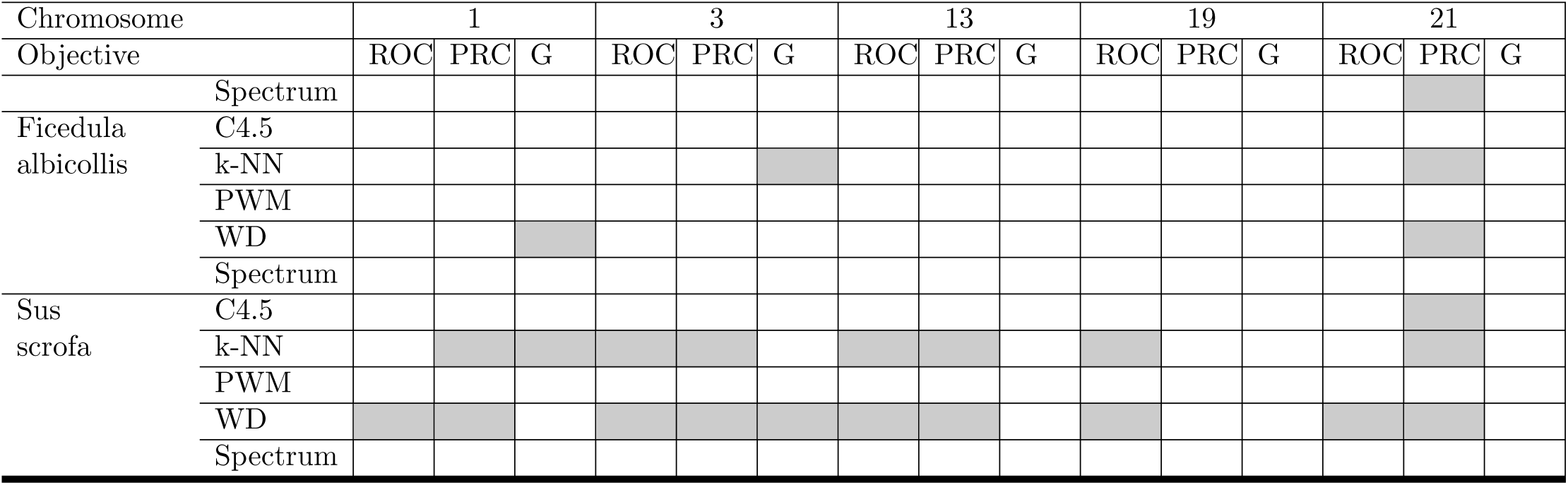
Models selected for acceptor site recognition.

The ROC and PRC curves for our approach and standard method are shown in Figs. 11 to 15. These results demonstrate that the overall performance of the proposed method was better than the performance of the best model.

**Figure 11.**
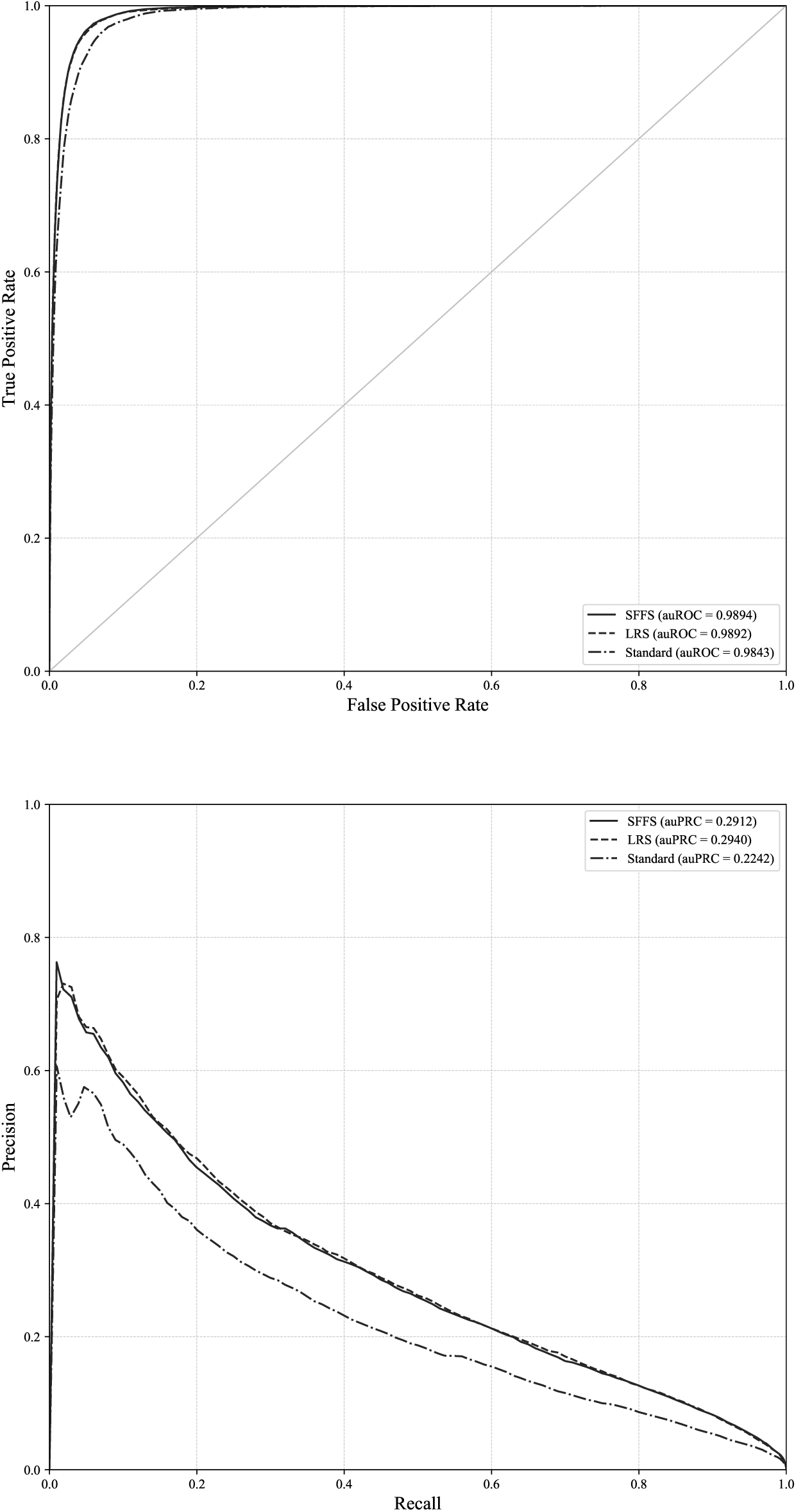
ROC and PRC curves for the acceptor site and chromosome 1. ROC and PRC curves for chromosome 1 and the standard approach and our proposed method when auROC and auPRC are optimized.

**Figure 12.**
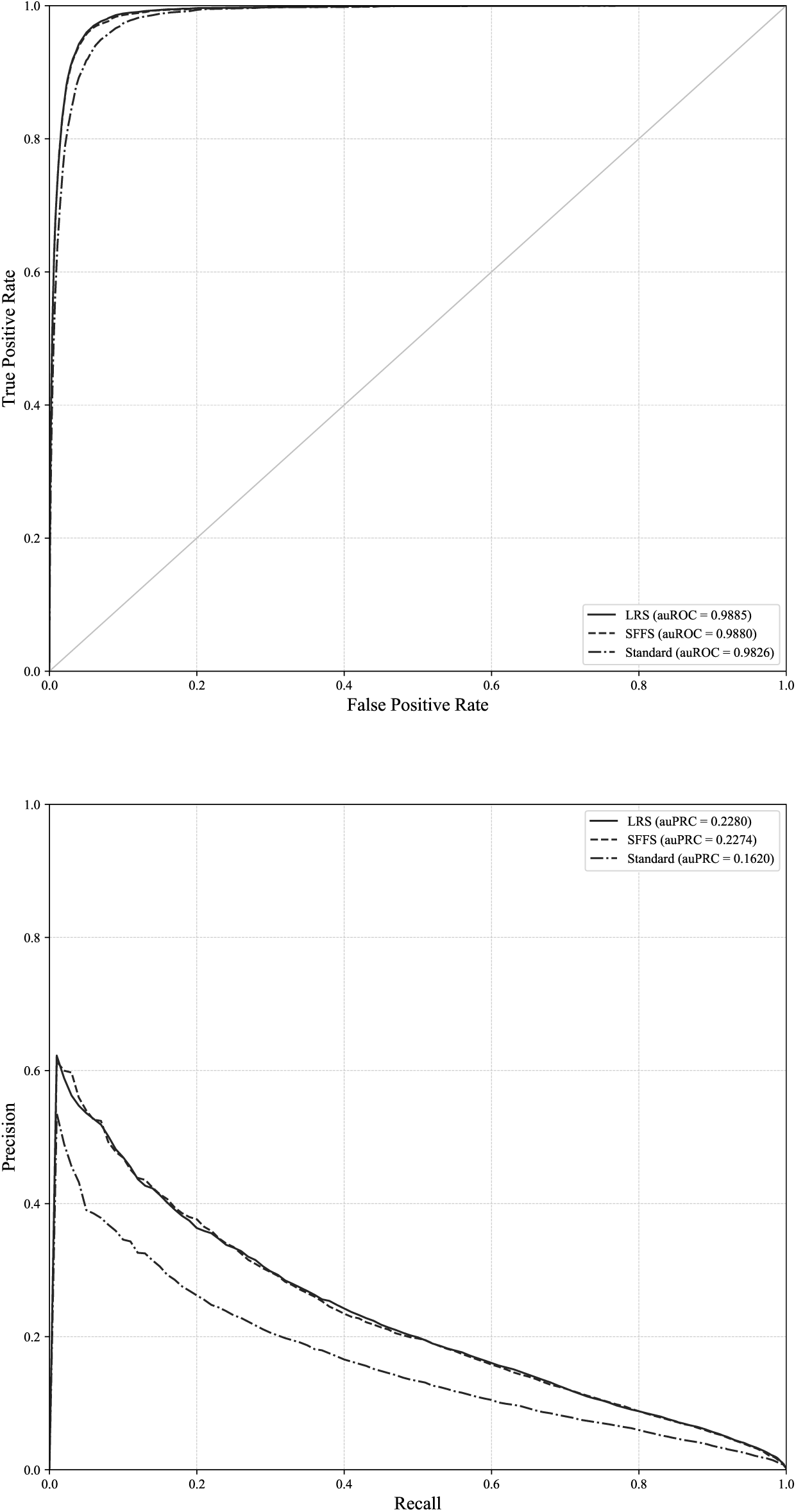
ROC and PRC curves for the acceptor site and chromosome 1. ROC and PRC curves for chromosome 3 and the standard approach and our proposed method when auROC and auPRC are optimized.

**Figure 13.**
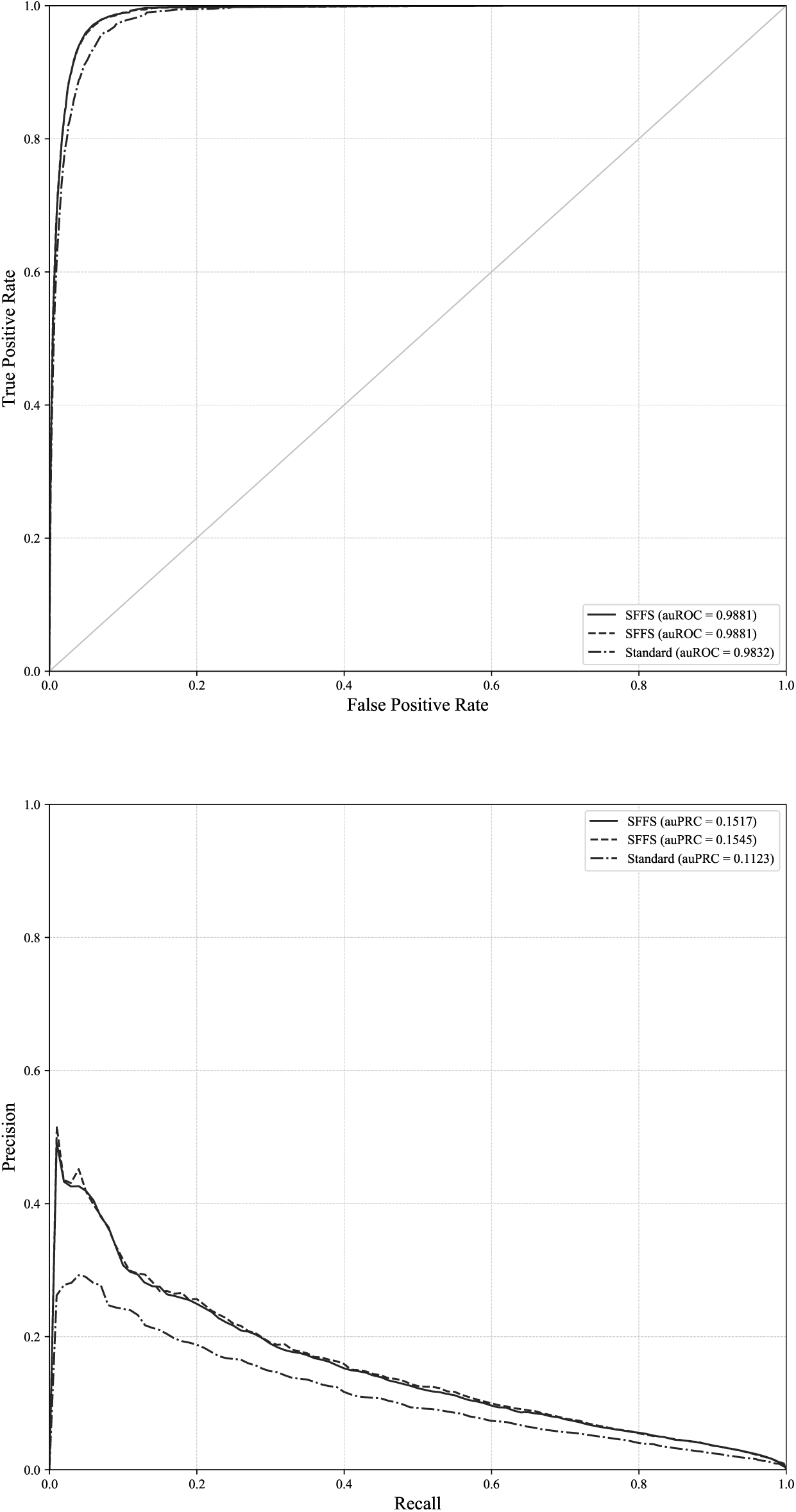
ROC and PRC curves for the acceptor site and chromosome 1. ROC and PRC curves for chromosome 13 and the standard approach and our proposed method when auROC and auPRC are optimized.

**Figure 14.**
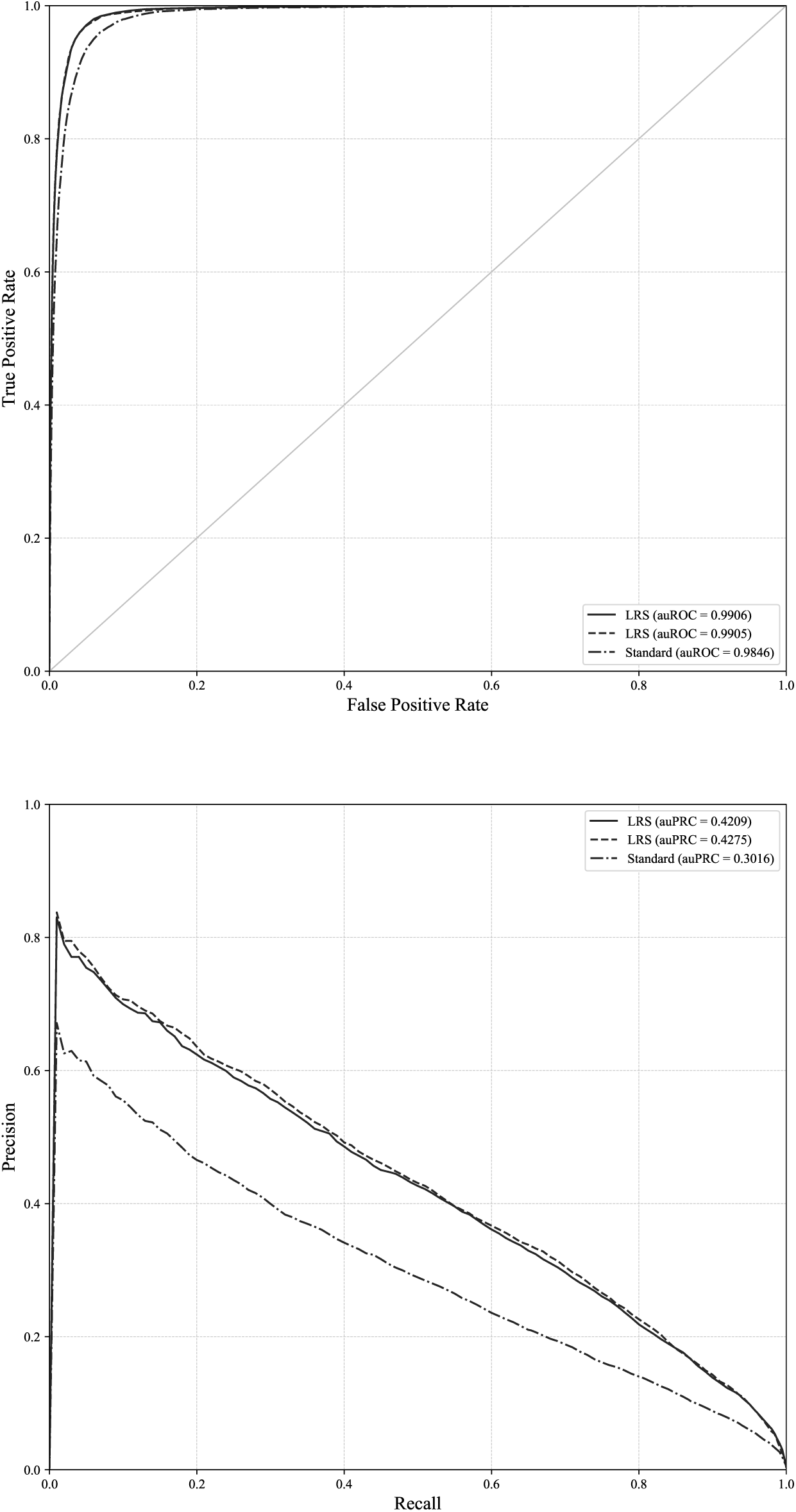
ROC and PRC curves for the acceptor site and chromosome 19. ROC and PRC curves for chromosome 19 and the standard approach and our proposed method when auROC and auPRC are optimized.

**Figure 15.**
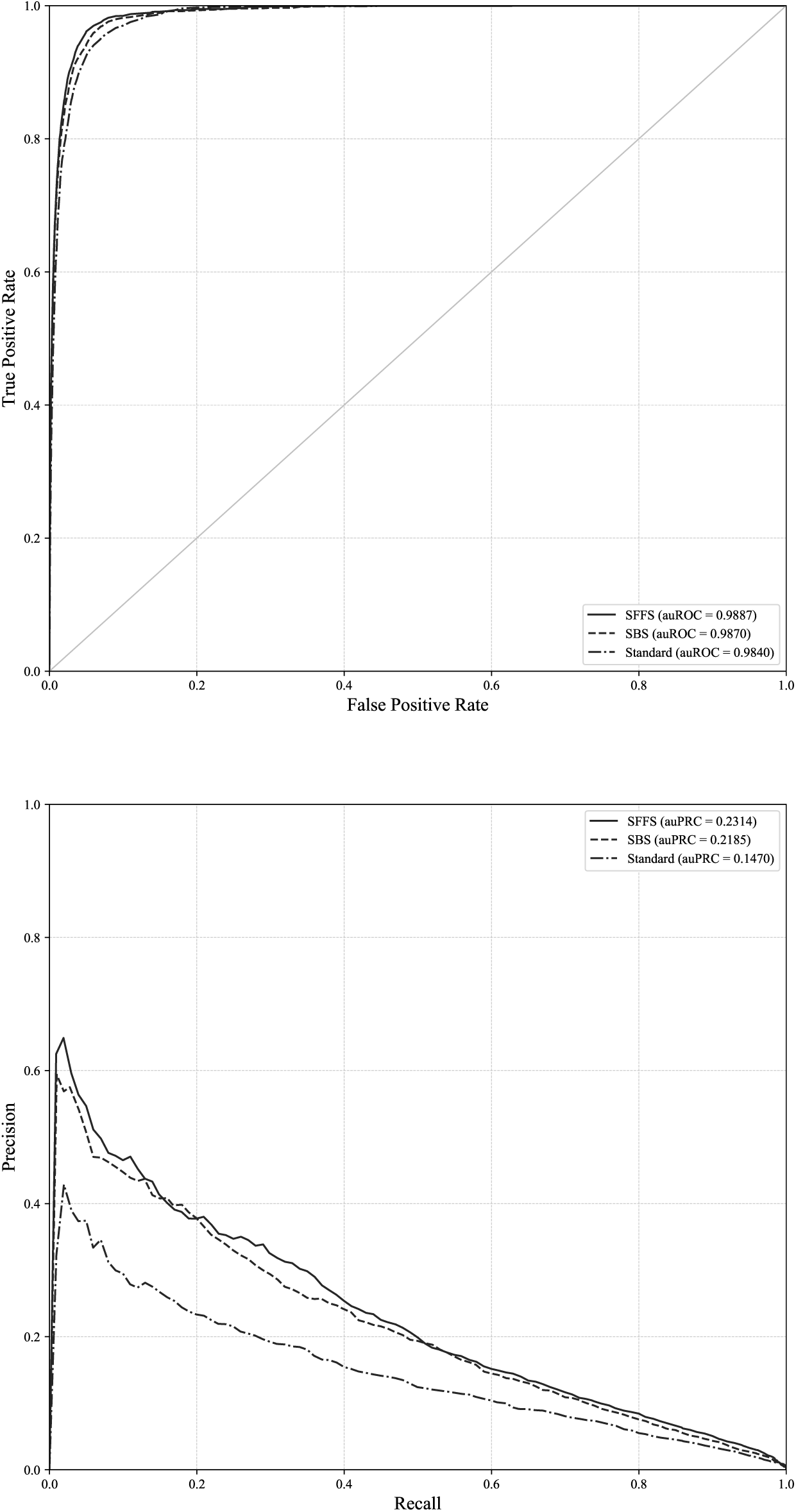
ROC and PRC curves for the acceptor site and chromosome 1. ROC and PRC curves for chromosome 21 and the standard approach and our proposed method when auROC and auPRC are optimized.

### Results for stop codon recognition

Stop codon recognition is the most difficult task of the four types of recognition. One of the major sources of this increased complexity is the number of negative instances, which means a much larger minority:majority ratio. The global majority:minority ratio for TISs is 1:4,155, for donors 1:326, for acceptors 1:455 and for stop codons 1:12,237. Table 8 shows the results for the five chromosomes and the three optimization measures.

**Table 8.** Results for stop codon recognition for human chromosomes 1, 3, 13, 19 and 21.

| Chrom. | Objective | Method | Combination | G | auROC | auPRC | TP | FN | TN | FP |
| --- | --- | --- | --- | --- | --- | --- | --- | --- | --- | --- |
|  | State-of-the-art |  |  | 0.8363 | 0.9233 | 0.0100 | 1,690 | 464 | 21,015,2752 | 2,557,756 |
| 1 | auROC | SFFS | Sum | 0.8769 | **0.9692** | 0.0285 | 1,704 | 450 | 22,912,5416 | 660,490 |
|  | auPRC | LRS | Sum | 0.8211 | 0.9567 | **0.0518** | 1,487 | 667 | 23,019,3613 | 553,670 |
|  | G | LRS | Sum | **0.9079** | 0.9705 | 0.0322 | 1,856 | 298 | 22,548,6526 | 1,024,379 |
|  | State-of-the-art |  |  | 0.8298 | 0.9177 | 0.0060 | 862 | 252 | 19,149,9512 | 2,372,549 |
| 3 | auROC | SFBS | Sum | 0.8758 | **0.9691** | 0.0314 | 875 | 239 | 21,019,7347 | 502,766 |
|  | auPRC | SBS | Sum | 0.8204 | 0.9568 | **0.0349** | 764 | 350 | 21,123,6862 | 398,814 |
|  | G | SFFS | Majority | **0.9097** | 0.9592 | 0.0028 | 979 | 135 | 20,268,8975 | 1,253,603 |
|  | State-of-the-art |  |  | 0.8120 | 0.9163 | 0.0031 | 242 | 91 | 9,870,9594 | 1,007,343 |
| 13 | auROC | SFFS | Sum | 0.8462 | **0.9598** | 0.0116 | 243 | 90 | 10,673,3721 | 204,930 |
|  | auPRC | SFBS | Sum | 0.7584 | 0.9497 | **0.0317** | 194 | 139 | 10,740,1530 | 138,149 |
|  | G | SFFS | Majority | **0.8961** | 0.9057 | 0.0004 | 287 | 46 | 10,134,5112 | 743,791 |
|  | State-of-the-art |  |  | 0.8426 | 0.9284 | 0.0295 | 1,141 | 281 | 4,128,2713 | 537,533 |
| 19 | auROC | SFFS | Sum | 0.9055 | **0.9697** | 0.0824 | 1,236 | 186 | 4,400,9924 | 264,812 |
|  | auPRC | RSFBS | Sum | 0.8818 | 0.9634 | **0.1167** | 1,176 | 246 | 4,386,4729 | 279,332 |
|  | G | SFFS | Majority | **0.9145** | 0.9630 | 0.0066 | 1,319 | 103 | 4,207,1045 | 458,700 |
|  | State-of-the-art |  |  | 0.7820 | 0.9017 | 0.0065 | 156 | 81 | 3,462,5527 | 264,407 |
| 21 | auROC | SFBS | Sum | 0.8487 | **0.9681** | 0.0254 | 175 | 62 | 3,635,8196 | 91,140 |
|  | auPRC | SBS | Sum | 0.7989 | 0.9600 | **0.0362** | 155 | 82 | 3,637,0015 | 89,958 |
|  | G | SFFS | Majority | **0.8994** | 0.9610 | 0.0025 | 205 | 32 | 3,485,0936 | 241,866 |

For auROC, the results showed a clear improvement. In the worst case, our approach improved the results by more than 4% and in the best case by more than 6%. These improvements were also achieved for the auPRC and *G-*mean measures. The usefulness of our approach can be corroborated by comparing the number of FPs between the standard method and our proposed method. Overall, for the five chromosomes, the standard approach obtained 6,739,588 FPs, whereas our method reduced that number to 1,459,923, which means that more than five million FPs were removed. The effect of that dramatic improvement on the recognition ability for stop codons must be significant over any gene structure prediction program.

As in previous results, the most common combination method was to sum the outputs, although majority voting was selected as a general rule for the *G-*mean measure, with the exception of chromosome 1. The searching strategies that obtained the best results depended on the experiment, which demonstrates the advantage of using all of them.

The numbers of models and selected classifiers and genomes for every case are shown in Table 9. *G-*mean, as in the previous results, was the measure that required fewer models, from 2 for chromosome 13 to 6 for chromosomes 1, 3 and 21. auROC selected from 7 to 15 models. Again, auPRC required a comparatively large number of models, from 31 to 58 selected models.

**Table 9.**
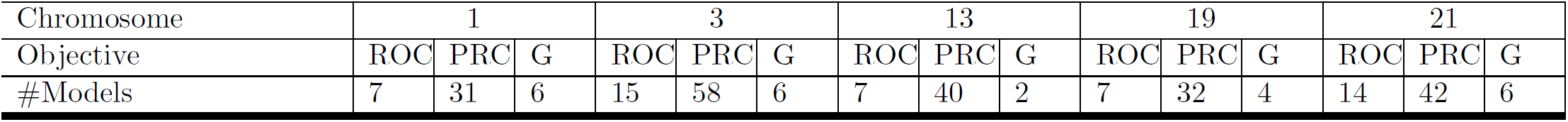
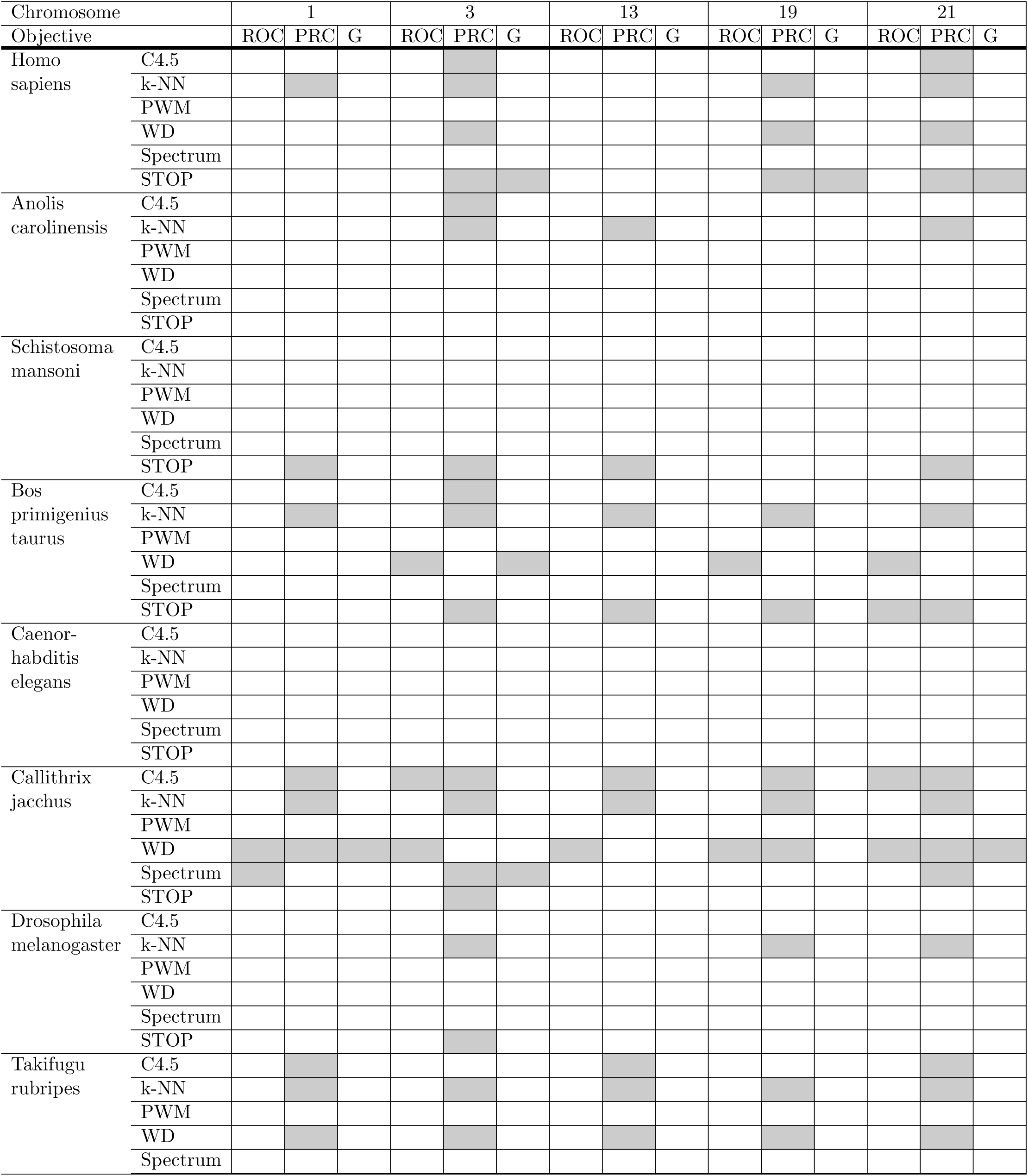
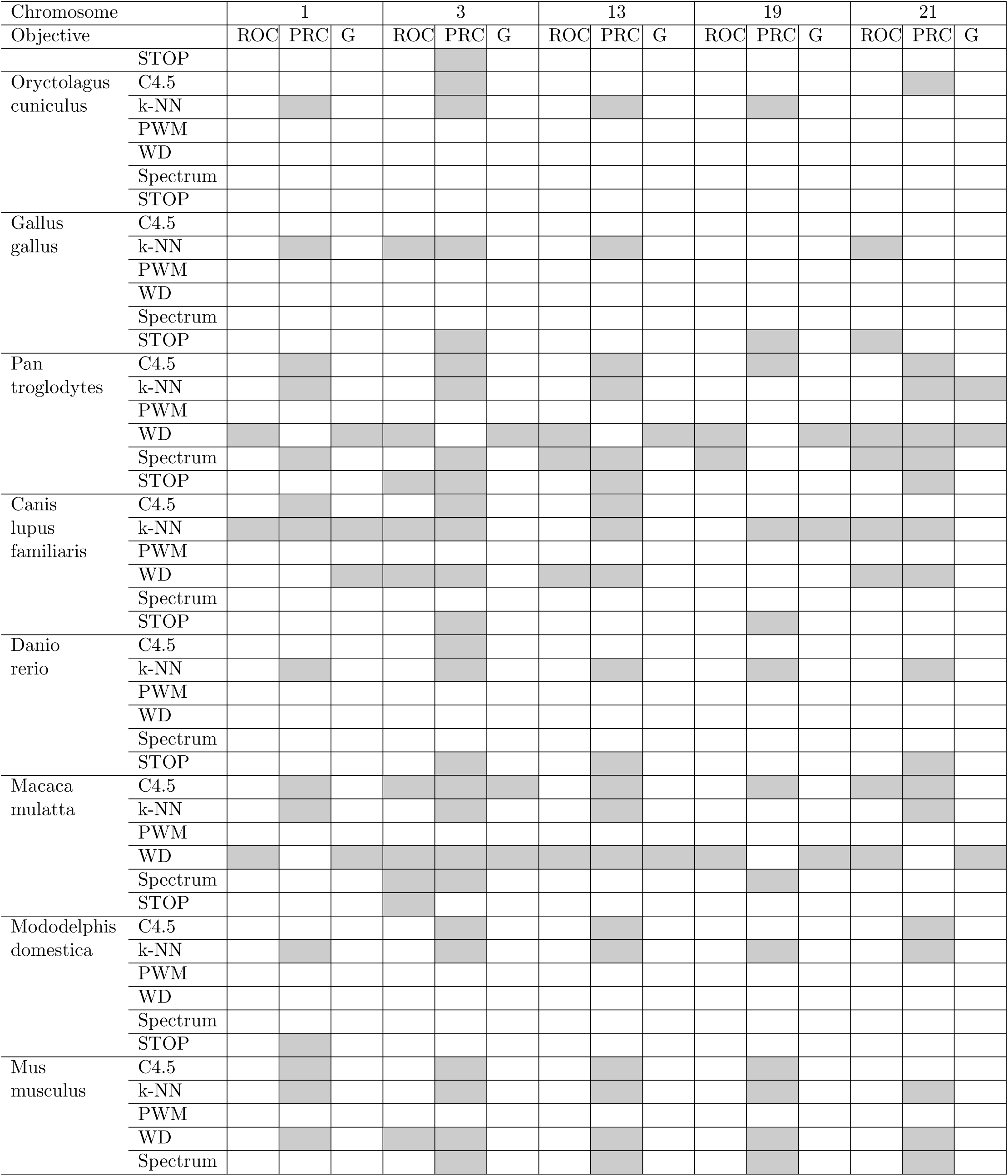
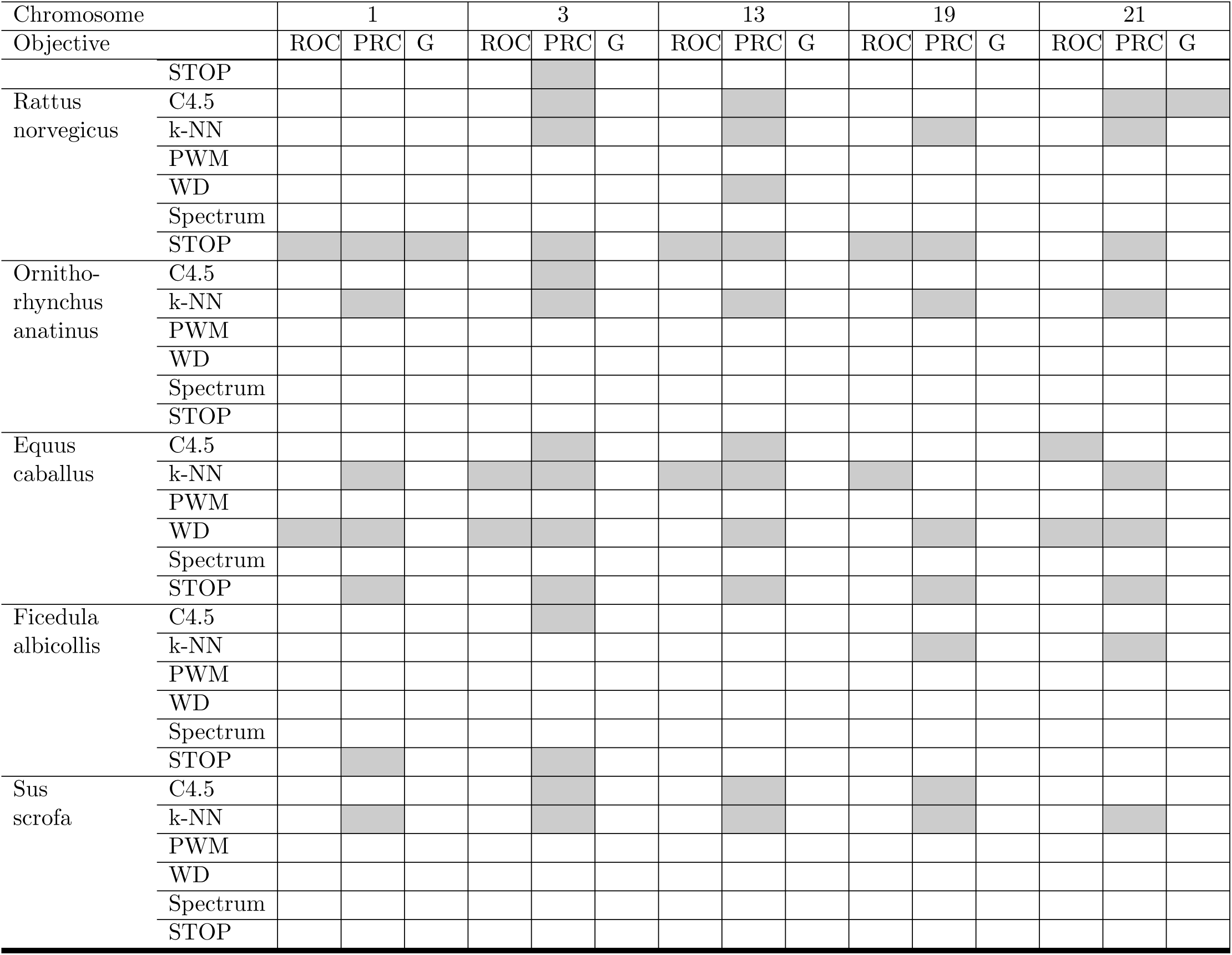
Models selected for stop codon recognition.

*Caenorhabditis elegans* was the only genome that was never used. The use of all the genomes was more balanced for the recognition of stop codons, using even genomes that were far removed from the human genome, such as those of *Takifugu rubripes* of *Danio rerio*. The use of classifiers was also more equally distributed among the six methods, with the exception of PWM, which was never used.

With respect to the three objectives, optimizing the *G-*mean required fewer models, from 2 to 6. For the five chromosomes, the SVM method for *Macaca mulatta* and *Pan troglodytes* was always selected. *Callithrix jacchus* and *Canis lupus familiaris* were also selected in most chromosomes. For auROC, more models were selected, from 7 to 15. The SVM method for *Macaca mulatta* and *Pan troglodytes* was always chosen, but the remaining methods depended on the chromosome. This is another interesting result because most stop codon recognition programs rely on common models for any task. Finally, for auPRC, significantly more models were selected, from 31 to 58, with a significant variation among the chromosomes.

The actual ROC and PRC curves, which are shown in Figs. 16–20, show that the curves that correspond to our proposed method are always above the curves of the best model. This indicates better performance for all the possible thresholds of classification.

**Figure 16.**
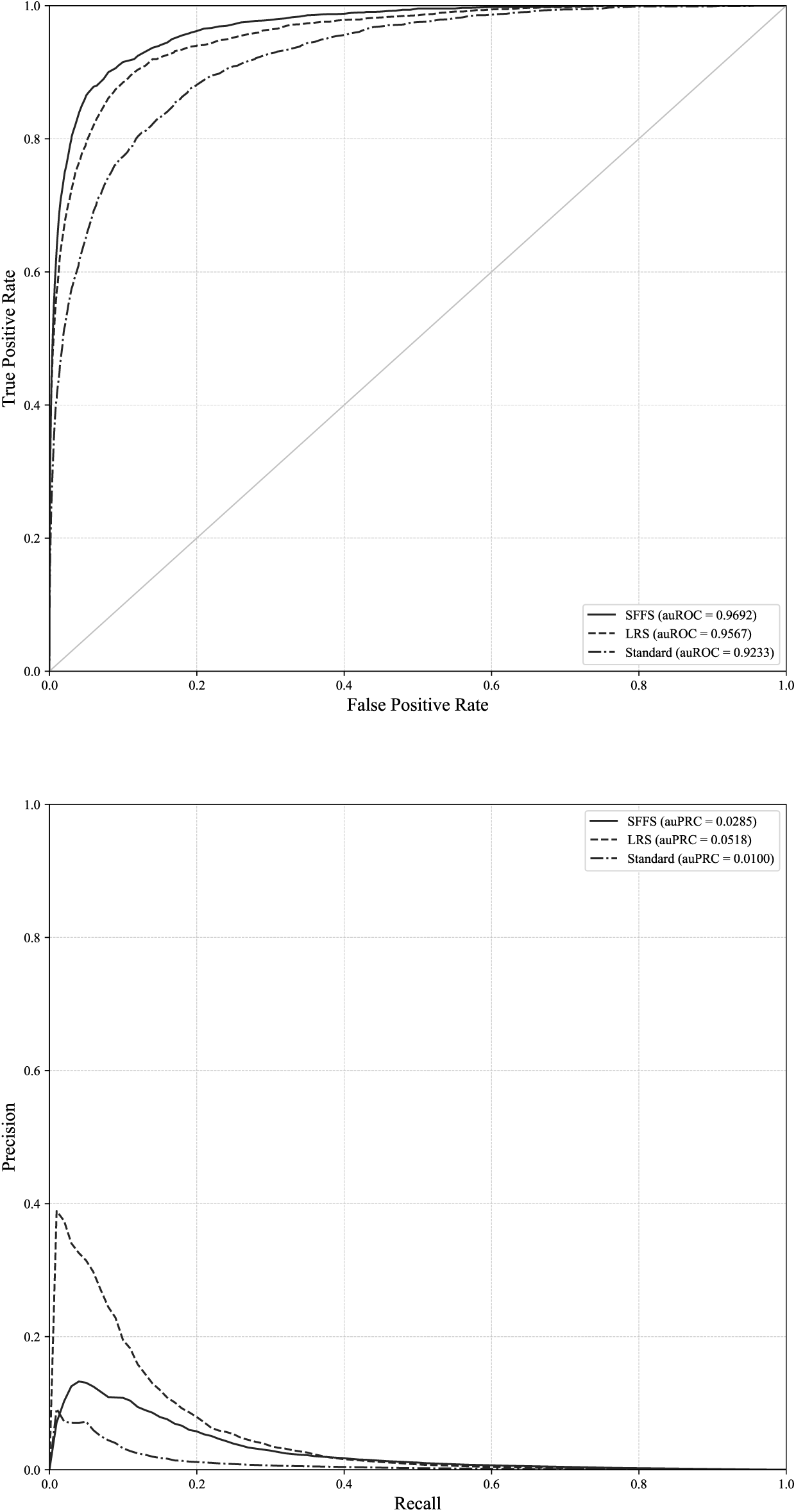
ROC and PRC curves for the STOP codon and chromosome 1. ROC and PRC curves for chromosome 1 and the standard approach and our proposed method when auROC and auPRC are optimized.

**Figure 17.**
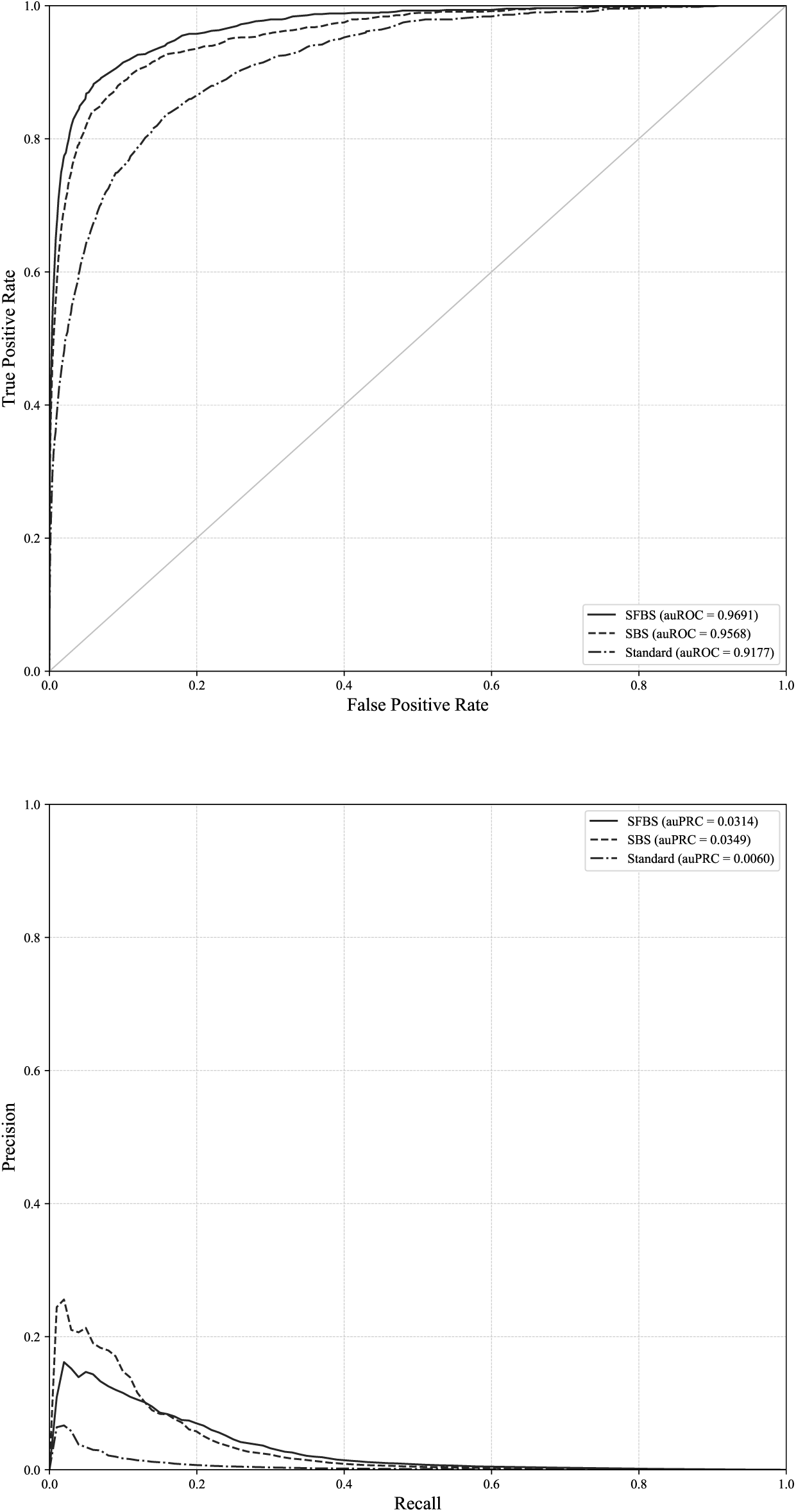
ROC and PRC curves for the STOP codon and chromosome 1. ROC and PRC curves for chromosome 3 and the standard approach and our proposed method when auROC and auPRC are optimized.

**Figure 18.**
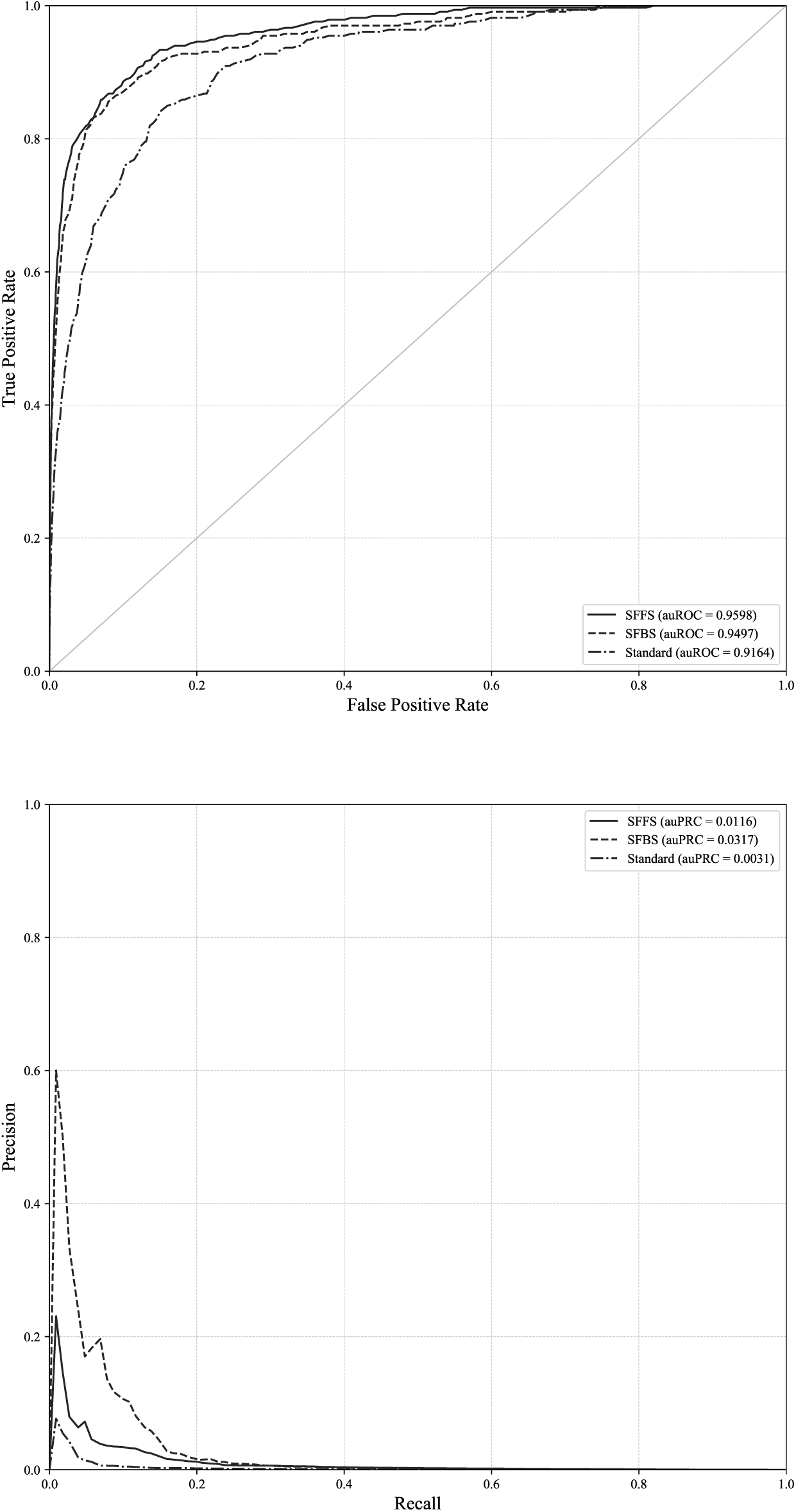
ROC and PRC curves for the STOP codon and chromosome 1. ROC and PRC curves for chromosome 13 and the standard approach and our proposed method when auROC and auPRC are optimized.

**Figure 19.**
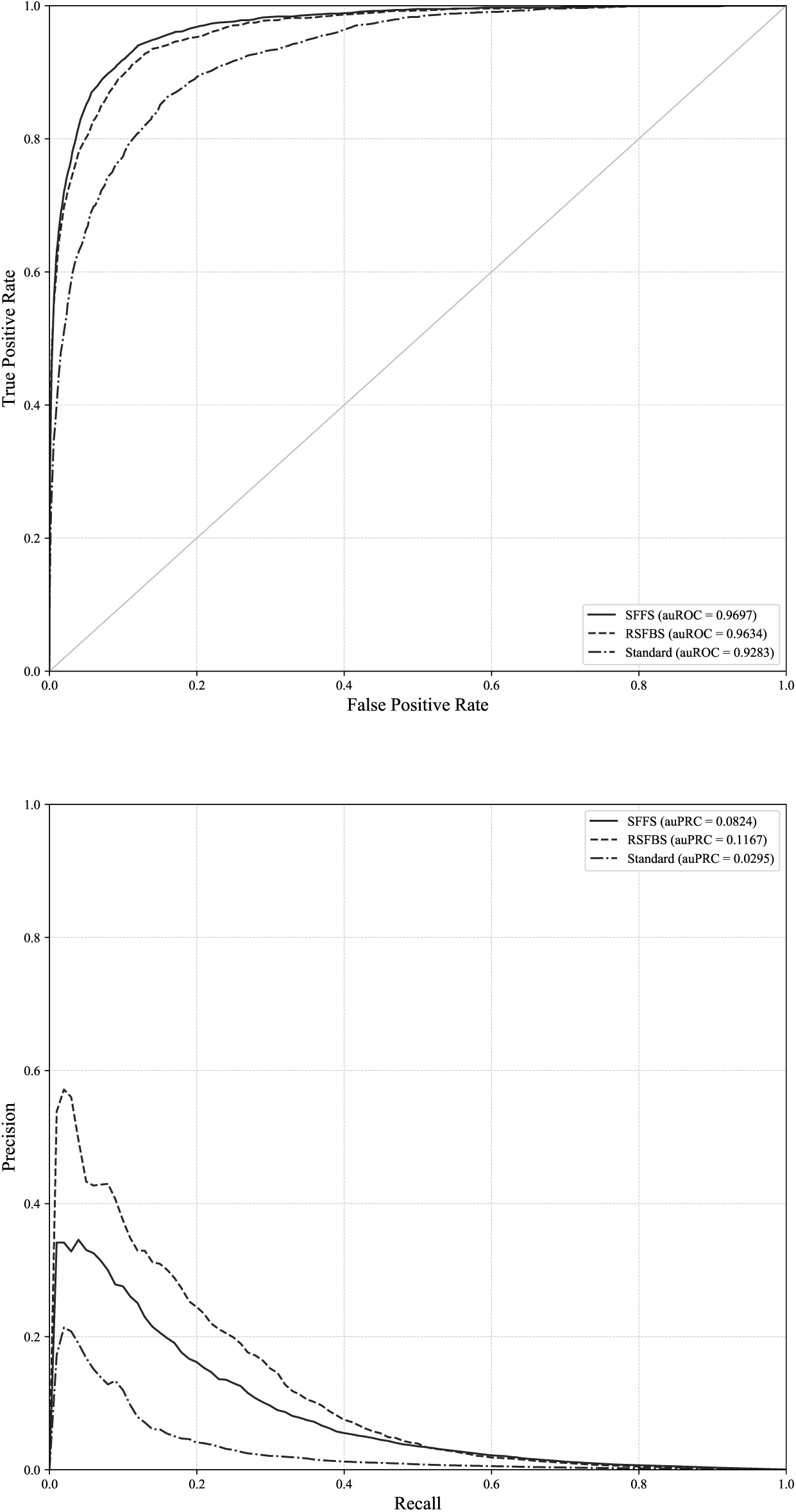
ROC and PRC curves for the STOP codon and chromosome 19. ROC and PRC curves for chromosome 19 and the standard approach and our proposed method when auROC and auPRC are optimized.

**Figure 20.**
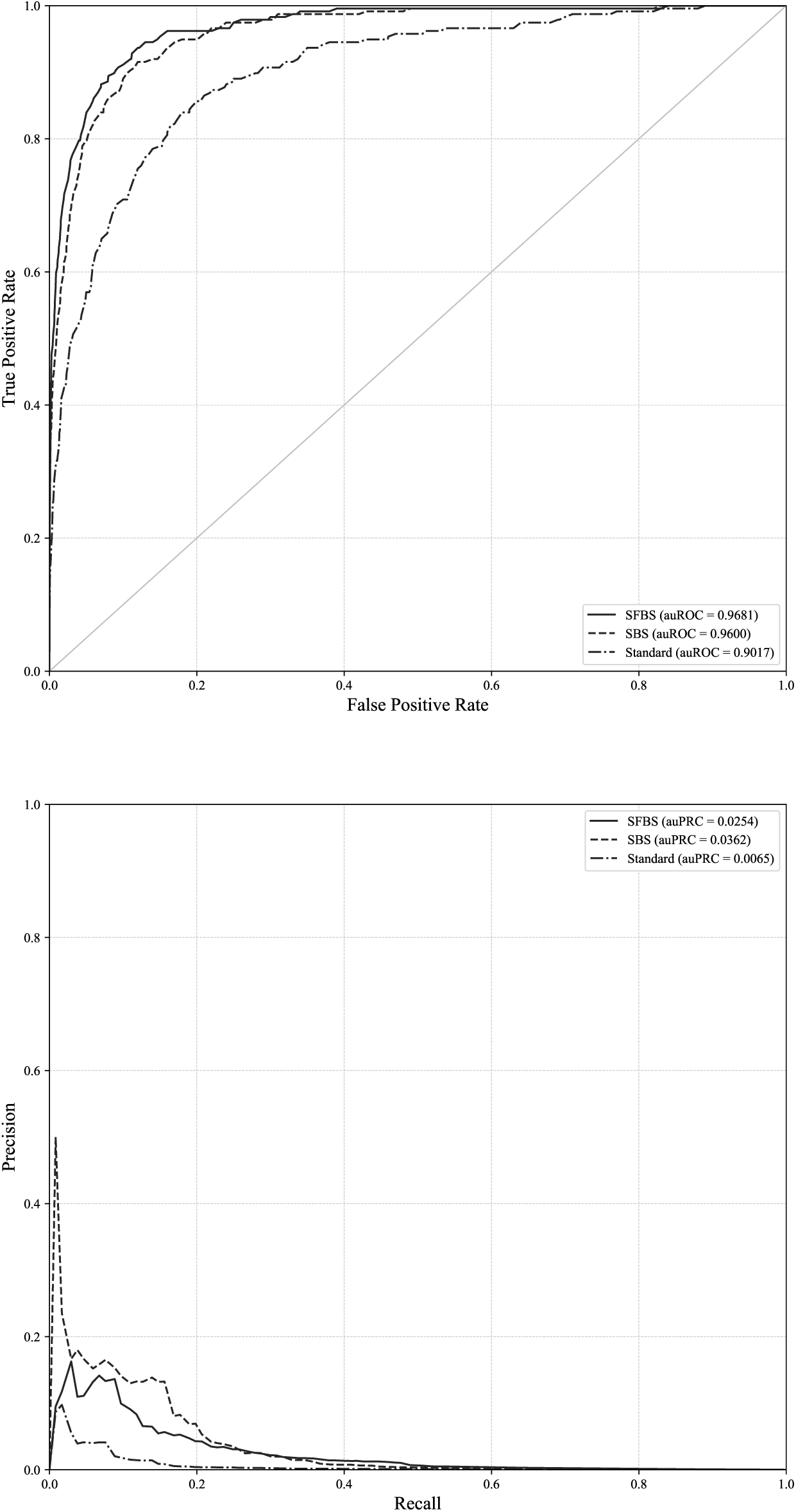
ROC and PRC curves for the STOP codon and chromosome 1. ROC and PRC curves for chromosome 21 and the standard approach and our proposed method when auROC and auPRC are optimized.

## Summary of the comparison

As a summary of the comparison for the five chromosomes and four sites, Figs. 21, 22, 23, 24 and 25 show the improvements for all four sites and five chromosomes. The figures show the relative improvement of our approach in terms of *G-*mean, auROC and auPRC. All of the figures show the improvement that obtained using the floating search strategy.

**Figure 21.**
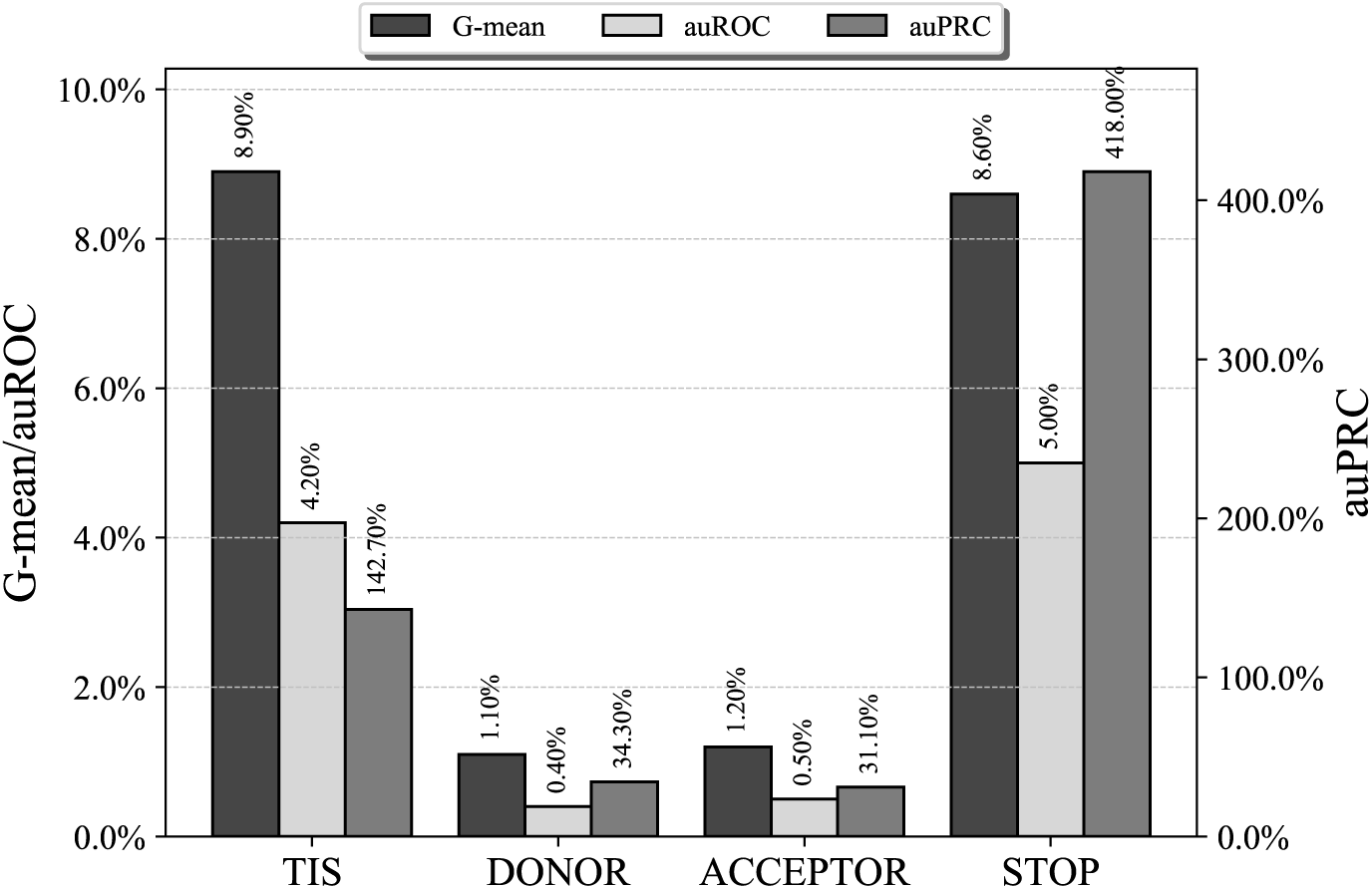
Chromosome 1 result comparison. Relative improvement in the chromosome 1 results of our method against a state-of-the-art standard method.

**Figure 22.**
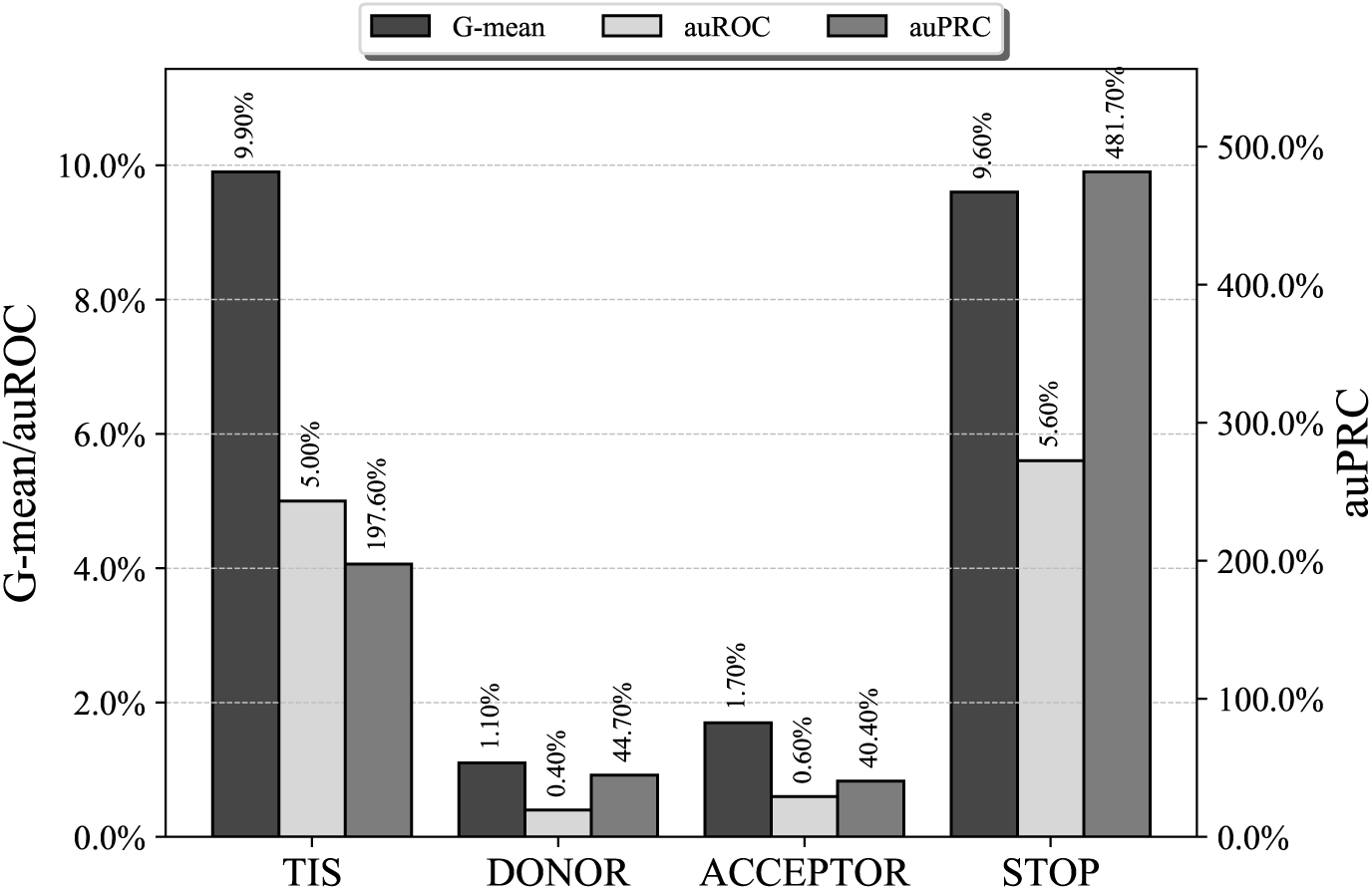
Chromosome 3 result comparison. Relative improvement in the chromosome 3 results of our method against a state-of-the-art standard method.

**Figure 23.**
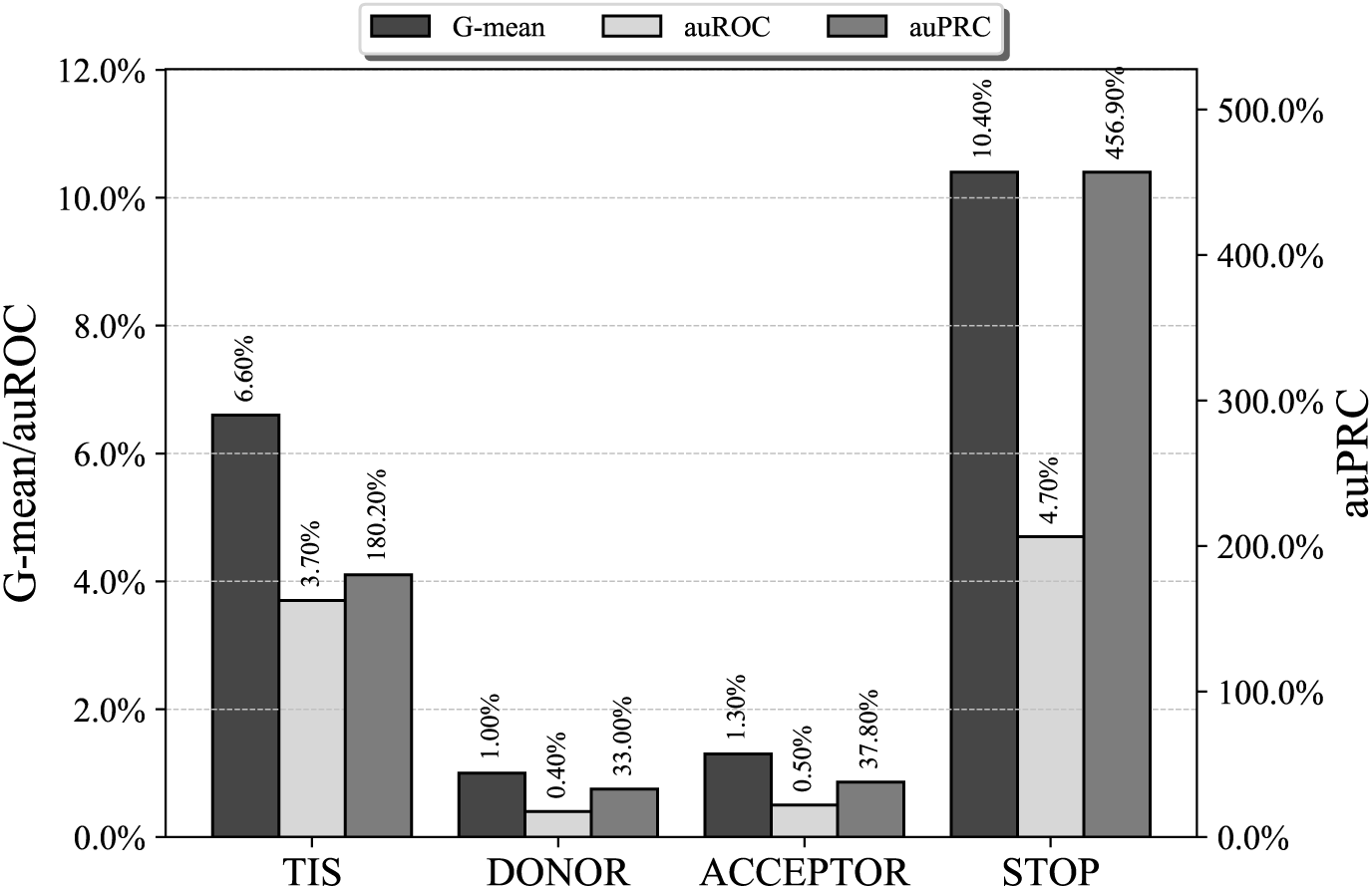
Chromosome 13 result comparison. Relative improvement in the chromosome 13 results of our method against a state-of-the-art standard method.

**Figure 24.**
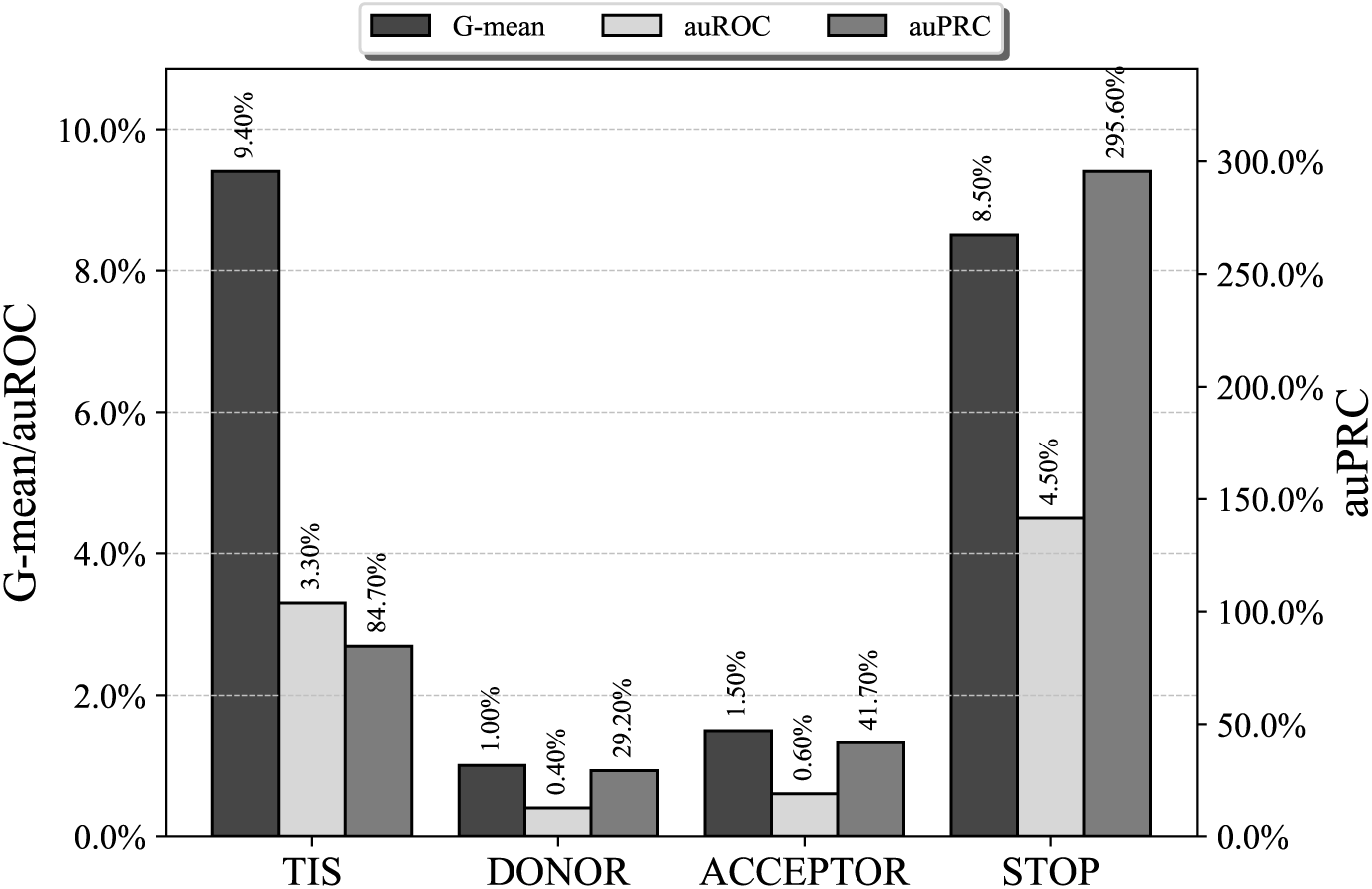
Chromosome 19 result comparison. Relative improvement in the chromosome 19 results of our method against a state-of-the-art standard method.

**Figure 25.**
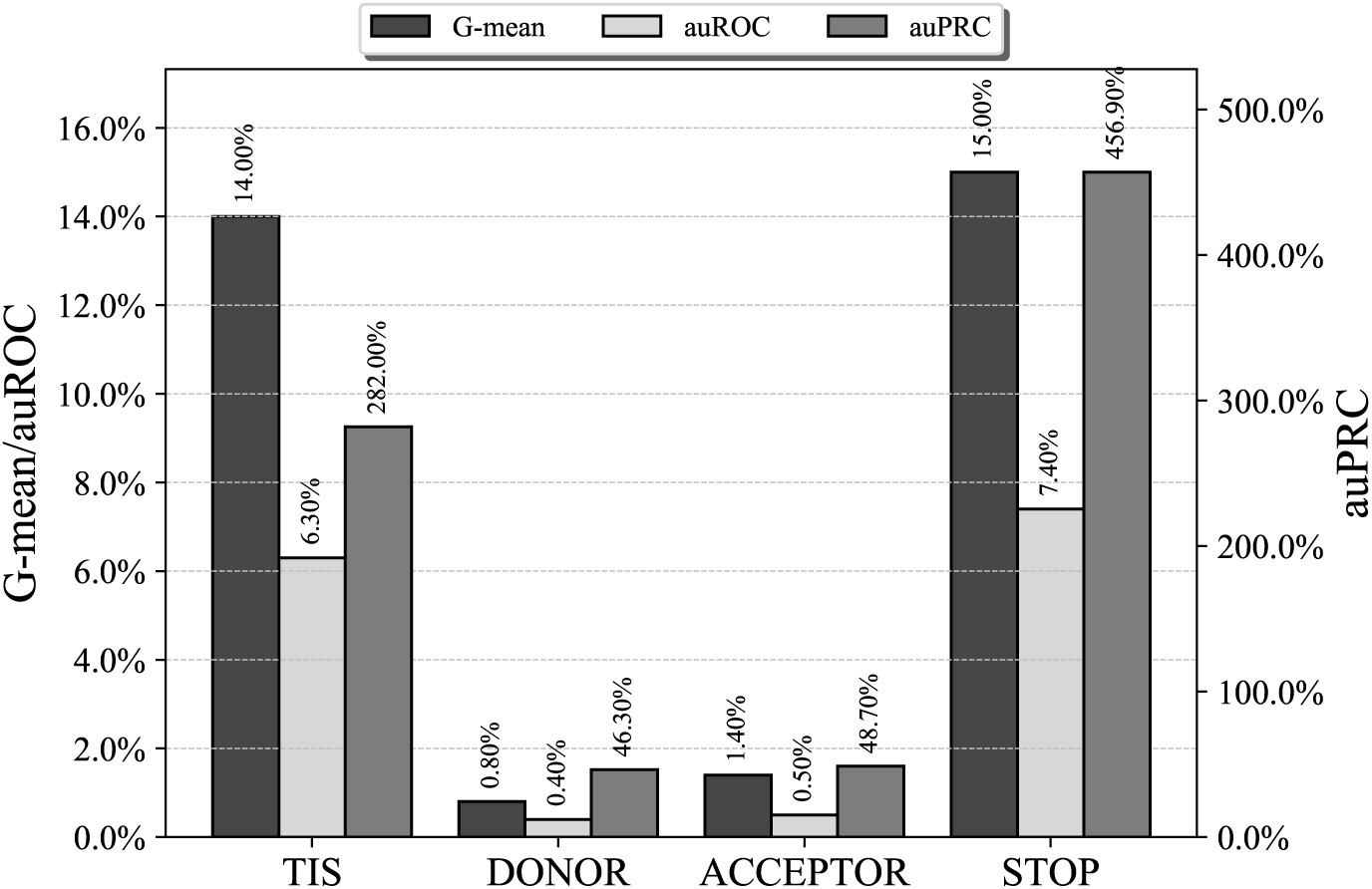
Chromosome 21 result comparison. Relative improvement in the chromosome 21 results of our method against a state-of-the-art standard method.

Finally, to study the effect on the performance of the proposed method of the optimization measure, we show in Figs. 26, 27 and 28 the overall improvements in terms of TPs, FNs, TNs and FNs for all of the chromosomes and the four sites. The first conclusion is that the optimization objective has a relevant impact on the distribution of the errors. That is a very important aspect if we plan to use site recognition as an initial step in a gene structure prediction task, as our prediction program might be more sensitive to a specific type of errors.

**Figure 26.**
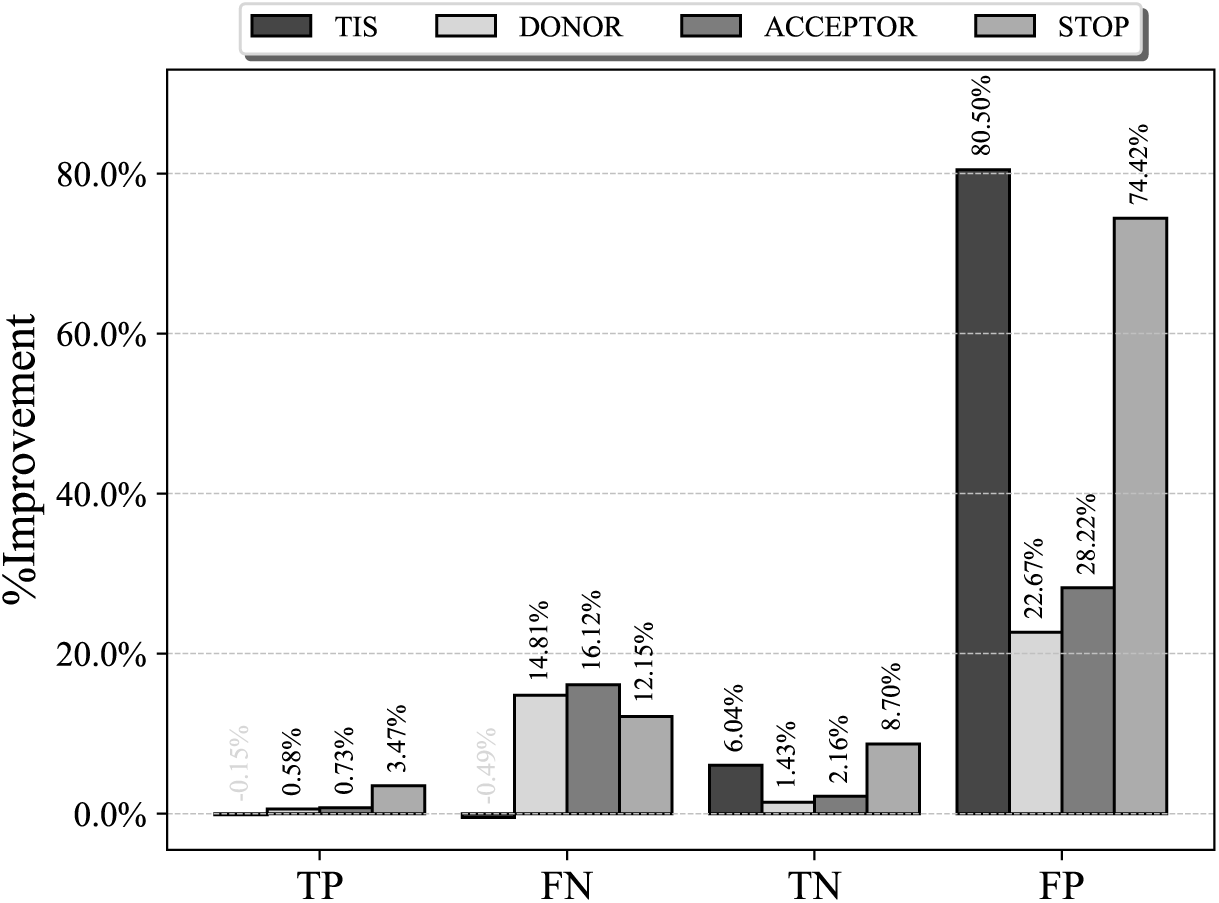
TP, FN, TN, and FP improvements for all four sites using auROC as the optimization objective. Overall improvement results of our method against a state-of-the-art standard method.

**Figure 27.**
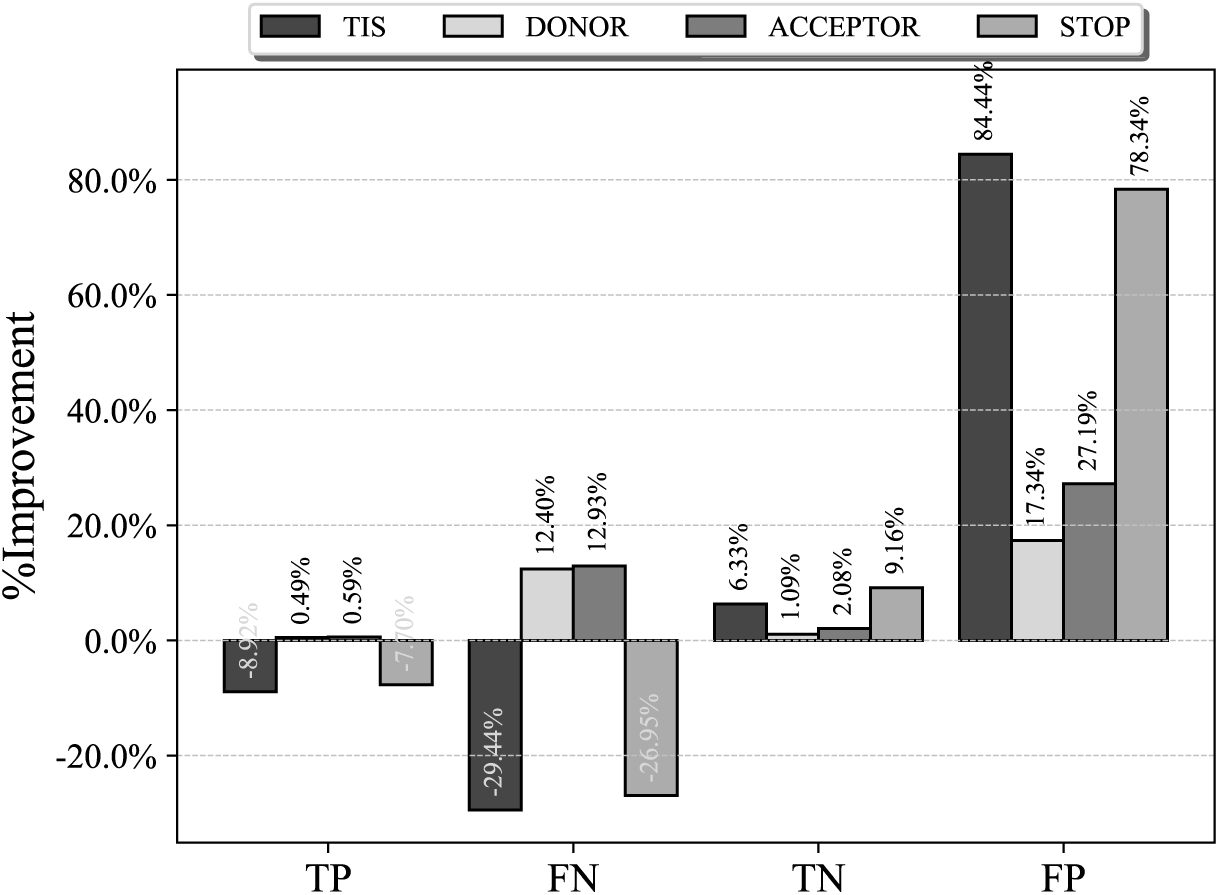
TP, FN, TN, and FP improvements for all four sites using auPRC as the optimization objective. Overall improvement results of our method against a state-of-the-art standard method.

**Figure 28.**
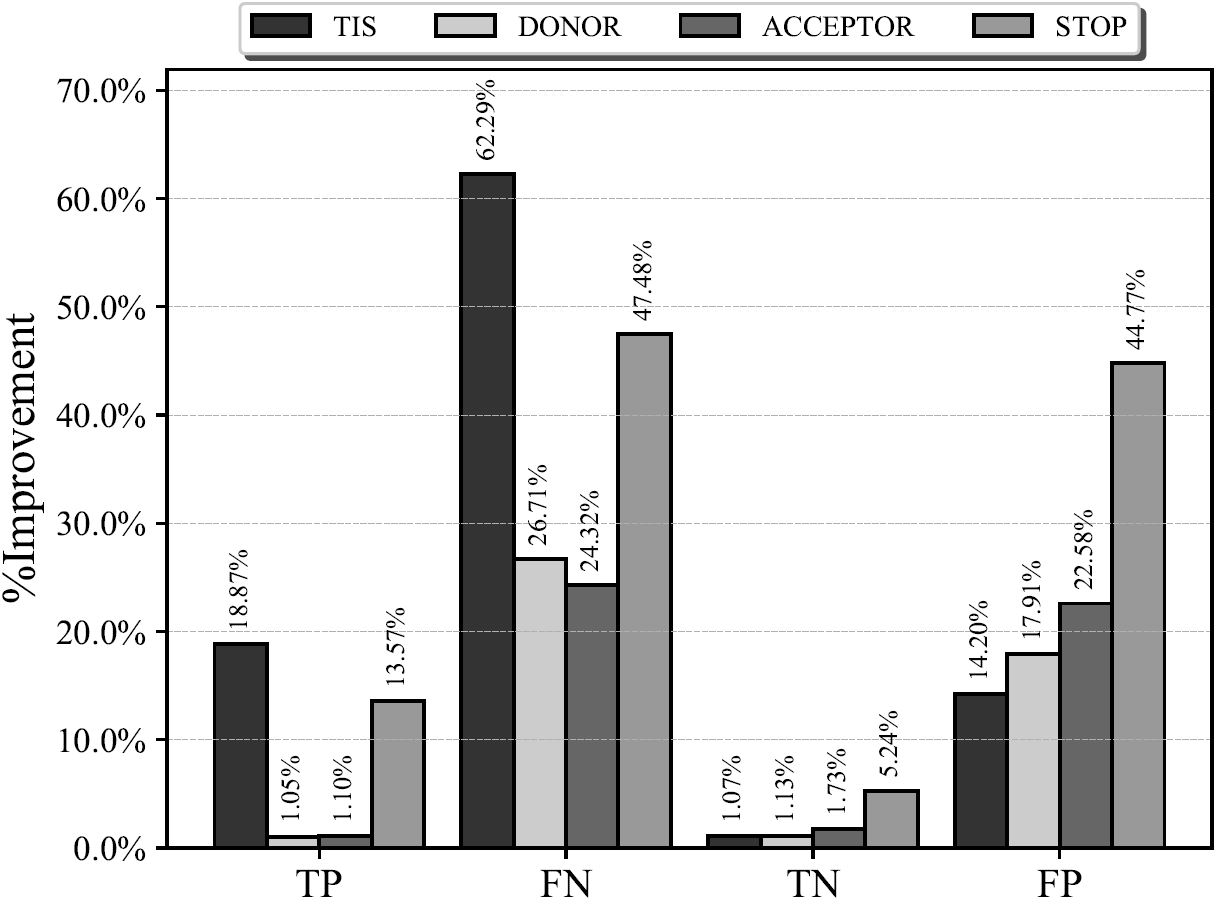
TP, FN, TN, and FP improvements for all four sites using *G-*mean as the optimization objective. Overall improvement results of our method against a state-of-the-art standard method.

For positive site prediction, the best results were obtained using *G-*mean as theobjective, whereas auROC only showed a minor improvement and auPRC was even worse than the standard approach for TIS and stop codon prediction. For negative instances, auROC and auPRC performed very well, with a small advantage of auPRC for TISs and stop codons and of auROC for donors and acceptors. *G-*mean achieved a marked improvement over the standard method but not as dramatic as those of auROC and auPRC. The best overall performance on both positive and negative samples was achieved by *G-*mean, which showed a more balanced behavior.

### Effect on gene prediction

We stated in the introduction that the improvement of the site prediction that was introduced by our method would have a significant impact on the prediction of the complete structure of genes, as site prediction is a relevant step in most current gene structure prediction programs. To test that statement, we performed a final experiment on gene prediction for chromosome 21. We constructed a very simple predictor that searched for exons using the sites that were found by the recognition program, which was either the standard approach or our proposed method, and constructed a gene using these exons. This simple program is not intended for gene structure prediction but only to test the ability of our proposed method in improving gene recognition.

To evaluate gene predictor performance over a test sequence, the predicted gene structure is compared with the annotated gene structure on the target sequence. The accuracy is evaluated at different levels of resolution. Commonly, these levels are the nucleotide, exon and gene levels. Due to the use of a very simple program, no gene-level accuracy is reported. Regarding nucleotide-level performance, we used as comparison measures Sensitivity (Sn):

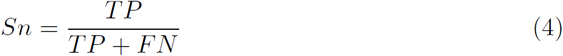

which is a relevant measure if we are interested only in the performance on the positive class, and Specificity (*Sp*), in its traditional machine learning form, which is defined as:

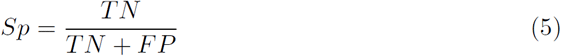

In bioinformatics, and particularly in gene prediction, specificity is usually defined in a different way. Specificity (*Sp*(*BIO*)) is calculated by dividing the number of correct predictions by the total number of predictions:

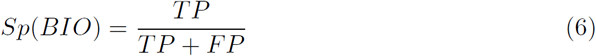

However, neither sensitivity nor specificity by itself constitutes a measure of global accuracy. A good measure that summarizes both at the nucleotide level is the Correlation Coefficient (CC):

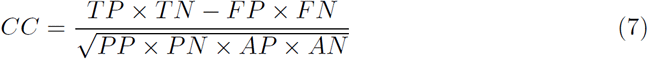

where *PP* is the number of predicted positives, *AP* the actual positives, *PN* the predicted negatives and *AN* the actual negatives. We also calculate the Average Conditional Probability (ACP) measure:

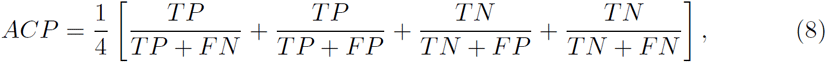

and the Approximate Correlation (AC):

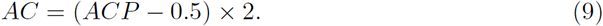

At the exon level, an exon is considered to have been correctly predicted when both boundaries are correctly predicted. If a predicted exon contains at least one actual base, it will be considered a partially correct exon. At the exon level, we show Sp, Sn and the numbers of missed exons (ME), which are exons that are not found by the program, and wrong exons (WE), which are predicted exons that do not correspond to any actual exon. As a representative of our proposed method, we used the model that was obtained when optimizing *G-*mean, as the previous section showed that it achieved the best overall behavior.

Fig 29 shows the performances of our proposed method and the standard method for the ten measures that are presented above. Fig 30 shows the relative improvementof our method with respect to the standard approach. Our approach improved the results of the standard method in terms of all measures. At the nucleotide level, CC and AC were improved by over 100%. At the exon level, Sn and Sp were improved significantly, while the numbers of ME and WE were also improved, but only marginally. These results show how our approach can be used to improve gene structure prediction.

**Figure 29.**
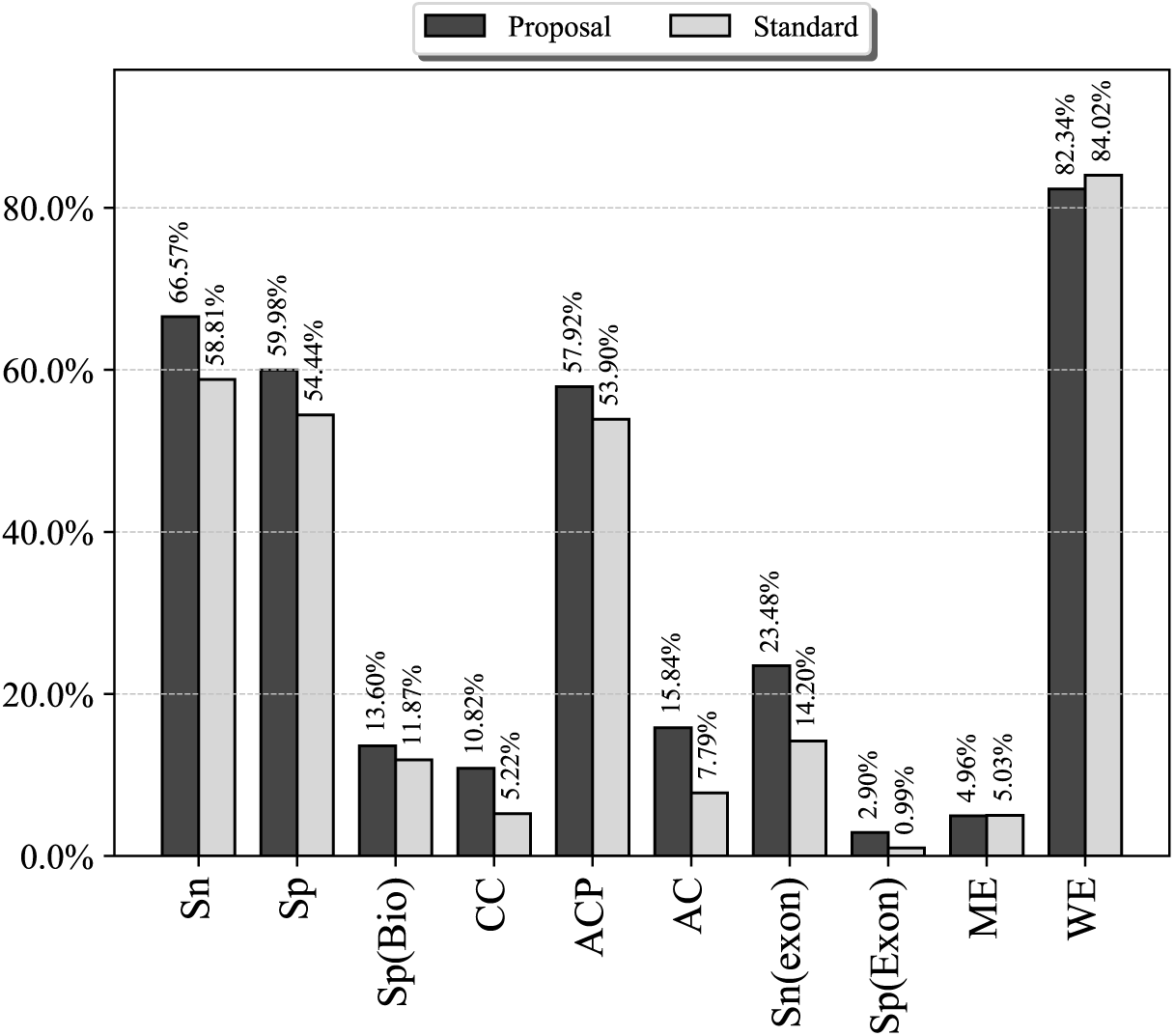
Chromosome 21 gene structure prediction. Chromosome 21 results of our method against a state-of-the-art standard method.

**Figure 30.**
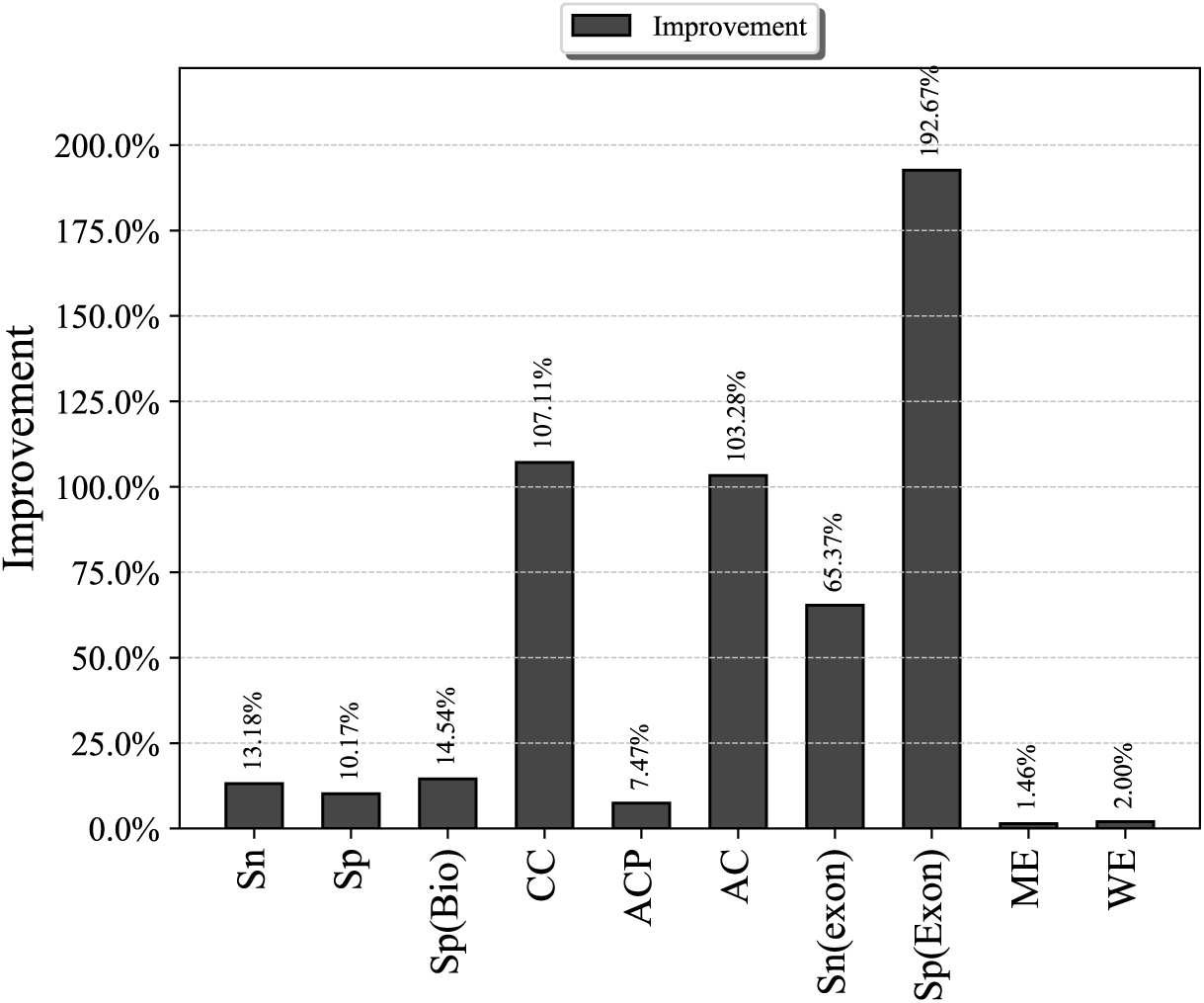
Chromosome 21 gene structure prediction. Relative improvement for chromosome 21 results of our method against a state-of-the-art standard method.

## Conclusions

In this paper, we presented a floating-search-based strategy for functional site recognition in genomic sequences. The use of floating search enables an efficient search for the best combination of more than a hundred of classification models that are trained on the genomes of many species. The presented approach can also be used for other combination tasks.

The proposed method also enabled the optimization of various performance measures. In the reported experiments, we showed results on searching for the best combination that optimizes three measures: auROC, auPRC and *G-*mean. The method was successfully applied to the recognition of TIS, donor and acceptor sites and stop codons. The reported experiments showed a clear improvement over the current best methods. The reported results also showed that to obtain the best classification rates, many species should be used. Our approach efficiently improved the performance of a very simple program for gene structure prediction.

## Acknowledgments

This work has been financed in part by Project TIN-2011-22967 of the Spanish Ministry of Science and Innovation and Excellence in Research Projects P09-TIC-4623 and P07-TIC-2682 of the Junta de Andalucía.

## Footnotes

1 In our experiment, this subset was obtained selecting each classifier with a probability of 0.5.

2 We always tested all the methods with all the negative samples, which means that the ratio of the minority/majority class was more than 1:3200 for the worst case, namely, stop codon recognition for chromosome 13 (see Table 1), which yielded low auPRC values. We must take into account that with only a few thousand FPs among several million TNs, we obtain a very low precision value. The situation for stop codon recognition is even worse, as the number of TNs is multiplied by three.

